# Long-time analytic approximation of large stochastic oscillators: simulation, analysis and inference

**DOI:** 10.1101/068148

**Authors:** Giorgos Minas, David A Rand

**Author notes:** **Abbreviations:** pcLNA, phase-corrected Linear Noise Approximation; KS, Kolmogorov-Smirnov; FIM, Fisher Information Matrix.

## Abstract

In order to analyse large complex stochastic dynamical models such as those studied in systems biology there is currently a great need for both analytical tools and also algorithms for accurate and fast simulation and estimation. We present a new stochastic approximation of biological oscillators that addresses these needs. Our method, called phase-corrected LNA (pcLNA) overcomes the main limitations of the standard Linear Noise Approximation (LNA) to remain uniformly accurate for long times, still maintaining the speed and analytically tractability of the LNA. As part of this, we develop analytical expressions for key probability distributions and associated quantities, such as the Fisher Information Matrix and Kullback-Leibler divergence and we introduce a new approach to system-global sensitivity analysis. We also present algorithms for statistical inference and for long-term simulation of oscillating systems that are shown to be as accurate but much faster than leaping algorithms and algorithms for integration of diffusion equations. Stochastic versions of published models of the circadian clock and NF-_*κ*_B system are used to illustrate our results.

## 1 Introduction

Dynamic cellular oscillating systems such as the cell cycle, circadian clock and other signaling and regulatory systems have complex structures, highly nonlinear dynamics and are subject to both intrinsic and extrinsic stochasticity. Moreover, current models of these systems have high-dimensional phase spaces and many parameters. Modelling and analysing them is therefore a challenge, particularly if one wants to both take account of stochasticity and develop an analytical approach enabling quantification of various aspects of the system in a more controlled way than is possible by simulation alone. The stochastic kinetics that arise due to random births, deaths and interactions of individual species give rise to Markov jump processes that, in principle, can be analyzed by means of master equations. However, these are rarely tractable and although an exact numerical simulation algorithm is available [1], for the large systems we are interested in, this is very slow.

It is therefore important to develop accurate approximation methods that enable a more analytical approach as well as offering faster simulation and better algorithms for data fitting and parameter estimation. A number of approximation methods aimed at accelerating simulation are currently available. This includes leaping algorithms [2, 3] and algorithms for integration of diffusion equation (or chemical Langevin equation (CLE)) [4] that provide faster simulation. However, these methods do not provide analytical tools for studying the dynamics of the system and they can also be extremely slow for data fitting and parameter estimation. One obvious candidate for overcoming these limitations is the Linear Noise Approximation (LNA). The LNA is based on a systematic approximation of the master equation by means of van Kampen’s Ω-expansion [5] which uses the system size parameter Ω that controls the number of molecules present in the system. The large system size validity of the LNA has been shown in [6], in the sense that the distribution of the Markov jump process at a fixed finite time converges, as the system size Ω tends to ∞, to the LNA probability distribution. The latter distribution is analytically tractable allowing for fast estimation and simulation algorithms. However, the LNA has significant limitations, particularly in approximating long-term behaviour of oscillatory systems.

We show below that for the oscillatory systems that we study, the LNA approximation of the distribution *P_t_* = *P*(*Y*(*t*)|*Y*(0)), of the state *Y* (*t*) of the system at some time *t* becomes inaccurate when the time *t* is greater than a few periods of the oscillation. However, if we rather consider a similar system which in the Ω→∞ limit instead of a limit cycle has an equilibrium point that is linearly stable, then the LNA approximation of *P_t_* remains accurate for a much longer time-scale. For example, in Fig C in S3 Appendix we give an example where the LNA fails in a matter of a period or two for the oscillatory system, but for the corresponding equilibrium system it is very accurate for over a hundred times as long (and probably much longer). Similar behaviour is also observed in other systems and using different measures in [8].

The observation that non-degenerate limit cycles have such linearised stability in the directions transversal to the limit cycle suggests the way forward for oscillatory systems. Our approach exploits the fact that, because of this transversal linearised stability, the distributions *P_t_* for a general class of systems with a stable attracting limit cycle in the Ω→∞ limit are, like the above fixed point systems, similarly well-behaved on long time-scales provided one conditions *P_t_* on appropriate transversal sections to the limit cycle.

We introduce a modified LNA, called the phase-corrected LNA, or pcLNA, that exploits the above observations to overcome the most important shortcomings of the LNA and we develop methods for analysis, simulation and inference of oscillatory systems that are accurate for much larger times. We build on previous work of Boland et al. [9] which uses the 2-dimensional Brusselator system as an exemplar to investigate the failure of the LNA in approximating long-term behaviour of oscillatory systems and presents a method for computing power spectra and comparing exact simulations with LNA predictions of the same phase rather than time. Using various low-dimensional oscillatory systems for illustration, a related analysis has been employed to study the temporal variability of oscillatory systems in the tangental direction of the Ω→∞ limit cycle [10] and/or the amplitude variability in the transversal direction of the limit cycle [11–13]. Other papers derive related descriptions of the asymptotic phase of stochastic oscillators [14, 15].

We extend these results in a number of ways including the following: (i) we develop a theory that treats the general case and provide analytical arguments that justify our approximations and enable computation of trajectory distributions, (ii) we show that the approach is practicable for larger nonlinear systems, (iii) we present a new powerful system-global sensitivity theory for such systems using measures such as the Fisher Information Matrix and the Kullback-Leibler divergence that are analytically computed, (iv) we present a simulation algorithm and show it is as accurate but faster than leaping and integration of diffusion equation algorithms, and (v) we derive the Kalman filter associated with the pcLNA in order to provide a practical way to accurately approximate the likelihood function thus facilitating estimation of system parameters *θ* and predictive algorithms. The approach in [9] uses transversal sections which are normal to the limit cycle. We follow this but in the supplementary information (S1 Sects. & 8.3) we show that for most considerations one can use any transversal to the limit cycle, including those defined in [14, 15].

To illustrate and validate our approach we apply it to a relatively large published stochastic model of the *Drosophila* circadian clock due to Gonze et al. [16] (see S2 Sect. 1). This model involves 10 state variables and 30 reactions and its structure is discussed in S2. The large system limit is given by the differential equation system of 10 kinetic equations that are listed in the supplementary information (S2 Sect. 1) along with the reaction scheme of the system. The stochastic version of the Brusselator system and a stochastic version of a well-studied model of the NF-*κ*B signalling system [17] are also used to illustrate our methods and the results can be found in S3 Appendix.

These systems are free-running oscillators in the sense that they correspond to a limit cycle of an autonomous differential equation in the the Ω → ∞ limit. However, our results also apply to the equally important classes of entrained forced oscillators and damped oscillations. We therefore consider two such systems in S2 and S3: the light-entrained *Drosophila* circadian clock model of [18] which is an example of a forced oscillator and the NF-*κ*B system model [17]. The latter has the extra feature that the analysis is not concerned with a limit cycle but of a transient solution that converges to the limit cycle as time increases. This solution is the biologically interesting one that describes how the system responds to being stimulated by TNF*α*. The supplementary information S1 includes technical derivations and S2 and S3 contain further illustrative figures that we refer to in this paper.

## 2 Results

Stochastic models of cellular processes in signaling and regulatory systems are usually described in terms of reaction networks. A system of multiple different molecular subpopulations has state vector, *Y* (*t*) = (*Y*_1_(*t*),…, *Y_n_*(*t*))^*T*^ where *Y_i_*(*t*), *i* = 1,…, *n*, denotes the number of molecules of each species at time *t*. These molecules undergo a number of possible reactions (e.g. transcription, translation, degradation) where the reaction of index *j* changes *Y* (*t*) to *Y* (*t*) + *v_j_*, *v_j_* ∈ ℝ^*n*^. The vectors *v_j_* are called stoichiometric vectors. Each reaction occurs randomly at a rate *w_j_*(*Y* (*t*)) (often called the intensity of the reaction), which is a function of *Y* (*t*) but also depends periodically on *t* when we are studying forced oscillators. Such systems can be exactly simulated using the Stochastic Simulation algorithm (SSA) [1].

It is common in studying stochastic systems to introduce a system size Ω which is a parameter that occurs in the intensities of the reactions *w_j_*(*Y* (*t*)) and controls molecular numbers (see discussion in S1 Sect. 2). While having a system size parameter is not necessary to apply our methods, it allows one to study the dependence of stochastic fluctuations upon system size and to calculate the deterministic equations that describe the evolution of the concentration vector *X*(*t*) = *Y* (*t*)*/* Ω in the limit of Ω → ∞ (see S1 Sect. 3). Although more general conditions can be used, a condition that will be sufficient for our purposes is that the rates *w_j_*(*Y*(*t*)) depend upon Ω as *w_j_*(*Y*) = Ω*u_j_*(*Y/* Ω) (cf. [5–7]).

In the limit Ω → ∞ the time dependence of *X*(*t*) is given by the solution *x*(*t*) of the differential equation

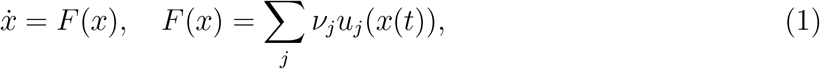

with the appropriate initial condition (see S1 Sect. 3). For free-running oscillators the differential equation (1) is autonomous, whereas for forced oscillators *F* also depends periodically on *t*.

Throughout we will be interested in the case where the solution *x*(*t*) of interest is a stable limit cycle I of minimal period *τ* > 0 given by *x* = *g*(*t*), 0 ≤ *t* ≤ *τ*. We shall also always assume the generic situation for stable limit cycles of autonomous systems in which one of the characteristic exponents of the limit cycle is equal to zero and the rest have negative real part ([19],[20, Sect. 1.5] and S1 Sect. 1). For an entrained forced oscillator all the characteristic exponents of the limit cycle have negative real parts.

Fig 1(A) displays a stochastic trajectory of the concentrations *X*(*t*) = *Y*(*t*)*/* Ω of two of the species of the circadian clock system obtained using exact SSA simulation over a period of time *t* ∈ [0, 8.5*τ*] where *τ* ≈ 26.98 hours is the period of the limit cycle *γ*. Here the system size is Ω = 300, imposing moderate to high levels of stochasticity (see also Table C in S2 Appendix). Results for system sizes Ω = 200, 500 and 1000 are also reported in Fig B in S2 Appendix. Fig 1(B) shows realizations of the key probability distributions

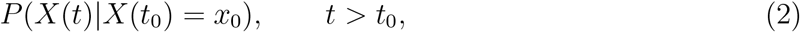

which describe the state of the system at some time *t > t*_0_. It is very rare that accurate analytical approximations for such probability distributions can be derived from the exact Markov-jump process when *t* is large. Furthermore, as we can see, the SSA samples of *P*(*X*(*t*) |*X*(*t*_0_) = *x*_0_) spread along the curved periodic solution, *x* = *g*(*t*), of the limiting (Ω → ∞) deterministic system, implying that for large *t* this distribution is far from being normal and has a complex structure.

**Fig 1.**
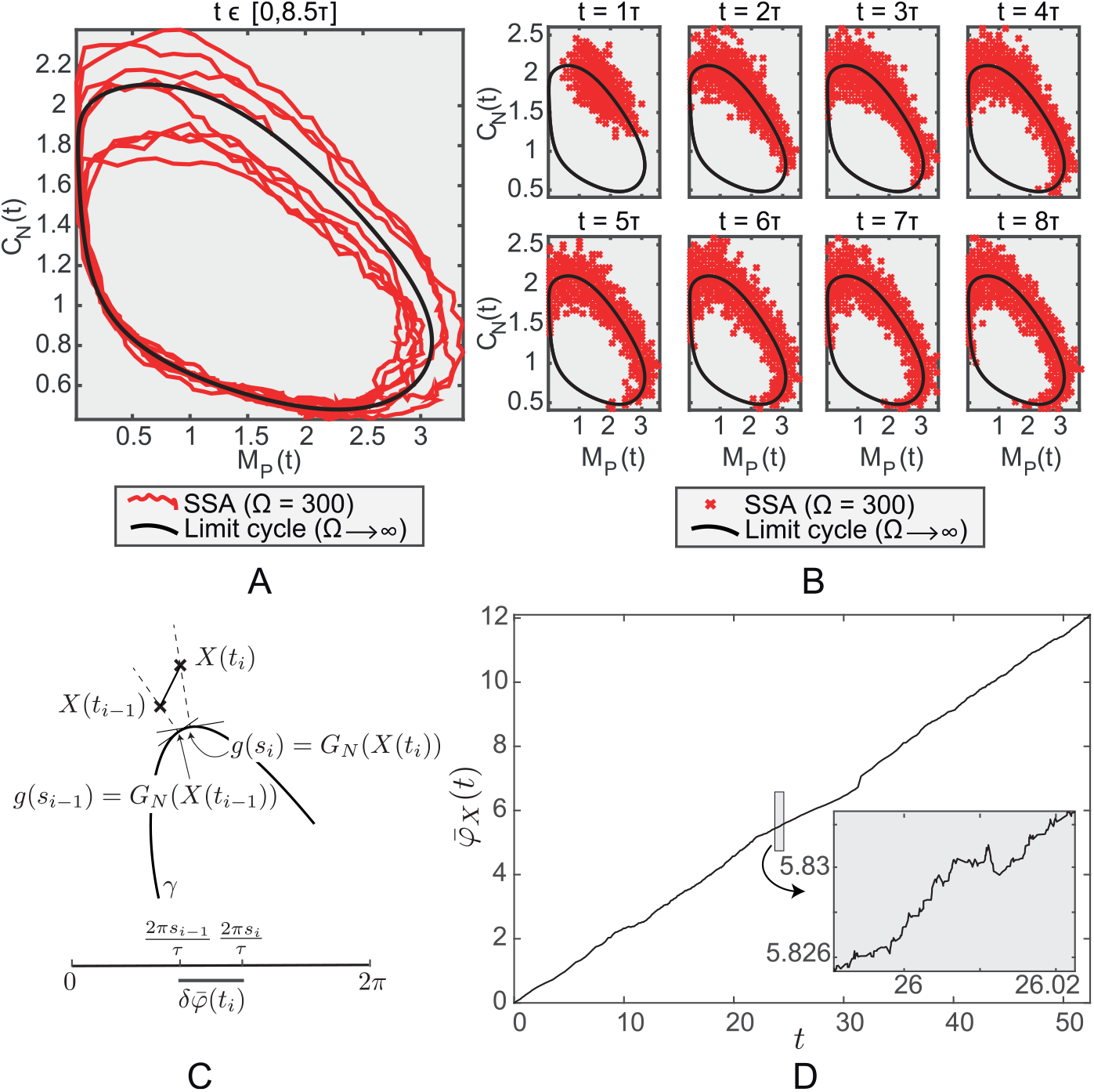
Exact stochastic simulation of the *Drosophila* circadian clock system. (A) A stochastic trajectory obtained by running the SSA over the time-interval *t* ∈ [0, 8.5*τ*] and (B) SSA samples (*R* = 3000) at times *t* = *τ* 2*τ, …*, 8*τ*. The concentrations, *X*(*t*) = *Y*(*t*)*/* Ω, of two (out of 10) of the species are displayed (*per* mRNA *M_P_* (x-axis) and nuclear PER-TIM complex *C_N_* (y-axis)). For this model the system size Ω is Avogadro’s number in units of nM^*-*1^ multiplied by the cell volume in litres and the concentrations are nanomolar. The value of Ω used here is 300 in units of nM which assumes a cell volume of approximately 0.5 × 10^*-*12^ litres. The number of *M_P_* molecules ranges from 0 to 1200 and the number of *C_N_* molecules from 100 to 900. It is only through Ω that the cell volume appears in the equations. The black solid curve is the Ω → ∞, limit cycle solution. (C) Schematic diagram illustrating the mapping *G_N_*(•) and the relative phase *δφ_X_* (•). (D) A plot of the lifted phase function *φ_X_*(*t*) for a trajectory of this system. Note the long-term linear increase combined with frequent reversals

However, we will show that there are important related distributions that can be well approximated even for large times *t*. For example, for each point *x* on the Ω → ∞ limit cycle *γ*, consider the (*n* – 1)-dimensional hyperplane *𝒮_x_* normal to *γ* at *x* ∈ *γ* (i.e. orthogonal to the tangent vector *F*(*x*) at *x*). We will show that the distribution of the intersection points of stochastic trajectories *X*(*t*) with *𝒮_x_* can be well-approximated by a multivariate normal (MVN) distribution that can be relatively easily calculated.

We need to define more precisely what we mean by intersection points. Consider the mapping *G_N_* defined by *G_N_*(*X*) = *x* ∈ *γ* if *X* ∈ *𝒮_x_* (see Fig 1(C)). Suppose that the limit cycle is given by *x* = *g*(*t*) and extend *g*(*t*) to all *t* ∈ ℝ by periodicity. Now consider a stochastic trajectory *X*(*t_i_*), *i* = 0, 1, 2,…. Suppose that *G_N_* (*X*(*t_i-_*_1_)) = *g*(*s_i-_*_1_), *i* > 0. If *G_N_* (*X*(*t_i_*)) = *g*(*s*) then it equals *g*(*s* + *qτ*) for all integers *q*. Choose *q* so that *s_i_* = *s* + *qτ* satisfies *s_i-_*_1_ – *τ*/2 ≤ *s_i_ < s_i-_*_1_ + *τ*/2. Note that the difference between *s_i_* and *s_i-_*_1_ can be both positive and negative and the choice of *s_i_* minimises *|s_i_ – s_i-_*_1_*|*. We define the relative phase *δφ_X_*(*t_i_*) to be 2*π*(*s_i_ – s_i-1_*)*/τ*. With this definition the lifted phase *φx* is defined to be the function

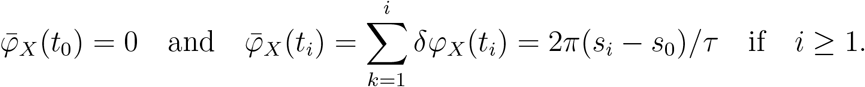

An example of *φx(t)* is shown in Fig 1(C). Now, as shown in that figure, if the system size is not too small, although there will be some reversals, the stochastic process *G_N_* (*X*(*t*)), *t >* 0, will move around *γ* in the direction of the deterministic flow given by (1) so that *φx(t)* increases at a definite positive rate because phase advances exceed retreats. Our approximations (S1 Sect. 6) give that the long term rate is 2 *π/τ* and that the variance of the fluctuations *φx*(*t*) 2*πt/τ* grows linearly with a growth rate that is O(*t /* Ω).

Now consider stochastic trajectories *X*(*t_i_*) with *X*(*t*_0_) ∈ *𝒮_g_*_(*T*0)_ and consider how this trajectory passes through *𝒮_g_*_(*T*1)_. We can assume that *T*_0_ *< T*_1_ ≤ *T*_0_ + *τ* by the periodicity of *g*. The *r*th pass will occur between the first time *t_i_* when

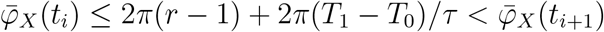

and *t_i_*_+1_. Therefore, we define the *first intersection point of the rth pass* to be

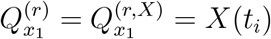

since *X* does not change between *t_i_* and *t_i_*_+1_.

These points of intersection 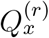 describe the stochasticity of the system around a particular phase *x* of the system. The above ideas work equally well with any transversal to *γ*. For example, the time at which the *i*-th variable is maximal in the deterministic system is given by the transversal submanifold Σ defined by *ẋ*_*i*_ = 0, *ẍ*_*i*_ < 0 so the intersection of the stochastic trajectory with Σ can be regarded as giving the statistics of the maxima of the stochastic trajectory (see Fig 2(A)). Close to the limit cycle, Σ is well-approximated by its tangent space at the point of intersection with the limit cycle. Therefore, these transversal distributions are useful for analysing various aspects of the system.

**Fig 2.**
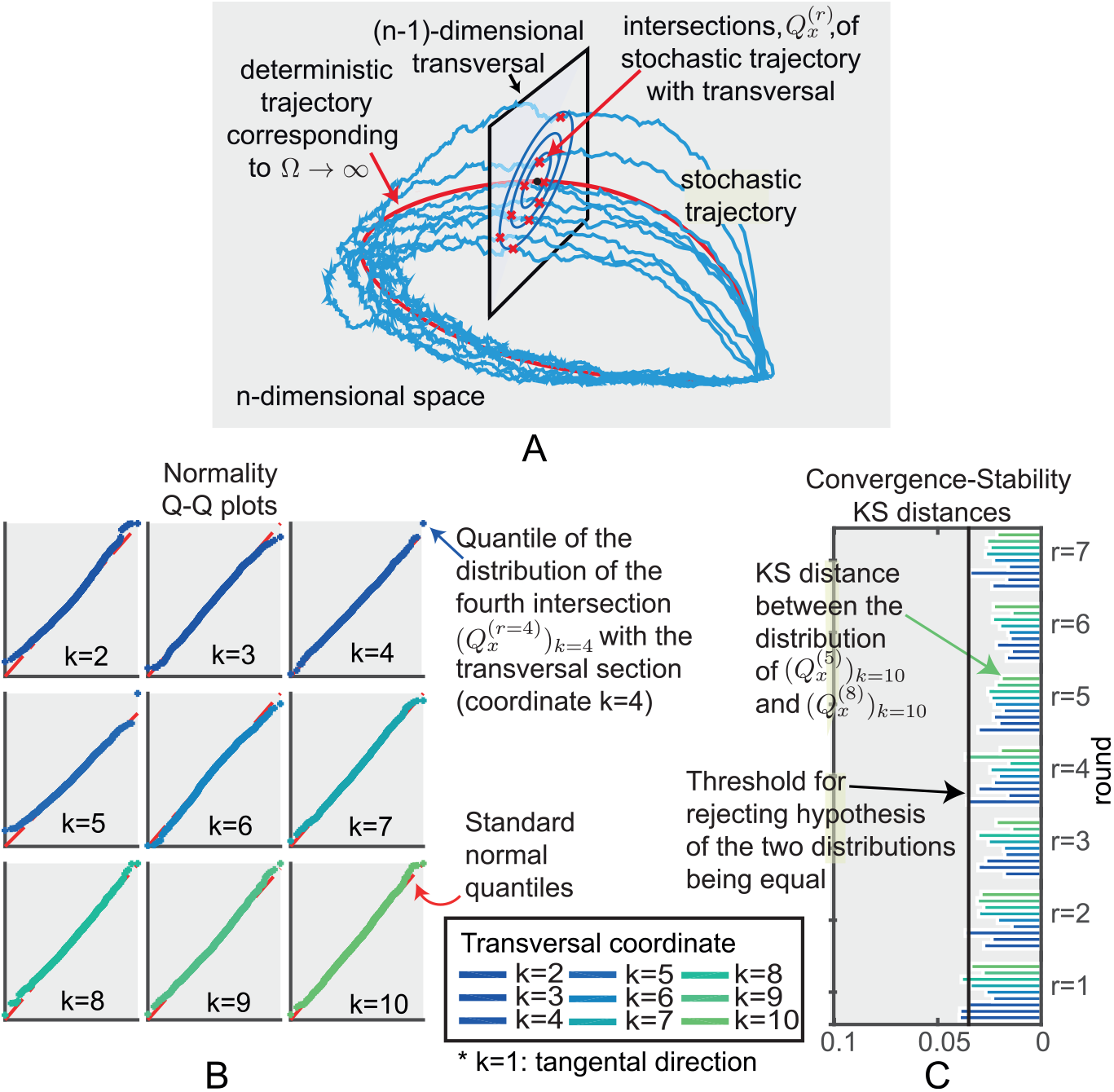
Intersections of the stochastic trajectories with a transversal section 𝒮 _g(t0)_. (A) Schematic representation of the intersections. (B) We consider an adapted coordinate system (*x*_1_,…, *x*_10_) (see S1 Sect. 1) so that (*x*_2_,…, *x*_10_) are orthogonal coordinates on *S_g_*_(*t*0)_ and then consider the distribution *P ^k,r^* of the values of *x_k_* for *k* = 2,…, 10 at the intersection points 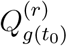. Quantile-Quantile (Q-Q) plots of these distributions for the fourth pass, 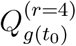 show that the distributions are very close to being normal, (see Fig C in S2 Appendix for similar plots for *r* = 1, 2,…, 8). (C) KS distances between the above distributions for the first *r* = 1, 2,…, 7 rounds of the stochastic trajectory and the distribution in the 8^th^ round (*r* = 8), *k* = 2, 3,…, 10.

Our first observation is that the empirical transversal distributions

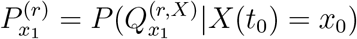

obtained by exact simulation are approximately multivariate normal (Fig 2(B)). Moreover, as *r* increases 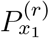 and 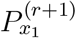 are hardly distinguishable and appear to converge to a fixed, approximately normal transversal distribution as *r* increases (Fig 2(C)). Similar results hold if we use a different family of transversal sections to γ as explained in S1 Sect. 8.2 & 8.3.

A natural question that arises is whether one obtains a different distribution when instead of taking the first intersection point of the *r*th pass one takes a later intersection point of the same round. This is addressed in S1 Sect. 12 where we show that in exact simulations there appears to be no difference as would be expected from the LNA approximation.

## The Linear Noise Approximation (LNA)

The convergence of the transversal distributions to approximately normal distributions naturally raises the question of whether asymptotic approximation methods such as the LNA, which provide multivariate normally distributed approximation of the stochastic system, can be used to accurately approximate these transversal distributions.

The LNA as formulated by [6] is derived directly from the underlying Markov jump process and is valid for any time interval of finite fixed length. It is based on the ansatz

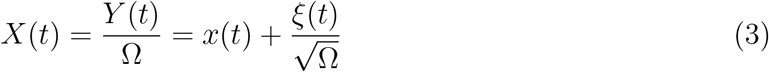

where *x*(*t*) is a solution of the limiting (Ω → ∞) deterministic system (1) and 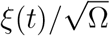 describes the stochastic variations. In our case we always take *x*(*t*) to be the periodic solution *g*(*t*). A key aspect of this ansatz is that *ξ*(*t*) satisfies a linear stochastic differential equation that is independent of Ω, with drift and diffusion matrices that are functions of the deterministic solution *g*(*t*). Details are given in S1 Sect. 4 & 5.

Given an initial time *t*_0_ and an initial condition *ξ*(*t*_0_) for *ξ*, the LNA determines the distribution, of ξ (*t*), *t > t*_0_, and hence 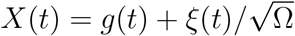 that we respectively denote by *P*_LNA_(ξ (*t*) |*t*_0_, ξ (*t*_0_)) and *P*_LNA_(*X*(*t*) |*t*_0_, *ξ*(*t*_0_)). If *ξ*(*t*_0_) is only known up to a multivariate normal (MVN) distribution *P*_0_ then we denote these distributions, respectively, by *P*_LNA_(*ξ*(*t*) |*t*_0_, *ξ*(*t*_0_) ~ *P*_0_) and *P*_LNA_(*X*(*t*) | *t*_0_, *ξ*(*t*_0_) ~ *P*_0_). Details of how to calculate these distributions are given in S1 Sect. 4. Each of the above distributions is MVN enabling analytical approaches, for example in analysing the stochastic sensitivities of the system.

If we fix *t > t*_0_ then as Ω → ∞ the true distribution of *ξ* converges to the distribution *P*_LNA_(*ξ*(*t*) | *t*_0_, *ξ*(*t*_0_)) (see e.g. [5]). However, one most certainly cannot reverse the limits i.e. for a fixed Ω one cannot expect the approximation to hold for large time *t* → ∞. As we now show, this is certainly the case for oscillators and we aim to overcome this limitation by developing methods that remain accurate for much larger times than the LNA.

We first consider the distribution *P* (*X*(*t*) | *t X*(0) = *x*_0_) and compare this for SSA simulated samples and the LNA at a sequence of times *t* = *τ*, 2*τ, …*, 8*τ* and for an arbitrary (fixed) initial state *x*_0_ ∈ *γ*. As we can see in Fig 3, the LNA fits the SSA simulations relatively well in the short run (*t* = *τ*), but as time progresses the Kolmogorov-Smirnov (KS) distance between the two distributions for each state variable for the LNA and the SSA increases substantially beyond the threshold level (see Fig 3(B)). The LNA predictions spread along the tangental direction and therefore fail to accurately reflect the SSA samples that have instead spread along the curved limit cycle.

**Fig 3.**
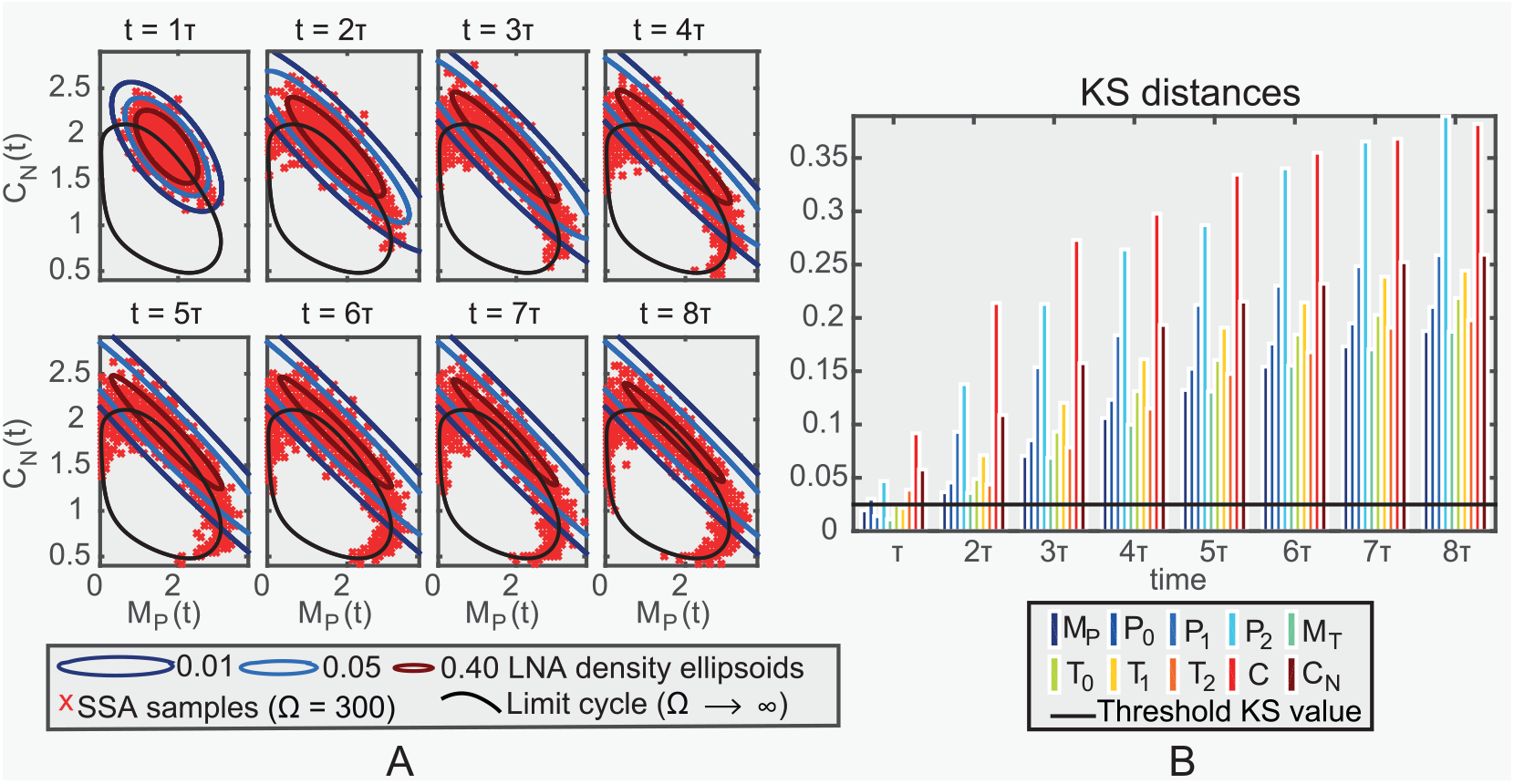
Comparison between LNA and exact simulations. (a) Samples (in nanomolar concentrations) obtained from the SSA simulation algorithm (red crosses) and 0.01, 0.05, 0.40 contours of the LNA probability density (black ellipsoids) at fixed times, *t* = *τ*, 2*τ, …*, 8*τ* (*τ*: minimal period), for the circadian clock system (*M_P_ per* mRNA; *C_N_* nuclear PER:TIM complex *P*_0_ & *T*_0_ PER & TIM protein; *P*_1_ & *T*_1_ phosphorylated PER & TIM protein; *P*_2_ & *T*_2_ twice phosphorylated PER & TIM protein; *C* cytoplasmic PER:TIM complex, units as in Fig 1). The limit cycle ODE solution is also displayed (black solid line). (b) KS distance between the empirical distribution of SSA samples and the LNA distribution of each species (different colors, see legend) at the fixed times. The threshold level is also displayed (black solid line). The system size is Ω = 300.

On the other hand, as we saw earlier, the transversal distributions 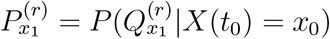 of the *Drosophila* circadian clock system are approximately normal (Fig 3(B)). We next derive an approximation of 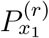 under the LNA and show that it accurately approximates 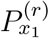 for the *Drosophila* circadian clock, Brusselator and NF-*κ*B systems.

## Calculating transversal distributions

We now generalise slightly and consider the set of stochastic trajectories *X* where the initial conditions *X*(*t*_0_) have a MVN distribution *P*_0_ that is supported on the normal transversal section *𝒮*_*g(t_0_*_ (denoted *X*(*t_0_*) ~ *P*). We consider how to approximate the distribution 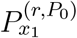 of the intersection points 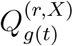 of these trajectories with the normal transversal section *𝒮*_*g*(*t*_1_)_, *t*_1_ > *t*_0_. As an approximation we take the conditional distribution

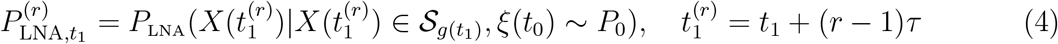

given by conditioning 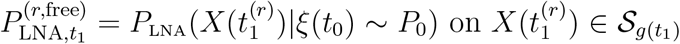. It gives a MVN distribution supported on ***𝒮**g*(t1).

In S1 we show that, although, for free-running oscillators, 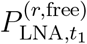 diverges as *r* → ∞, the mean and covariance of the MVN transversal distribution 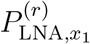 converge exponentially fast to those of a MVN distribution 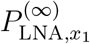 (S1 Sect. 8, cf. Fig 4). The distribution 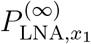 is a fixed point in the sense that if the distribution of *X*(*t*_1_ + *τ*) is given by the LNA using as initial condition 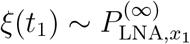 then conditioning on *X*(*t*_1_ + *τ*) ∈ *𝒮_g_*_(*t*1)_ gives

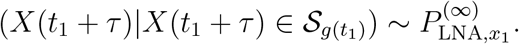

Using this fact enables us to calculate 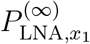 directly because we show in S1 that its mean and covariance matrix satisfy a simple fixed point equation that is easily solved numerically (S1 Sect. 9.1).

**Fig 4.**
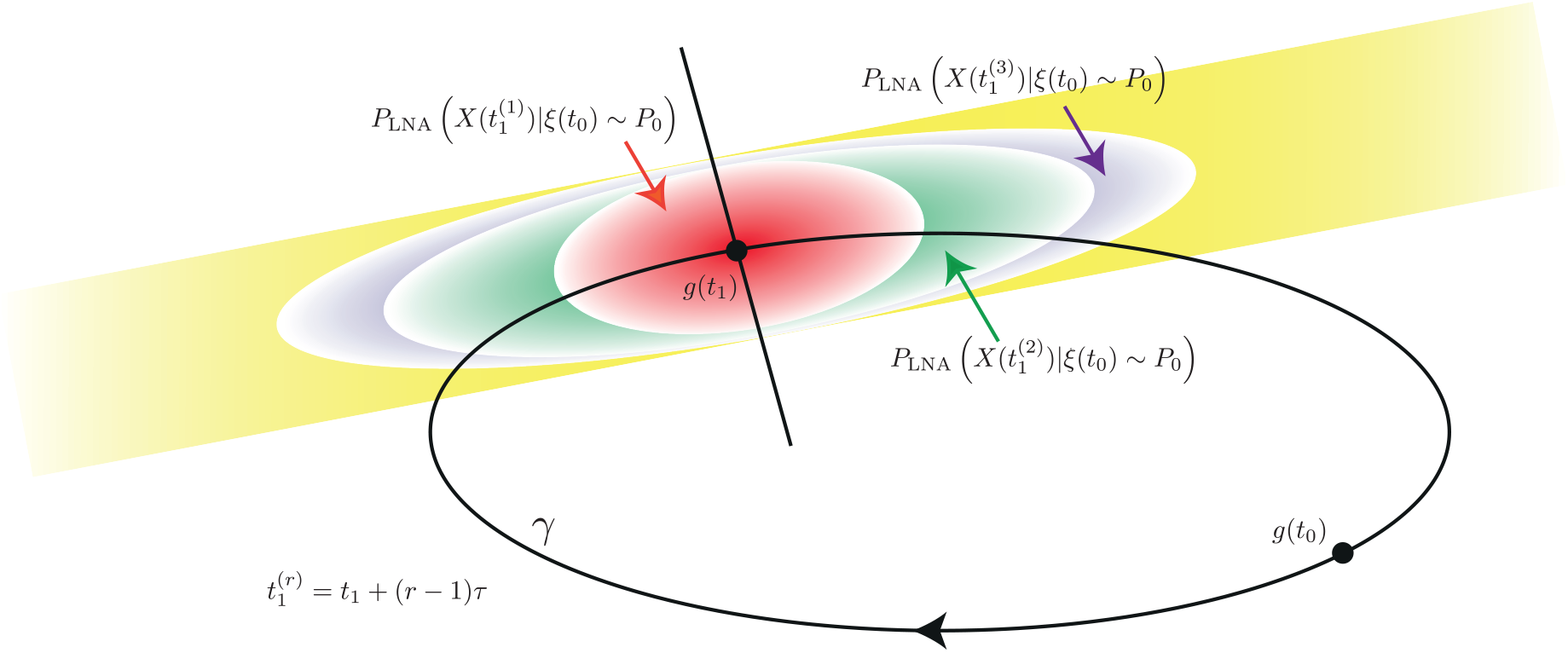
Schematic diagram of the distributions 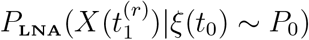 as *r* increases. The yellow distribution represents the limit as *r* → ∞. Although, for free oscillators the latter distribution diverges, the corresponding conditional distributions on the transversal converge.

The reader will note that in (4) we approximate by conditioning on 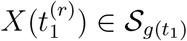 whereas we should have conditioned on *X*(*t*) ∈ 𝒮_*g*(*t*1)_ for arbitrary *t* corresponding to the *r*th round. In S1 Sect. 7 we argue that the error in the mean and variance of the distribution due to taking 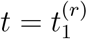 is O(Ω^−1^).

The question remains as to how well these distributions capture the exact simulation transversal distribution 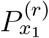 This is addressed in Fig 5 where it is shown that the fit is excellent even for Ω as low as 300. The fit is even better for higher system sizes (Fig E in S2 Appendix). In S3 we also show similar low Ω results for the Brusselator (Fig B in S3 Appendix) and the NF-*κ*B system (Fig E in S3 Appendix). The result is also true for the light-entrained *Drosophila* circadian clock system (see Fig J in S2 Appendix) and the transient oscillations of the NF-B system (see Fig H in S3 Appendix). Thus we note that although the LNA cannot be used directly to accurately compute *P* (*X*(*t*)*|X*(0)) for a fixed Ω and increasing *t*, using it to compute the transversal distributions provides accurate estimates of 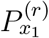 for much larger times *t*_1_ + *rτ*.

**Fig 5.**
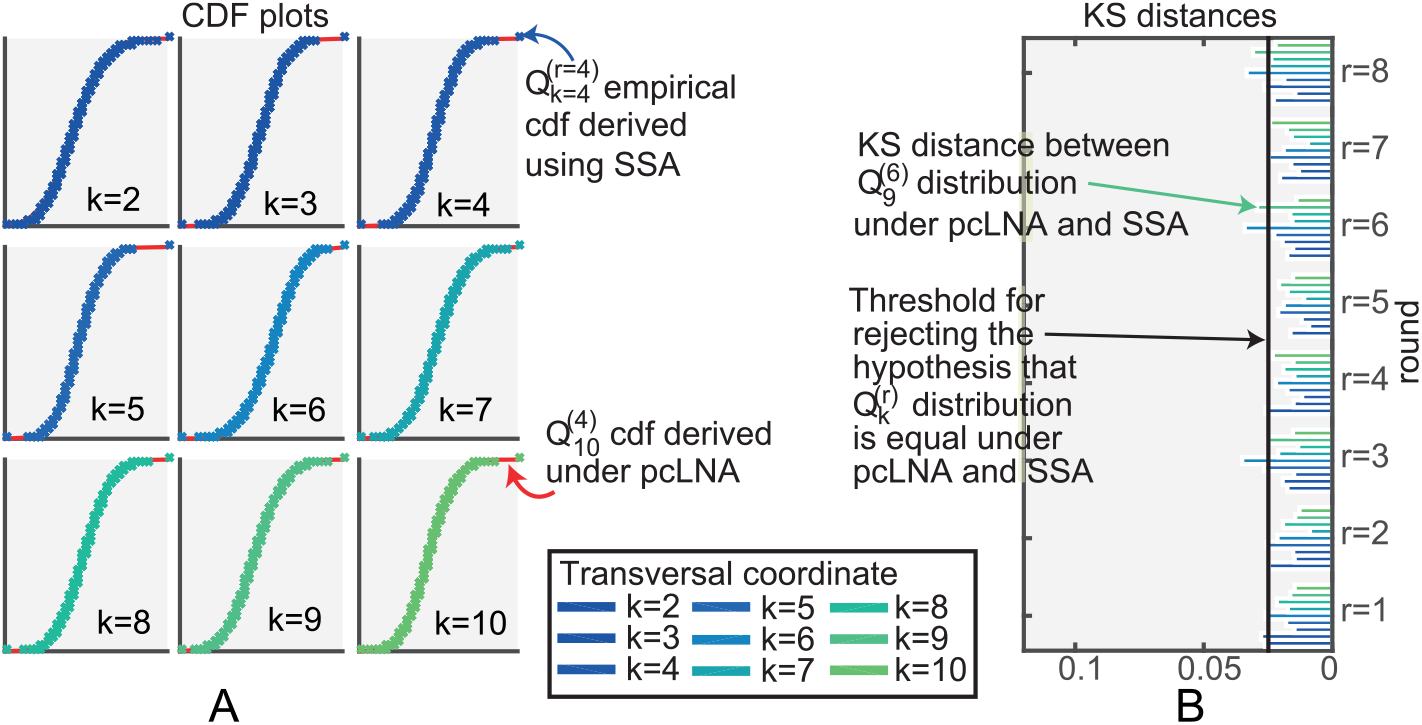
Comparison of pcLNA and exact transversal distributions for the *Drosophila* circadian clock (Ω = 300). (A) CDF plots of the distributions *P ^k,r^* as in Fig 2 for the pcLNA (red line) and the SSA (empirical CDF, crosses) for *k* = 2, 3,…, 10 and round *r* = 4 (see Fig D in S2 Appendix for *r* = 1, 2,…, 8). (B) KS distances between the corresponding distributions under pcLNA and SSA, *r* = 1, 2,…, 8, *k* = 2, 3,…, 10.

Moreover, in S1 Sects. 8.2 & 8.3 we also explain why the convergence of the distribution on normal hyperplanes implies convergence on other transversal sections to *γ*.

In S1 Sect. 8.4. we explain that in contradistinction to free-running oscillators, for entrained forced oscillators, *P_r_* = *P*_LNA_(*X*((*r* – 1) *τ* + *t*_1_) *x*_0_, *ξ*_0_ ~*P*_0_) converges as *r* → ∞ so that, under the LNA, the phase fluctuation have a variance that is bounded independently of *r*. The corresponding conditional distribution is therefore a correspondingly good approximation to the transversal distribution 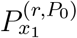(see Fig J in S2 Appendix). However, it does not mean that 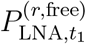 is a good approximation to the corresponding distribution 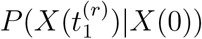 for an exact simulation. In fact, we show in Fig I in S2 Appendix that 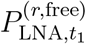 is a poor approximation of the empirical distribution 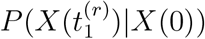 derived from exact simulations for the light-entrained *Drosophila* circadian clock (Ω = 300). The bounded variance of the phase fluctuations as *r* → ∞ for forced oscillators is the basic mechanism behind the population-level entrainment of stochastic oscillators introduced in [21].

## Stochastic fluctuations in periods and timing

We now consider the fluctuations *δt* in the time taken for the lifted phase of a stochastic trajectory to go from a given phase *φ*_1_ to a greater one *φ*_2_. If *φ*_2_ – *φ*_1_ = 2*r π* then this corresponds to the time taken to perform *r* cycles.

In Fig 6 we give an example using the *Drosophila* circadian clock model where we take *φ*_1_ to be 0 (with a fixed initial condition *x*_0_ ∈ *γ*) and *φ*_2_ = 2*πr* for *r* = 1,…, 8. The distributions of *δt* appear to be very close to normal and the variance appears to grow linearly with *r*. We also consider the case where *φ*_1_ = 2(*r* – 1)*π* and *φ* _2_ = 2*r π* for *r* = 1,…, 8. Again the distributions are approximately normal but the variances are approximately constant (Fig 6(C)). Because for a given *r* the trajectory has done *r* – 1 cycles before reaching the lifted phase *φ*_1_ the distribution of the state at this phase is changing with *r*. We expect that this distribution is converging with increasing *r* and this result (Fig 6(C)) is in accordance with this.

**Fig 6.**
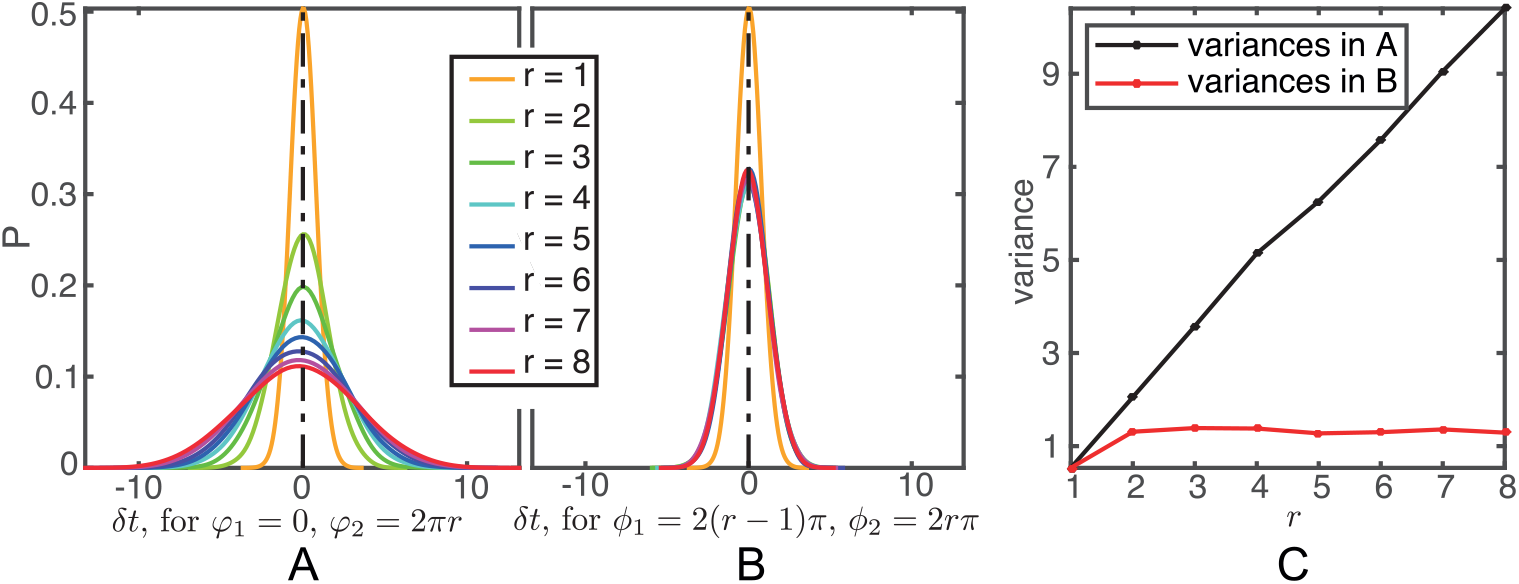
Exact empirical distribution of the fluctuations *δt* in *Drosophila* circadian clock system size Ω = 300. (A) The (smoothed) empirical density function of *δt* in the time taken for the lifted phase of a stochastic trajectory to go from *φ*_1_ = 0 to *φ*_2_ = 2*r π*, *r* = 1, 2,…, 8. (B) The (smoothed) empirical density function of the fluctuations in the time taken for the lifted phase of a stochastic trajectory to go from *φ*_1_ = 2(*r* – 1)*π* to *φ*_2_ = 2*r π*, *r* = 1, 2,…, 8. (C) The variance of the distributions in (A) and (B).

In S1 Sect. 6 we approximate the statistics of *δt* using the LNA and show that as a random variable 5*t* is approximately normal with mean that is O(Ω^*-*3/2^) and we also calculate its variance up to terms that are O(Ω^*-*3/2^) and the extent of its divergence from normality.

All points with a given lifted phase *φ*_1_ lie in a particular transversal *𝒮_g_*_(*t*1)_ with 0 ≤*t*_1_ *< τ*. If *t*_2_ = *t*_1_ + (*φ*_2_ *-* φ_1_)*τ*/2*π*, then, the mean and variance of *δt* can be calculated in terms of *t*_1_ and *t*_2_. If the initial conditions *ξ*(*t*_1_) are MVN distributed on *𝒮_g_*_(*t*1)_ with mean 0 and covariance *V*_1_, this variance is (1/α^2^ Ω) V̌ _11_ + O(Ω^*-*3/2^) where V̌ _11_ = V̌ (*t*_1_, *t*_2_)_11_,

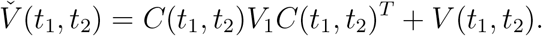

written in adapted coordinates at *g*(*t*_2_) (see S1 Sect. 1). All terms on the right hand side of this equation are defined in S1 Sect. 4. The above exact simulations of the *Drosophila* circadian clock agree with these theoretical predictions. It is easy to see (cf. S1 Sect. 8) that *V̌*_11_ grows roughly linearly with *t*_2_ *-t*_1_.

## pcLNA Simulation Algorithm

Given the ability to accurately approximate the transversal distributions and the results in [9] we realised it should be possible to use this to construct a rapid simulation algorithm. The linear increase of the variance of the deviations *δt*, or equivalently, the linear growth in the variance of the deviation of the lifted phase *φ*_*X*_(*t_i_*) from 2*πt_i_/τ*, indicates the reasons for the long-time failure of the standard LNA. It is unable to cope with the increasing phase deviations. This motivates the phase correction approach used in the simulation algorithm we now define.

The approach is to amend the LNA Ansatz *X*(*t*) = *g*(*t*)+ Ω^*-*1/2^ *ξ*(*t*) to *X*(*t*) = *g*(*s*)+ Ω^*-*1/2^ *κ*(*s*) where *g*(*s*) = *G_N_*(*X*(*t*)) and to use resetting of *t* to *s* to cope with the growth in the variance of *φ*_*X*_(*t_i_*) – 2*πt_i_/τ* keeping the LNA fluctuation *κ*(*s*) normal to *γ*. While for free-running oscillators the variance of *ξ*(*t*) grows without bound as *t* increases, *κ*(*s*) has uniformly bounded variance.

The pcLNA simulation algorithm iteratively uses standard LNA steps of length Δ*τ* to move from a state *X*(*s_i-_*_1_) to a new state *X*(*s_i-_*_1_ + Δ*τ*) = *X_i_*, *i* = 1, 2,…. After each LNA step, the phase of the system is reset or “corrected” such that *g*(*s_i_*) = *G_N_* (*X_i_*) and the (global) fluctuations *ξ*(*s_i-_*_1_ + Δ*τ*) = Ω^*-*1/2^(*X_i_ g*(*s_i-_*_1_ + Δ*τ*)) are replaced by the normally transversal fluctuation *K*(*s_i_*) = Ω^*-*1/2^(*X_i_* –*g*(*s_i_*)) which are MVN distributed and, as we showed in the previous section, approximate well the transversal fluctuations under the exact Markov Jump process.

The steps of the pcLNA simulation algorithm are described next in more detail (see also Fig 7).

1. Choose a time-step size Δ*τ >* 0.
2. Input **initial conditions** *κ*(*s*_0_) and *X*_0_ = *g*(*s*_0_)+ Ω^*-*1/2^ *κ*(*s*_0_).
3. **For iteration** *i* = 1, 2,….

a. sample *ξ*(*s_i-_*_1_ + Δ*τ*) from MVN(*C_i_*(*s_i-_*_1_), *V_i_*);
b. compute *X_i_* = *g*(*s_i-_*_1_ + Δ*τ*)+ Ω^*-*1/2^ ξ (*s_i-_*_1_ + Δ*τ*);
c. set *s_i_* to be such that *G_N_* (*X_i_*) = *g*(*s_i_*) and *κ_i_* = Ω^1/2^(*X_i_ – g*(*s_i_*)).

**Fig 7.**
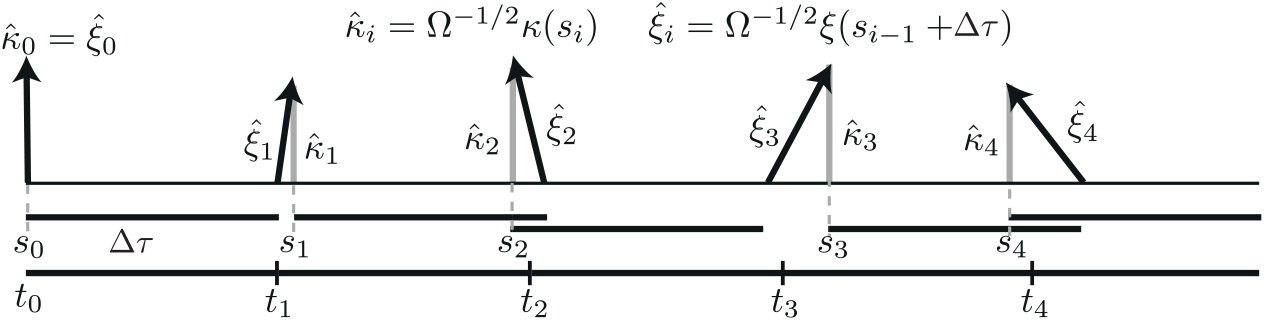
The main step in the pcLNA algorithm. The solid horizontal bars below the horizontal axis are all of length Δ*τ*, the basic time step of the algorithm. The black arrows show *ξ̂*_*i*_ = Ω^*-*1/2^ *ξ*(*s_i-_*_1_ + Δ*τ*) and the grey arrows *κ̂*_*i*_ = Ω^*-*1/2^ (*s_i_*). Having calculated *κ̂*(*s_i-_*_1_) one uses *κ*(*s_i-_*_1_) as the initial state and updates it using the LNA and a time-step Δ*τ* to obtain *ξ* at *s_i-_*_1_ + Δ*t*. Then *ξ*(*s_i-_*_1_ + Δ*τ*) is replaced by *κ*(*s_i_*) so that *g*(*s_i-_*_1_ + Δ*τ*)+ Ω^*-*1/2^ *ξ*(*s_i-_*_1_ + Δ*τ*) = *g*(*s_i_*)+ Ω^−1/2^*κ*(*s_i_*) where *κ*(*s_i_*) is normal to the limit cycle. Therefore, *s_i_* gives the phase of *κ*(*s_i_*) and the corresponding time is *t_i_* = *t*_0_ + *i* Δ*τ*.

In the for loop *C_i_* = *C*(*s_i-_*_1_, *s_i-_*_1_ + Δ*τ*) and *V_i_* = *V*(*s_i-_*_1_, *s_i-_*_1_ + Δ*τ*) are the drift and diffusion matrices in the linear SDE describing the evolution of the noise process *ξ*(*t*) under the LNA (see S1 Sect. 4).

The simulated sample *X_i_* corresponds to time *t_i_* = *t*_0_ + *i* Δ*τ*, *i* = 1, 2,…, where *t*_0_ is the initial time. The time *t_i_* is not necessarily equal to the phase *s_i_*, defined by the relation *g*(*s_i_*) = *G_N_*(*X_i_*), which is stochastic and has variance linearly increasing with the time step Δ*τ*.

If one wants to record simulated trajectories at a finer time-scale than Δ*τ* then one can run the algorithm with Δ*τ* replaced by Δ*τ /M* for some integer *M >* 1 and only carry out the phase correction in step 3(c) every *M* th step and at all the other steps just proceeding as in the standard LNA (ignoring step 3(c)). This gives the same distribution as if the intermediate points had not been calculated because of the transitive nature of the LNA i.e. the distribution *P*_LNA_(*X*(*s* + *t*) |*X*(0)) is equal to the distribution *P*_LNA_(*X*(*t*) |*X*(*s*) ~ *P*_LNA_(*X*(*s*) |*X*(0))). In the simulation results described below the time-step Δ*τ* = 6 hours and *M* = 3 so that there are *τ* /6 ≈ 4.5 corrections in every round of the limit cycle. The effect of less frequent correction is studied in S2 Sect. 5.

### Comparisons to other simulation algorithms

We compared the pcLNA simulation algorithm in terms of both CPU time and precision of the approximation with some of the most important alternatives, the tau-leap method and integration of the chemical Langevin equation (CLE) method. Exact simulations produced by the SSA are also used as a reference for the comparison.

For the tau-leap simulation we use the algorithm described in [3], which is a refinement of the original tau-leap method first proposed in [2]. In [3] the authors suggest an optimal method to compute the largest possible time step such that the leap condition is still approximately satisfied. The leap condition ensures that the state change in any time step is small enough so that no rate function will experience a macroscopically significant change in its value. The error of this approximation is controlled by a parameter ∈. For integrating the CLE described in [4], we use the classical Euler method (see [32]) with a fixed time step Δ*t*. The integration of the CLE can be done using methods that include higher-order terms in the integration and this has been shown to improve the speed of implementation in low-dimensional systems, albeit with a cost in the complexity of the algorithm (see [33, 34]). However, we are not aware of any implementations of these methods for high-dimensional systems such as those considered here, or of comparisons of such methods to the Euler method for such problems.

For both the tau-leap and the CLE approximation we explored different values of their parameters ∈ and Δ*t* to attain a good balance of precision and CPU time. Here we present the results for the largest values of both ∈ and Δ*t*, hence smallest CPU time, which attain similar performance with the pcLNA algorithm in terms of precision. If little improvement could be achieved in terms of precision by lowering either ∈ or Δ*t*, the larger values are preferred.

The algorithms are implemented for a fixed time-interval (8.5 times the period of the limit cycle of the system) and the comparison is made at 8 fixed time points using the KS distances between the empirical distribution of each algorithm and the empirical distribution derived using the SSA simulations. Note that for all approximation algorithms considered here, the probability of generating negative populations is non-zero and there are a number of methods for dealing with this. Our simple approach is described in S1 Sect. 13.

Figure 8 displays the median CPU times for a single trajectory simulation in *t* ∈ [0, 8.5*τ*], under the competing approaches for different system sizes, Ω = 300, 1000, 3000, along with a comparison in terms of precision for Ω = 300 (see Figs G & H in S2 Appendix for Ω = 1000 & 3000). A sample of size *R* = 2000 is produced for each algorithm. We see that the precision of all approximation methods is fairly similar, with their empirical distributions almost indistinguishable to exact simulations. In terms of CPU times, we see that the SSA is much slower than the other algorithms particularly for large system sizes. The tau-leap offers some improvement to the CPU times but this is relatively small compared to the CLE approximation and especially the pcLNA algorithm. One reason for this relatively small improvement for the tau-leap algorithm is the stiffness of the considered system, a property that is however very common in biological systems and it is known to slow down the tau-leap method by requiring small values of the ∈ parameter to ensure that the leap condition is satisfied and hence the approximation is fairly precise. Note that for similar reasons, a small Δ*t* was necessary to achieve good precision with the CLE approximation.

**Fig 8.**
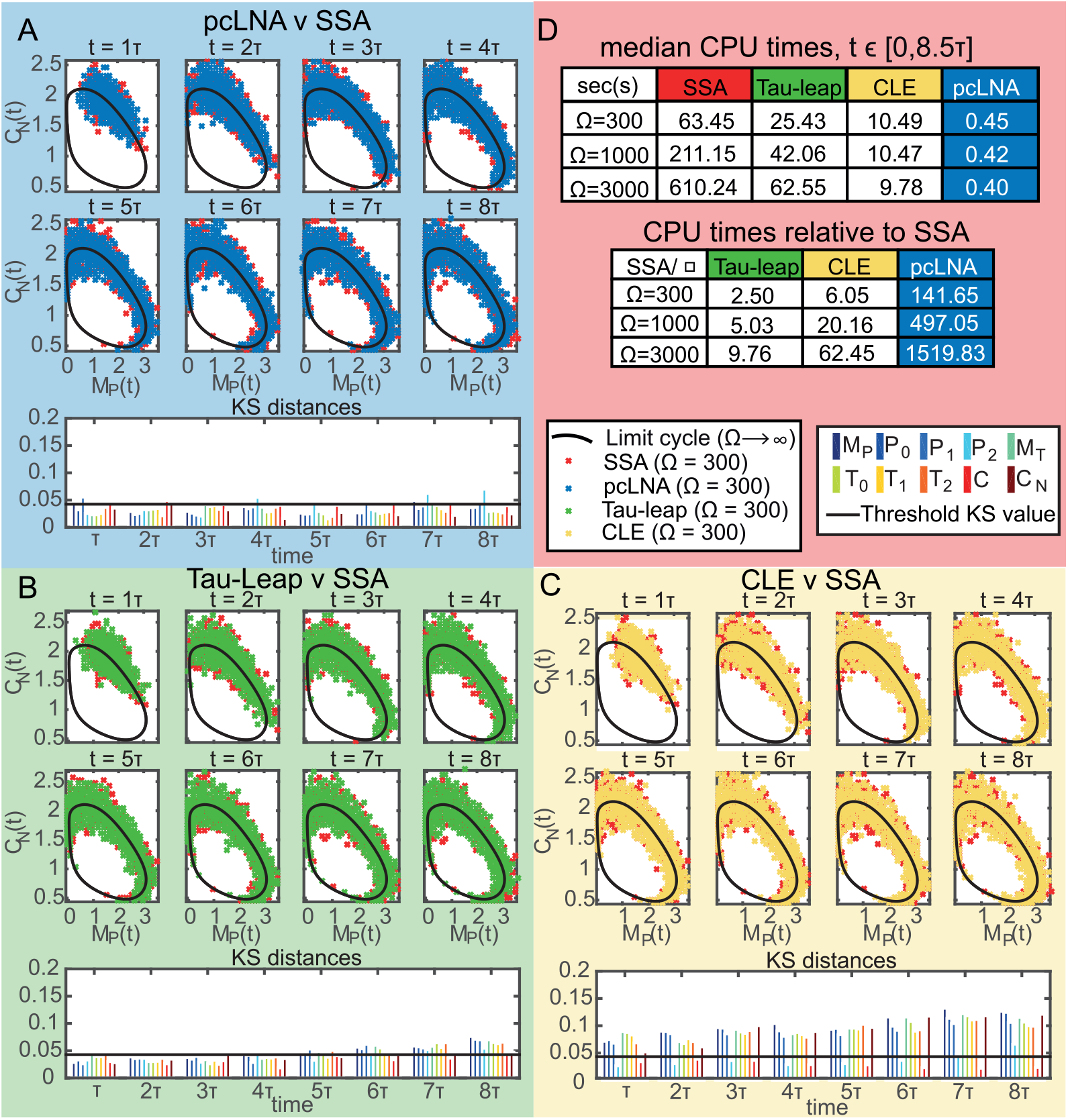
Comparison between pcLNA, tau-leap and CLE simulation algorithms for the *Drosophila* circadian clock. Panels (A), (B) and (C) contain the samples (in concentrations, units as in Fig 1) produced by respectively the pcLNA, tau-leap (∈ = 0.002) and CLE (Δ*t* = 0.002) algorithms and the exact simulation (SSA) at time-points *t* = *τ*, 2*τ, …*, 8*τ*, for Ω = 300, along with the KS distances between the empirical distributions of each approximation and the SSA for each system variable (coloured bars; variable names as in Fig 3). Panel (D) provides the median CPU times for a single trajectory run in the time-interval [0, 8.5*τ*] under the different simulation algorithms along with the ratio of the median CPU time under SSA and each approximation algorithm.

As we can see in Fig 8, the pcLNA algorithm is about 24 times faster than the CLE approximation, tens to hundreds of times faster than the tau-leap and hundreds to thousands of times faster than the SSA. In our simulations, it took 0.4sec to derive this long-time trajectory, which means that in about 7 minutes one can generate more than 1000 trajectories of this large system over a long-time compared to about 2.7 hours with CLE approximation and much longer times for the other methods. Therefore, the pcLNA offers a substantial improvement in CPU times compared to standard approaches in simulating oscillatory systems without compromising the precision of the simulation substantially.

Perhaps more importantly, this pcLNA simulation algorithm has the advantage of being based on an analytical framework that allows calculation of some key distributions. Therefore, it enables more rigorous methods for assessing accuracy and robustness.

## Analysis and inference of oscillatory systems using pcLNA

The derivation of analytical expressions of the transversal distributions allows us to analyse various aspects of the stochastic behaviour of these systems that can possibly involve a large number of variables and parameters. Here we illustrate the use of pcLNA transversal distributions to perform such an analysis. We begin by describing the pcLNA joint distribution of multiple intersections to possibly different transversal sections on the limit cycle and then discuss Fisher information, sensitivity analysis and estimation by Kalman filtering.

### Joint distribution on multiple transversals

Consider *q* phase states of the limit cycle *x_i_* = *g*(*t_i_*), *i* = 1,…, *q*, on *γ* where 0 ≤ *t*_1_ *< t*_2_ *<… < t_q_ < τ*. If *X*(*t*) is a stochastic trajectory, we consider how it meets the transversal sections at the *x_i_* as *t* increases using the lifted phase function *φ*_*X*_. We can talk of the times when *G_N_*(*X*(*t*)) first takes the phase *x_i_* during the *r*-th revolution of *G_N_*(*X*(*t*)) around *γ*. Using *φ*_*X*_ to define the points 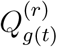 we have that it first meets *𝒮_x_i__* in 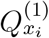 for *i* = 1,…, *q*. If *i< q* then we let 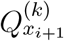 denote the first point in *𝒮_x_i_+1_* that *X* meets after it leaves 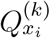. If *i* = *q* then the next transversal it meets is *𝒮_x_*_0_ and the intersection point is 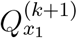. In this way (see Fig 9) we derive a sequence of intersection points 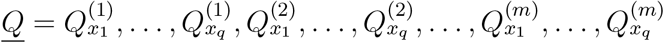.

**Fig 9.**
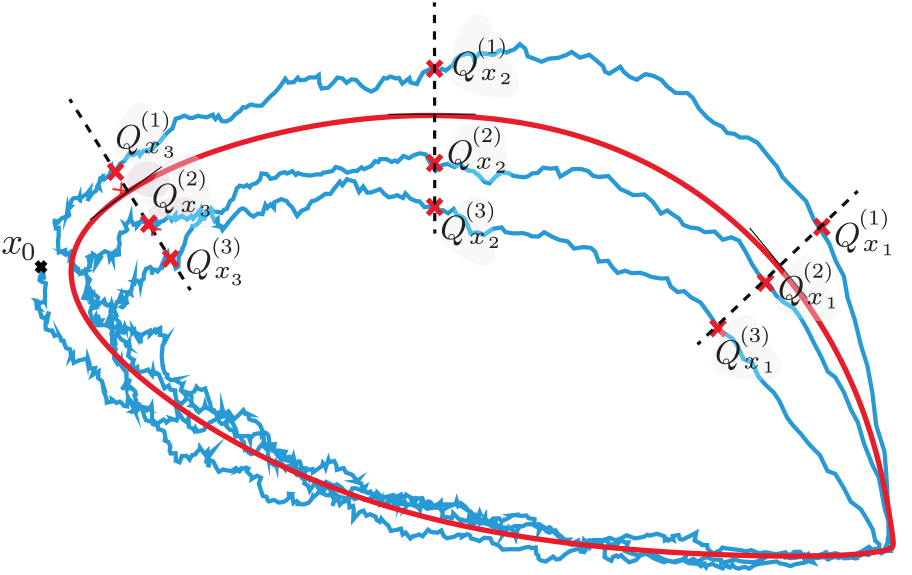
The sequence *Q* in two-dimensions. The stochastic trajectory *X*(*t*) (blue line), initiated from *x*_0_, intersects each of the transversal sections *S_x_*_1_, *S_x_*_2_ and*S_x_*_3_ (dashed lines) of the limit cycle (red solid line) three times, following a path 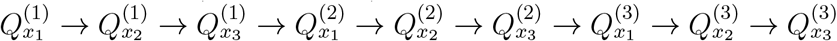

We shall be interested in the distribution

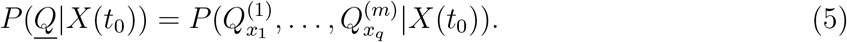

Remarkably, in our approximation, this distribution is MVN with a covariance matrix whose inverse has a simple tridiagonal form in terms of the drift and diffusion matrices coming from the LNA (S1 Sects. 9.2 & 9.3).

The fact that the above transversal distributions are MVN allows us to analytically compute the Fisher Information matrix and associated quantities that can be used to perform a stochastic sensitivity analysis of oscillatory systems.

## Fisher Information

Fisher Information quantifies the information that an observable random variable carries about an unknown parameter *θ*. If *P* (*X, θ*) is a probability distribution depending on parameters *θ*, the Fisher Information Matrix (FIM) *I* = *I_P_* has entries

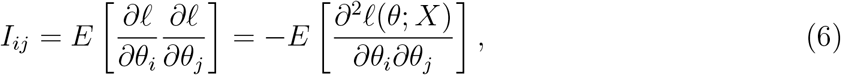

where *ℓ* = log *P*, and *θ_i_* and *θ_j_* are the *i*th and *j*th components of the parameter *θ*. If *P* is MVN with mean and covariance *μ* = *μ*(*θ*) and Σ = Σ (*θ*) then

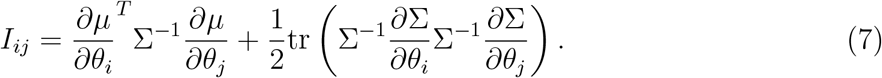

The FIM measures the sensitivity of *P* to a change in parameters in the sense that

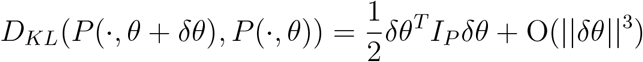

where *D_KL_* is the Kullback-Leibler divergence. The significance of the FIM for sensitivity and experimental design follows from its role in (6) as an approximation to the Hessian of the log-likelihood function at a maximum. Assuming non-degeneracy, if *θ*^*^ is a parameter value of maximum likelihood there is a *s* × *s* orthogonal matrix *V* such that, in the new parameters *θ′ = V*. (*θ*– *θ*^*^),

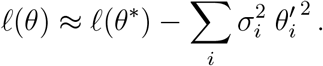

for *θ* near *θ*^*^. From these facts it follows that the 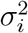 are the eigenvalues of the FIM and that the matrix *V* diagonalises it. If we assume that the σ_*i*_ are ordered so that 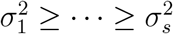 then it follows that near the maximum the likelihood is most sensitive when *θ*′_1_ is varied and least sensitive when *θ*′_s_ is. Moreover, σ_*i*_ is a measure of this sensitivity.

The theory of optimal experimental design is based on the idea of trying to make the σ_*i*_ decrease as slowly as possible so that the likelihood is as peaked as possible around the maximum, thus maximising the information content of the experimental sampling methods. Various criteria for experimental design have been proposed including D-optimality that maximises the determinant of the FIM and A-optimality that minimise the trace of the inverse of the FIM [6]. Diagonal elements of the inverse of FIM constitute a lower-bound for variance of any unbiased estimator of elements of *θ* (Cramer-Rao inequality). However, for the systems we consider here the a_*i*_ typically decrease very fast and there are many of them. Thus, in general, criteria based on a single number are more likely to be of less use than consideration of the set of σ_*i*_ as a whole.

Calculation of the FIM for stochastic systems using the LNA has been carried out in [22] but only for small systems and short times where the LNA is accurate. It is notable that the pcLNA enables one to do such sensitivity analysis for large systems and large times. As an example, we analyse the stochastic behaviour of the *Drosophila* circadian clock based on the limit distribution *P* (*<u>Q</u>*|*Q*_0_) when

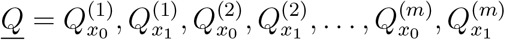

where *x*_0_ = *g*(*t*_0_) and *x*_1_ = *g*(*t*_1_) are chosen so that *t*_0_ is the time of the peak of *per* mRNA *M_P_*, and *t*_1_ is the peak of the nuclear complex of PER and TIM proteins *C_N_*. We compute the Fisher Information of the distribution *P*(*<u>Q</u>*|*Q*_0_) using the closed form expression (S1 Sect. 9.3) for this distribution. As we can see in Fig 10(A) the eigenvalues of the Fisher Information matrix decay exponentially, with a sharp decline followed by a slower decrease. This reveals that the influential directions in the parameter space of the system are much less than its total dimension and that only a few parameters appear to be most influential. The eigenvectors associated with the two largest eigenvalues of Fisher Information matrix (see Fig. 10(B)) have large entries only for the parameters *k_dn_* (PER-TIM complex nuclear degradation), *k_d_*(*per* mRNA linear degradation), *k*_2_ (PER-TIM complex transportation to cytosol), *v_st_* (*tim* mRNA transcription), *k_ip_* (*per* mRNA Hill coefficient) and *k_it_* (*tim* mRNA Hill coefficient).

**Fig 10.**
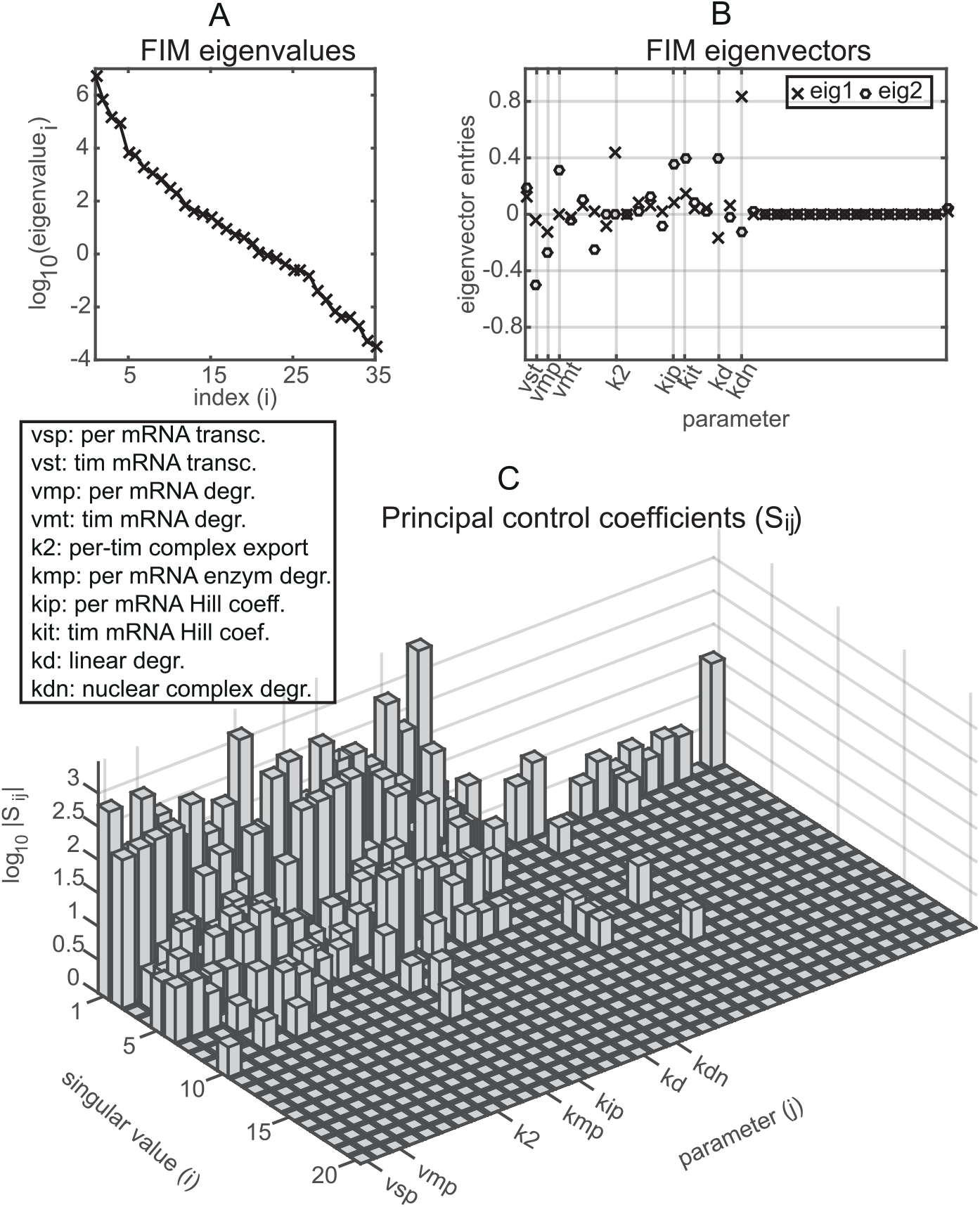
Fisher Information & Stochastic sensitivity analysis of the transversal distribution of the *Drosophila* circadian clock system at the times of the peaks of *per* mRNA and the nuclear complex of PER and TIM proteins. (A) The logarithm of the eigenvalues of the Fisher Information Matrix (FIM). (B) The entries/weights of the eigenvectors corresponding to the 2 largest eigenvalues of FIM (C) The largest principal control coefficients *S_ij_*. Small values are reduced to 0.

The exponential decrease of the eigenvalues is typical of tightly coupled deterministic systems [25–31], but has to our knowledge not been demonstrated before for stochastic systems. It has important consequences. For example, it tells us that only a few parameters can be estimated efficiently from time-series data unless the system is perturbed in some way to get complementary data and that there will be identifiability problems that can be analysed using the FIM. It can also be used to design experiments by considering the FIM of a combination of models including one for the proposed new experiment, choosing the new experiment so as to optimally alleviate the decline of the eigenvalues.

## Sensitivity analysis for stochastic systems

The fact that we can calculate the Fisher Information allows a new approach to sensitivity analysis for stochastic systems. Anderson [23] and Srivastava et al. [24] also perform sensitivity analysis for small stochastic systems (up to 4 species and 8 reactions) in which they calculate the dependence of certain summary functions or statistics at one or more times to individual parameters. Our approach is different in that we use the fact that our distributions of interest are MVN and measure the change in the distribution of the system state at any given set of phases without recourse to any summary function and, moreover, this change is calculated for any combination of parameter variations. A major difference is our use of SVD below to find a basis of mean-covariance space using the principal components that enables us to decompose these changes into different orthogonal directions that pick out the important and unimportant directions. The approach can also be formulated for the wider class of exponential families i.e. distributions that admit a representation of the form

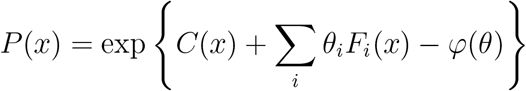

in terms of functions *C, F*_1_,… *F_m_* of the state variable *x* and a function *φ* of the parameters *θ*.

We consider a family of probability distributions *P* (*X, θ*) which we assume are MVN with mean *μ*(*θ*) and covariance matrix Σ(*θ*) depending on the parameters *θ*. We show that there is a natural matrix of sensitivities *S_ij_* associated with such a system. These are system-global in that they look at how all components of the systems change with parameters. They also have an intimate relationship with Fisher information. Note that these results are not restricted to the transversal distributions derived in previous sections but apply more generally to any MVN distribution with mean *μ* = *μ*(*θ*) and covariance matrix Σ = Σ (*θ*) parameterised by a *s*-dimensional vector *θ*, *s* ≥1.

As is well-known in Information Geometry, the set of multivariate normal distributions MVN^*n*^ on ℝ^*n*^ can be given the structure of a Riemannian manifold in which the Riemannian metric is given by the line element

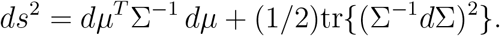

Points in MVN^*n*^ are denoted by Θ = (*μ*, Σ) where *μ* is the mean and Σ the covariance matrix. The corresponding inner product in the tangent space at Θ_0_ = (*μ*, Σ) is given by

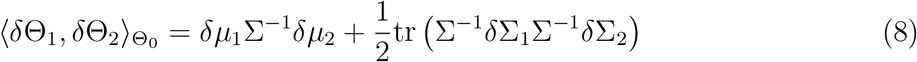

where δΘ_j_ = (δμ_j_, δΣ_j_), j = 1, 2.

In calculating the FIM we have to determine the partial derivatives *∂μ⁄∂θ_i_* and *∂*Σ⁄*∂θ_i_*. The derivative *M* of the mapping *θ*→ (*μ*(*θ*), Σ(*θ*)) at a parameter value *θ*_0_ is given by

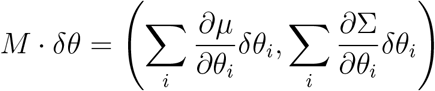

where the derivatives are calculated at *θ*_0_.

We can regard *M* as a linear mapping between the parameter space ℝ^*s*^ and MVN^*n*^ with the inner product given in equation (8). We can then prove (S1 Sect. 11) that we can find *s* orthonormal vectors *V_i_* spanning the parameter space ℝ^*s*^, *s* orthonormal vectors *U_i_* in the space MVN^*n*^ and numbers σ_1_ ≥*⋯* ≥ σ_*s*_ ≥0 such that

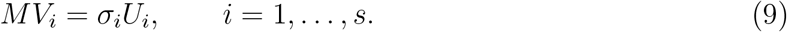

Note that the orthonormality of the *U_i_* is with respect to the inner product 〈·, ·〉_Θ_0__. The eigenvalues of the FIM *F* are the squares of the σ_*i*_ because with respect to the standard inner product on *θ*-space and〈·, ·〉_Θ_0__ on MVN^*n*^ the adjoint *M^*^* satisfies *M*M* = *F* (S1 Sect. 11).

If we let 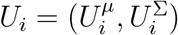 denote the decomposition of *U_i_* into *μ* and Σ components, then the following key property follows from (9): if *δ*θ is any change of parameters, the change in *μ* and Σ is given by

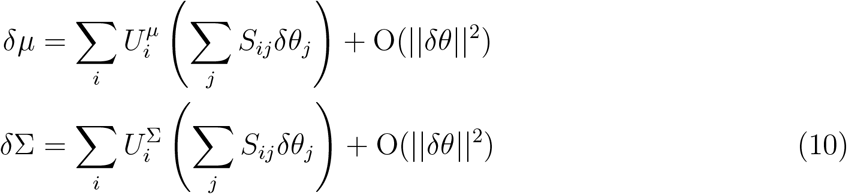

where *S_ij_* = *σ_i_V_ji_*.

One can define other sensitivities in a similar way but using a different orthogonal basis of MVN^*n*^, but the above *S_ij_* satisfy an important optimality condition explained in S1 Sect. 11 which asserts that the basis *U_i_* and the corresponding sensitivities *S_ij_* are optimal for capturing as much sensitivity as possible in the low order principal components *U_i_*.

In view of this we call the *S_ij_* the *principal control coefficients*. Note that the role of the *S_ij_* as sensitivities is seen from the following relation which follows from (10),

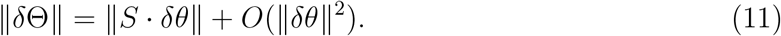

These sensitivities are relatively easy to calculate using the information in S1 Sect. 11. In Fig 10(C) we show the *S_ij_* for the transversal distribution of the *Drosophila* circadian clock at the times of the peak of *per* mRNA and the peak of the nuclear complex of PER and TIM proteins. As we can see, because *S_ij_* = *σ_i_V_ji_* the coefficients rapidly decrease with the singular values *σ_i_*, while a few parameters, similar to those with large eigenvector entries, have high coefficients.

## Calculating likelihoods via a pcLNA Kalman Filter

The likelihood function of a set of time-series observations of a system can be used for parameter estimation, hypothesis testing and other forms of statistical inference. For example, one may wishes to use the likelihood function to estimate parameters of a biological system. Although there is no elegant formula for

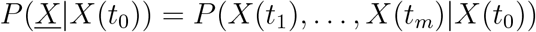

similar to that for *P*(*<u>Q</u>*|*X*(*t*_0_)) above, we can efficiently calculate it. To do this we derive a Kalman Filter for the pcLNA that is a modification of the Kalman Filter associated with the LNA [35]. This can be used to compute the likelihood function *L*(*θ*; *<u>X̂</u>*) of the system parameters *θ* with respect to observations *<u>X̂</u>* = (*X̂*(*t*_0_), *X̂*(*t*_1_),…, *X̂*(*t_N_*)) recorded at *N* times *t*_0_, *t*_1_,… *t_N_* that are noisy linear functions of the species concentrations. This is slightly more general than just calculating *P*(*<u>X</u>*|*X*(*t*_0_)) because we allow for a measurement equation. The Kalman filter can also be used for forward prediction.

We assume the measurement equation,

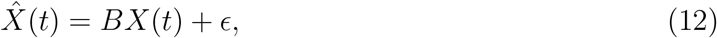

relating the observations *X̂*(*t*) to the state variables, *X*(*t*). Here *B* is a transformation matrix (often simply removing unobserved species or introducing unknown scalings) and ∊ = (∊_1_,…, ∊_*n*_) ~*MV N* (0, Σ_∊_) the observational error. The pcLNA likelihood can be decomposed as the product

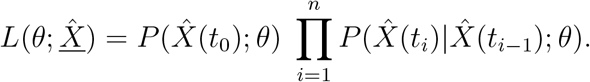

The pcLNA Kalman Filter algorithm, which we describe in more detail in S1 Sect. 15, uses a recursive algorithm for computing the terms in *L*(*θ*; *<u>X̂</u>*). The algorithm proceeds by iteration *i* = 1, 2,… and uses Bayes rule to derive the posterior distributions (*X*(*t_i-_*_1_)|*X̂*(*t_i-_*_1_)) and a phase correction to obtain *g*(*s_i_*-_1_) = *G_N_*(*μ^*^*(*t_i_*-_1_)) and the corrected noise distribution (κ(*s_i_*-_1_)*|X̂*(*t_i-_*_1_)). The LNA transition equation (S1 Eq. (4.2)) is then used to derive the distribution of (*ξ*(*t_i_*) |*X̂*(*t_i-_*_1_)) and the LNA ansantz equation (3) to obtain (*X*(*t_i_*) |*X̂*(*t_i_*-_1_)). The measurement equation (12) is finally used to obtain the (*i* + 1)th term of the likelihood function *P*(*X̂*(*t_i_*) |*X̂*(*t_i-_*_1_)) before proceeding to the next iteration. All the distributions obtained in this way are MVN with easily computable parameter values.

If the observations are recorded in short time intervals, the phase correction can be omitted in some steps, in which case the algorithm proceeds as in [35]. Computational methods such as those described in [32] and [35] can then be used to perform likelihood-based statistical inference.

## Methods

All computations have been carried out using MATLAB Release 2016b, The MathWorks, Inc., Natick, MA, USA. In particular, the empirical CDF plots, (q-q) plots, histograms, smooth probability density functions and KS distances are derived using the ecdf, qqplot, histogram, ksdensity, kstest functions of MATLAB and Statistics Toolbox. The computations for the SSA, tau-leap, integration of diffusion and pcLNA simulation algorithms, and the computation of Fisher Information and principal control coefficients for the sensitivity analysis were performed using the PeTTSy software which is discussed in S1 Sect. 14 and is freely available at http://www2.warwick.ac.uk/fac/sci/systemsbiology/research/software/. Further details concerning methods are given in S1 Sect. 16.

## 3 Discussion

We present a comprehensive treatment of stochastic modelling for large stochastic oscillatory systems. Practical algorithms for fast long-term simulation and likelihood-based statistical inference are provided along with the essential tools for a more analytical study of such systems.

There is considerable scope for future work in various directions. We expect that these results can be extended to a broader class of systems including those that are chaotic in the Ω → ∞ limit. Our approach should provide the opportunity to develop new methodology for parameter estimation, likelihood-based inference and experimental design in such systems. Finally, there is currently much interest in information transfer and decision-making in signaling systems and our methods provide new tools with which to tackle problems in this area.

If system biologists are to reliably use complex stochastic models to provide robust understanding it is crucial that there are analytical tools to enable a rigorous assessment of the quality and selection of these models and their fit to current biological knowledge and data. Our aim in this paper is to contribute to that but the results should be of much broader interest.

## Supporting Information

**S1 Appendix. Technical Details.** In this note we give further details about the mathematical underpinnings of the pcLNA methods discussed in the main paper.

**S2 Appendix. *Drosophila* circadian clock system.** In this note we give details about the *Drosophila* circadian clock and use this system to illustrate further the accuracy of distributions and simulations discussed in the main paper.

**S3 Appendix. Brusselator and NF-***κ***B systems.** In this note we give details about the deterministic and stochastic models of the Brusselator and NF-*κ* B systems and use them to illustrate further the results described in the main paper.

## Acknowledgments

This research was funded by the BBSRC Grant BB/K003097/1 (Systems Biology Analysis of Biological Timers and Inflammation). DAR was also supported by funding from the European Union Seventh Framework Programme (FP7/2007-2013) under grant agreement n° 305564.

## Supplementary information S1 for the paper Long-time analytic approximation of large stochastic oscillators: simulation, analysis and inference

> We refer to the paper “Long-time analytic approximation of large stochastic oscillators: simulation, analysis and inference” by **I**. In this note we give further details about the mathematical underpinnings of the pcLNA methods discussed in **I**.

### 1 Mathematical details

When we refer to a function, mapping or vectorfield as being smooth we mean that it has *C*^2^ dependence on the relevant variables. We are assuming throughout that the vector field *F* introduced in **I** is *C*^2^ and that the limit cycle γ given by *g*(*t*) is a stable elementary limit cycle in the sense we describe now. Here *x* ∈ ℝ^*n*^.

If *g*(*t*) is the periodic solution of interest (with period *τ*), consider the linear differential equation *dC/dt* = *J*(*t*) *C* where *J*(*t*) is the *n* × *n* Jacobian matrix whose *ij*th entry is ∂*F_i_/*∂*x_j_* evaluated at *g*(*t*). Let *C*(*s, t*) be the solution of this equation with initial condition *C*(*s, s*) = *I_n_* the identity *n* × *n* matrix. If *C*(*t*) = *C*(*s, t*) then *C*(*t*) has a representation of the form *C*(*t*) = *Z*(*t*)*e^R(t-s)^* where *Z*(*t* + *τ*) = *Z*(*t*)*e^Rτ^* (Theorem 6.1 of Chapter IV [5]). If λ_1_,…, λ_*n*_ are the eigenvalues of *R* then *e^t^*^λ1^,…, *e^t^*^λ1^ are the eigenvalues of *e^Rt^*. As such they are only determined modulo 2*π i/τ*. Modulo this indeterminacy, they are called the *characteristic exponents* of the limit cycle. The characteristic exponents are independent of the choice of the initial time *s*.

We assume that all limit cycles *γ* considered in this paper are *elementary and stable* in the following sense. In the case of a limit cycle *γ* of a free-running oscillator (i.e. when *F* does not depend directly upon *t*), we say that *γ* is *elementary* if *n* – 1 of the characteristic exponents have non-zero real parts and the remaining characteristic exponent is zero (modulo 2*π i/τ*). Such limit cycles are often called *hyperbolic*. If in addition the *n* – 1 non-zero characteristic exponents have negative real part then we say it is *stable and elementary*.

In the case of periodically forced oscillators the limiting differential equation is *ẋ* = *F*(*t, x*) and, in this case *γ* is elementary if all the characteristic exponents have non-zero real parts and it is stable if all have negative real parts.

A number of desirable properties flow from these conditions. Firstly, for elementary and stable limit cycles there is a neighbourhood of *γ* in ℝ^*n*^ such that any solution of *ẋ* = *F* (*t, x*) with its initial condition in the neighbourhood converges exponentially fast to *γ* (Theorem 11.1. Chap. IX [5]). Secondly, suppose *F* depends smoothly on parameters *θ* so that *F* = *F*_θ_. Then if *g_θ_*_0_(*t*) is a elementary limit cycle for *F* = *F_θ_*_0_, there is a unique family *g*_θ_ defined for *θ* near *θ*_0_ which depends smoothly upon *θ* such that *g_θ_* is an elementary limit cycle of *F* = *F_θ_* (Theorem 2.3, Chap. XII [5]).

Almost all the results described in the main paper that are for normal transversals *N*(*x*) hold for arbitrary transversals *𝒮*_x_. By a transversal section through *x* ∈ *γ* we mean a (*n* – 1)-dimensional linear hyperplane *𝒮_x_* containing *x* and transversal to the tangent vector, *F*(*x*), to *γ* at *x*. A particular example is the hyperplane normal to *γ* at *x*. A transversal system is a family *𝒮_g_*_(*t*)_ of transversal sections which vary smoothly with *t* in the sense that the unit normal vector to *𝒮_g_*_(*t*)_ varies smoothly with *t*. Throughout this document we consider this more general situation.

A transversal system defines a mapping *G* of a neighborhood of *γ* onto *γ* where if *X* ∈ *𝒮_x_* then *G*(*X*) = *x* ∈ *γ*. In cases where *X*(*s*) lies in more than one transversal sections, *𝒮_x_*_(*t′*)_, *t′* = *t*_1_, *t*_2_,…, then *G*(*X*(*s*)) = *x*(*t*) with *t* = min_*i*_ |*t_i_* – *s*| the closest time to *s*. We denote this mapping for the normal transversal system by *G_N_*.

An adapted coordinate system *𝒞_g_*_(*t*)_ at a point *g*(*t*) on *γ* is one determined by a set of orthonormal basis vectors *e*_1_(*t*),…, *e_n_*(*t*) with *e*_1_(*t*) the unit normal vector to *𝒮_g_*_(*t*)_ and the vectors *e*_2_(*t*),…, *e_n_*(*t*) forming an orthonormal basis of *𝒮_g_*_(*t*)_. If these are defined for *t* in some interval in ℝ then we always assume that the *e_i_*(*t*) have smooth (i.e. *C*^2^) dependence upon *t*. It is important that the coordinates are defined by an orthonormal basis in the original coordinates because this effectively preserves the covariance matrix in the sense that a covariance matrix *V* in the adapted coordinates is *UVU^T^* in the original coordinates with *U* a real orthogonal matrix. In particular, the eigenvalues are preserved.

### 2 The system size parameter Ω

It is very common to consider stochastic systems involving a system size parameter Ω which is a parameter that occurs in the intensities of the reactions *w_j_*(*Y*(*t*)) [8, 9]. The precise description of this parameter depends on the system and it governs the size of the fluctuations and therefore the size of the jumps. In population models it might be considered to be of the same order of magnitude as the total population size while in chemical systems a natural choice is to use molar concentrations and therefore regard Ω as Avogadro’s number in the appropriate molar units (e.g. nM^*–*1^) multiplied by the volume of the reacting solution in appropriate units (e.g. in litres (L)). In the *Drosophila* circadian clock model we consider it has units L/nM.

It can be interpreted as being the same order of magnitude as the mean size of the molecular populations. When the system and the systems size parameter satisfy the LNA Ansatz 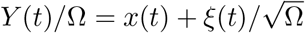 (see Sect. S4) then *Y*(*t*) = O(Ω) and the fluctuations are 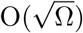.

In our examples, the rates *w_j_*(*Y*(*t*)) depend upon Ω as *w_j_*(*Y*) = Ω*u_j_*(*Y/* Ω) as used in the proof by Kurtz for the convergence of the LNA as Ω → ∞ [8]. For example, Hill functions in our rate equations have the form *w_H_* = *c* Ω*N ^h^/*(*N ^h^* + (*k* Ω)^*h*^) where *N* stands for the relevant population numbers and *c* and *k* are parameters. Therefore, if *n* = *N/* Ω, *w_H_* = *c* Ω*n^h^/*(*n^h^* + *k^h^*). Similarly, degradation and binding reaction rates have the form *w*_deg_ = *cN* and *w*_bind_ = *cN*_1_*N*_2_/Ω so that *w*_deg_ = *c* Ω*n* and *w*_bind_ = *c* Ω*n*_1_*n*_2_.

In a certain sense the system size parameter is just a mathematical convenience to enable the study of the dependence of stochastic fluctuations upon system size. Indeed, the methods we develop in this paper can and should be applied to systems that do not involve a system size parameter but then it will be necessary to ensure that the population sizes achieved in the given system are large enough.

### 3 Derivation of the classical deterministic equation

We provide a derivation of the macroscopic law of mass action that is similar to [1].

The state of the system at some time *t* is

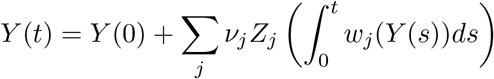

with *Z_j_* independent unit Poisson processes corresponding to the *j*-th reaction channel. The latter equation can be written in terms of *X* = *Y/* Ω as

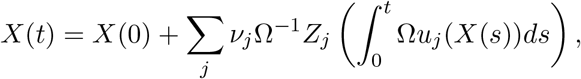

since *w_j_*(*Y*) = Ω*u_j_*(*X*) (see Section S2). Using the law of large numbers, as Ω → ∞, we have that

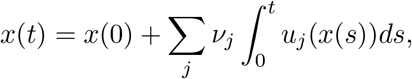

where here *x*(*t*) is the limit of *X*(*t*) for Ω → ∞. Consequently, *x*(*t*) satisfies the ordinary differential equation (ODE)

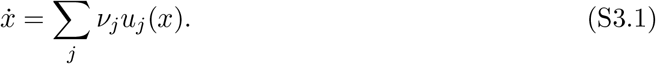

### 4 The Linear Noise Approximation (LNA)

The LNA is based on the Ansatz

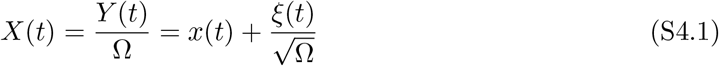

where *x*(*t*) is a solution of the limiting Ω → ∞ system in Eq. (S3.1) which we assume is a stable limit cycle of minimal period *τ >* 0 given by *x* = *g*(*t*), 0 ≤ *t* ≤ *τ*.

Recall the definition of the *n* × *n* matrices *C*(*s, t*) introduced in Section S1. In the LNA, the stochastic variable *ξ* in (S4.1) satisfies

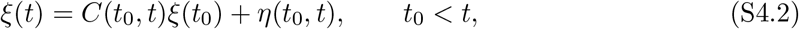

where *η*(*t*_0_, *t*) ~ MVN(0,*V*(*t*_0_, *t*)) is multivariate normal with mean 0 and covariance matrix

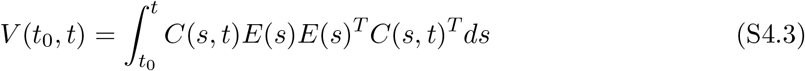

and *E*(*s*) the matrix product of the stoichiometry matrix (i.e. the matrix whose columns are the vectors *ν_j_* introduced in **I**) and the square root of the diagonal matrix with main diagonal the reaction rates *u_j_*(*x*(*s*)). The noise ξ (*t*) also satisfies the Stochastic Differential Equation,

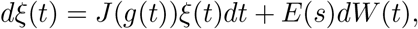

where *W*(*t*) is a *K*-dimensional Brownian motion (*K*: the number of possible reactions). Equation (S4.3) follows from this by using Itô’s change of variable formula and the Itô Isometry. The proof of convergence as Ω → ∞of the true distribution of *ξ* to a multivariate normal distribution with mean and covariance matrix given by (S4.2) and (S4.3) is contained in Chap. 8 of [8].

Henceforth, we write ζ ~ MVN(*m, S*) to mean that a random variable ζ is multivariate normal with mean *m* and covariance matrix *S*.

### 5 The LNA transversal distribution at a fixed time

We now suppose that our initial condition ξ (*t*_0_) is drawn from a MVN distribution *P*_0_ which is supported on the transversal section *S_g_*_(*t*0)_. This is denoted *ξ*(*t*_0_) ~ *P*_0_. For *t*_0_ *< t*_1_, we consider the distribution, *P*_LNA_(*X*(*t*_1_)*|X*(*t*_1_) ∈ *𝒮*_*g*(*t*1)_, and ξ (*t*_0_) ~ *P*_0_), of *X*(*t*_1_) conditional on *X*(*t*_1_) lying on the transversal sections *𝒮_g_*_(*t*1)_ and *i*(*t*_0_) ~ *P*_0_.

We have adapted coordinate systems (*y*_1_, **y**_2_) at *x*_1_ = *g*(*t*_1_) so that *𝒮_x_*_1_ is given by *y*_1_ = 0. We write the matrices *C*(*t*_0_, *t*_1_) and *V*(*t*_0_, *t*_1_) in the latter coordinate system as

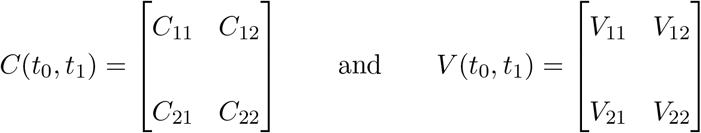

where 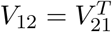 By *X*(*t*_0_) ∈ *𝒮_x_*_0_ we have that ξ (*t*_0_) = *κ*_0_. Then, under the LNA,

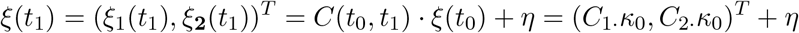

where *C*_1._ = [*C*_11_ *C*_12_], *C*_2·_ = [*C*_21_ *C*_22_] and *η* ~ MVN(0,*V*(*t*_0_, *t*_1_)). Consequently, ξ_**2**_(*t*_1_) conditional on ξ_1_(*t*_1_) = 0 is MVN with mean *Č*(*t*_0_, *t*_1_) *Čκ*_0_ and covariance matrix *V̌*(*t*_0_, *t*), where *Č*(*t*_0_, *t*_1_) = *C*_2*·*_ – *V*_21_*C*_1*·*_/*V*_11_ and *V̌*(*t*_0_, *t*_1_) = *V*_22_ – *V*_21_*V*_12_*/V*_11_. Thus if *κ*_0_ ~ MVN(*m*_0_, *S*_0_) and *κ*(*t*_1_) denotes ξ_**2**_(*t*_1_) conditional on *ξ*_1_(*t*_1_) = 0, then *κ*(*t*_1_) is MVN with mean and covariance matrix respectively given by

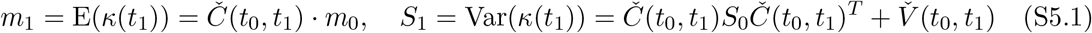

and *P*_LNA_(*X*(*t*_1_)*|X*(*t*_1_) ∈ *𝒮_x_*_1_, ξ (*t*_0_) ξ *MV N*(*m*_0_, *S*_0_)) is *MV N*(*μ*_1_, Σ_1_) with *μ*_1_ = Ω^−1/2^*m*_1_ and Σ_1_ = Ω^−1^*S*_1_.

### 6 Distribution of phase fluctuations *δt* and *g*(*t*) – *G_N_*(*X*(*t*))

In Sect. “Stochastic fluctuations in periods and timing” of **I**, we consider the distribution of the fluctuation δ*t* in the time taken for the lifted phase of a stochastic trajectory to go from a given phase ɸ_1_ to a greater one ɸ_2_. We showed that for moves from *ɸ*_1_ = 0 and *ɸ*_2_ = 2*πq* for *q* = 1,…, 8 the distribution of the fluctuations is approximately normal with variance increasing with *q* and that for *ɸ*_1_ = 2(*q* – 1)*π* and *ɸ*_2_ = 2*qπ* for *q* = 1,…, 8 the distributions are again approximately normal with approximately constant variance. We now consider an approximation of the distribution of the fluctuations under the LNA.

In this and the next section we will be considering how to approximate the distribution of first intersections with *𝒮_g_*_(*t*1)_ of stochastic trajectories *X*(*t*) that start in *𝒮_g_*_(*t*0)_. To do this we will consider LNA trajectories with initial condition a given MVN distribution *P*_0_ on *𝒮_g_*_(*t*0)_ and their first intersection at time *t* = *t*_1_ + *δt* with *𝒮_g_*_(*t*1)_. In this section we show that the distribution of the variation in time, b*t*, is approximately normal and calculate approximations of the mean and variance. Thus, we consider the distribution under the LNA of ξ(*t*) such that *X*(*t*) = *g*(*t*) + Ω^*-*1/2^ ξ(*t*) ∈ *𝒮_g_*_(*t*)_ for *t*_0_ ≤*t < t*_1_ + *τ* /2. We denote this distribution by *P*_LNA_(ξ(*t*)*|X*(*t*) ∈ *𝒮_g_*_(*t*1)_).

To calculate the statistics of the variation in time, *δt*, for the distribution of the *r*th intersection with *𝒮_g_*_(*t*1)_ of stochastic trajectories *X*(*t*) that start in *𝒮*_*g*(*t*_0_)_ one decomposes the trajectory into the part from *𝒮_g_*_(*t*_0_)_ to its first intersection with *𝒮_g_*_(*t*_1_)_ and then the (*r* – 1) passes from *𝒮_g_*_(*t*_1_)_ to itself and uses the fact that the total variation is the sum of the individual ones which are independent random variables. Since we show below that the individual variation are approximately normal it follows that the total is. Therefore in this section we now focus just on first intersections.

We assume that the initial conditions *ξ*(*t*_0_) have a MVN distribution *P*_0_ on *𝒮_g_*_(*t*_0_)_ with mean 0 and covariance matrix *V*_0_. We fix adapted coordinate systems at each *g*(*t*). Then *ξ*(*t*_1_ + *δt*) has mean 0 and covariance matrix *V̌*(*t*_0_, *t*_1_ + *δt*) = *C*(*t*_0_, *t*_1_ + *δt*)*V*_0_*C*(*t*_0_, *t*_1_ + *δt*)^*T*^ + *V*(*t*_0_, *t*_1_ + *δt*) where *V*(*t*_0_, *t*_1_ + *δt*) is given by (S4.3).

If *ξ*(*t*_1_ + *δt*) is such that *X*(*t*_1_ + *δt*) ∈ *𝒮_g_*_(*t*_1_)_ then *G*(*X*(*t*_1_ + *δt*)) = *g*(*t*_1_) and therefore 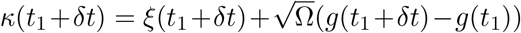. If *<u>n</u>*(*t*) denotes the unit vector *F*(*g*(*t*))/||*F*(*g*(*t*))||, since *κ*(*t*_1_ + *δt*) is orthogonal to *<u>n</u>*(*t*_1_), we have that

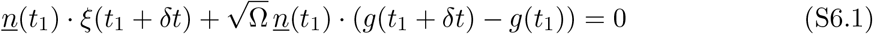

But since *ġ*(*t*) = *F*(*g*(*t*)) and the time derivative of *F*(*g*(*t*)) is *J*(*g*(*t*)) ·*F*(*g*(*t*)) it follows that, up to terms that are O(*|δt|*^3^),

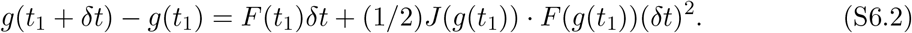

Now choose adapted coordinates (*x*_1_,…|, *x_n_*) at *g*(*t*_1_) so that *F*(*g*(*t*_1_)) = (α, 0,…, 0) and *<u>n</u>*(*t*_1_) = (1, 0,…, 0). Then, *<u>n</u>*(*t*_1_) · (*J*(*g*(*t*_1_)) · *F*(*g*(*t*_1_))) = α*J*_11_ where *J*_11_ is the first entry of *J*(*g*(*t*_1_)). Thus using (S6.1),

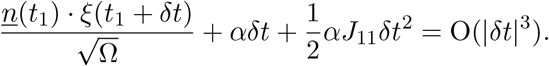

Since all terms but one are independent of Ω and since Ω(*t*_1_ + *δt*) has finite variance, it follows that the standard deviation of b*t* is O(Ω^−1/2^). Thus the rhs *η* is a random variable depending stochastically only on *δt* which is independent of Ω and has a standard deviation and mean that is O(Ω^−3/2^). Letting *κ* = *αJ*_11_/2 and 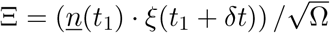, completing the square of the lhs gives 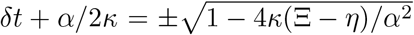 where *η* is the above O(*|δt|*^3^) term. Using 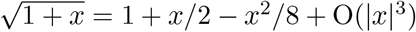 and ignoring the negative root gives,

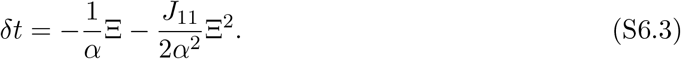

Note that *<u>n</u>*(*t*_1_)·ξ (*t*_1_ + *δt*) differs from *<u>n</u>*(*t*_1_) · ξ(*t*_1_) by an amount that is O(|*δt*|^2^) (since the base point moves a distance that is O(*|δt|*) around the limit cycle) and the mean of *ξ*(*t*_1_ + *δt*) given b*t* is *C*(*t*_0_, *t*_1_ + *δt*) *E*_*P*_0__(ξ (*t*_0_)) which is 0 if the mean *E_P_*_0_(*ξ*(*t*_0_)) of *P*_0_ is 0. Thus, the mean of Ξ is 0 and, up to an error that is O(Ω^*-*3/2^), the mean of the last term in (S6.3) is (*J*_11_/2*a*^2^)Var(Ξ) which is O(Ω^−1^) and the variance of this term is O(Ω^−2^).

The variance of *δt* is,

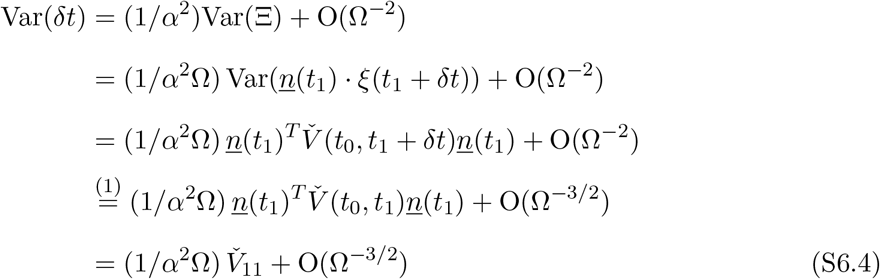

where *V̌*_11_ is the upper left entry of the covariance matrix *V̌*(*t*_0_, *t*_1_) written in the given adapted coordinates. The equality marked 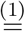 follows from the fact that *V̌*(*t_0_,t_1_*+*δt*) = *V̌*(*t_0_,t_1_*)+O(|*δt*|).

Since Ξ in (S6.3) has a MVN distribution, it follows that b*t* is approximately normal but is biased by the (*J*_11_/2*α*^2^) Ξ^2^ term which is O(Ω^*-*1^) and hence δ*t* has mean O(Ω^*-*1^) and variance (1/α^2^ Ω) *V*_11_ + O(Ω^−3/2^). For the *Drosophila* clock and system sizes Ω = 300, 500 and 1000, the standard deviation of *δt* for completing one round of the limit cycle under the above LNA formula is respectively 0.7218, 0.5591 and 0.3954 whereas the empirical standard deviation of δ*t* derived from the SSA is 0.7333, 0.5490 and 0.3862.

### 7 The approximation of the LNA transversal distributions

Recall the definition of *P*_LNA_(*ξ*(*t*)|*X*(*t*) ∈ *𝒮_g_*_(*t*1)_) in Sect. S6. In this section we consider how well this is approximated by *P*_LNA_(*ξ*(*t*_1_)|*X*(*t*_1_) ∈ *𝒮_g_*_(*t*1)_) i.e. by constraining time *t* to be the fixed time *t*_1_. We show that the variance Var_*δt*_(*X*(*t*_1_ + *δt*)) of *X*(*t*) = *X*(*t*_1_ + *δt*) conditional on *X*(*t*) ∈ *𝒮_g_*_(*t*_1__) equals the variance of *X*(*t*_1_) conditional on *X*(*t*_1_) ∈ *𝒮_g_*_(*t*_1__) up to a term that is O(Ω^*-*1^). The latter conditional variance is relatively easily calculated using the results in Sect. S5.

Throughout this section for a given *ξ*(*t*), *X*(*t*) denotes *g*(*t*) + Ω^−1/2^⇠(*t*) ∈ *𝒮_g_*_(*t_1_*)_. We first consider the variance of *X*(*t*_1_ + *δt*) conditional on a fixed value of δ*t* and *X*(*t*_1_ + *δt*) ∈ *𝒮_g_*_(*t*_1_)_. We omit the conditioning on *X*(*t*_1_ + *δt*) ∈ *S_g_*_(*t*_1_)_ from the notation in the rest of this section.

The corresponding covariance matrix Var(*X*(*t*_1_ + *δt*)*| δt*) has a Taylor expansion

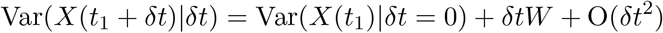

where the symmetric matrix *W* is easily calculated.

Note that *E*_*δt*_ [*δtW*] = O(Ω^*-*1^) since the mean of *δt* has this order (see Sect. S6). Consequently,

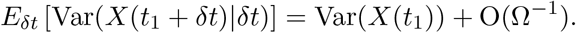

We now consider Var_*δt*_(*E*(*X*(*t*_1_ + *δt*)|*δt*)). The mean *E*(*X*(*t*_1_ + *δt*)*|*δ*t*) = *g*(*t*_1_ + *δt*) and therefore from (S6.2) and (S6.4), Var_δ*t*_(*E*(*X*(*t*_1_ + *δt*)|*δt*)) = *O*(Ω^−1^). Hence we have the result because, by the Law of Total Variation,

> Var(*X*(*t*_1_ + *δt*)) = *E*(Var(*X*(*t*_1_ + *δt*)|*δt*) + Var(*E*(*X*(*t*_1_ + *δt*)*| δt*)) = Var(*X*(*t*_1_)) + O(Ω^*-*1^).

### 8 Proof of convergence for LNA transversal distributions

#### 8.1 Free-running oscillators

We show that for any transversal *𝒮* = *𝒮_g_*_(*t*)_ to *γ* there is a unique MVN distribution 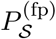 on *S* with the property that under the pcLNA, 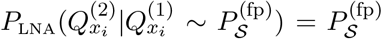. To do this we study the limit of 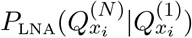 as *N*→ ∞ because this is such a fixed point.

We start by considering the case where *𝒮* is the normal hypersection and fix adapted coordinates (*x*_1_, **x**_2_) in which the transversal section *𝒮* is given by *x*_1_ = 0.

By periodicity the covariance matrix of *P*_LNA_(ξ (*t* + *n τ*)*|X*(0) = *g*(*t*)) = *P*_LNA_(ξ (0)*|X*(*t* – *n τ*) = *g*(*t*)) is *V*(*t* – *N τ, t*) (see (S4.3)). We have,

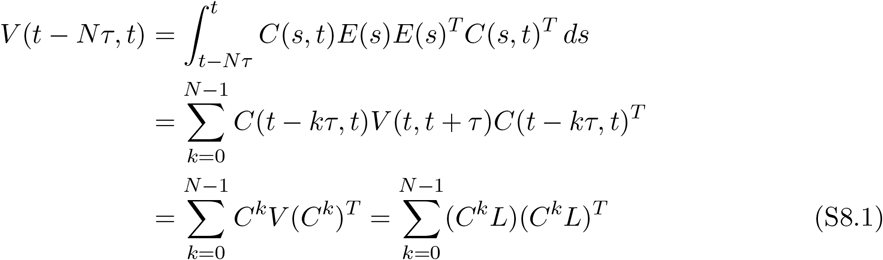

where C = C(*t, t*+ *τ*) and *V* = (*t, t*+ *τ*) = 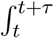 *C*(*s*, *t*)*E*(*s*)*E*(*s*)^T^*C*(*s*, *t*)^T^ *ds* and *V* = *LL^T^* is the Cholesky decomposition of the positive definite matrix *V*.

In what follows we utilise the vec operator and the Kronecker product ⊗. All the facts about these that are used here can be found in [7] Chapter 4. A basic equality is that vec(*AXB*) = (*B^T^* ⊗ *A*)vec(*X*). Therefore, since vec(*C^k^V*(*C^k^*)^*T*^) = (*C^k^* ⊗ *C^k^*)vec(*V*) = (*C* ⊗ *C*)^*k*^vec(*V*),

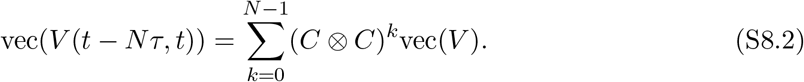

Our adapted coordinates (*x*_1_, **x**_2_) = (*x*_1_, *x*_1_,…, *x_n_*) at *g*(*t*) are determined by the the unit vectors *e_i_*(*t*) (see Sect. S5) where *e*_1_(*t*) = *F*(*g*(*t*))/||*F*(*g*(*t*))|| tangent to the limit cycle. Since *C e*_1_(*t*) = *e*_1_(*t*) it follows that

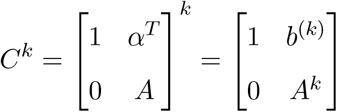

where *A* is an (*n* − 1) × (*n* − 1) matrix, *α^T^* is a *n* × 1 vector and *b^(k)^* = *α^T^* + *α^T^ A* ⋯ + *α^T^ A*^*k*-1^.

Since *γ* is elementary and stable, 1 is a simple eigenvalue of *C*, and therefore all eigenvalues λ of *A* have |λ| *<* 1 and therefore *b*^(*k*)^ converges to *b*^(∞)^ = *α^T^*(*I* – *A*)^−1^. Moreover, all eigenvalues λ≠ 1 of *C^k^* (i.e. all eigenvalues of *A^k^*) have modulus |λ| < *cv^k^* where 0 < *v* < 1 and *c* > 0 are constants independent of *k*.

It follows that **C**^(*k*)^ = (*C* ⊗ *C*)^*k*^ = *C^k^* ⊗ *C^k^* has the form

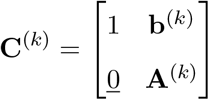

where **b**^(*k*)^ is 1 × (*n*^2^ – 1) and **A**^(*k*)^ is (*n*^2^ – 1) × (*n*^2^ – 1). The eigenvalues of *C^k^* ⊗ *C^k^* are all of the form λ*μ* where λ and *μ* are eigenvalues of *C^k^*. Thus 1 is a simple eigenvalue and all other eigenvalues λ have |λ| *< cv^k^* where 0 < *v* < 1 and *c* are constants independent of *k*. Consequently, all eigenvalues of **A**^(*k*)^ have the latter property.

Moreover, if 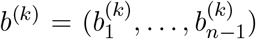 then the top row of **C**^(*k*)^ is obtained by concatenating the row vectors 1 ⊗ [1 *b*^(*k*)^] and *b*^(*k*)^ ⊗ [1 *b*^(*k*)^] for *i* = 1,…, *n* – 1, and therefore the entries in this row converge exponentially fast to finite limits. Consequently, if we write vec(*V*) as α_1_*E*_1_ + *Z* where *E*_1_ = (1, 0,…, 0)^*T*^ and *Z* = (0, *<u>z</u>*)^*T*^ is a vector perpendicular to *E*_1_,

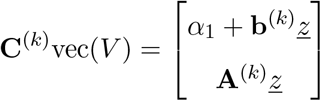

and the top entry converges exponentially fast to a limit *α*_1_ + **b**^(∞)^*<u>z</u>* > 0 i.e. |**b**^(*k*)^ – **b**^(^∞^)^| *≤ cv^k^* where 0 < *v* < 1 and *c* are constants independent of *k*.

It follows immediately by (S8.2) that if we write *V*(*t* – *N τ, t*) in terms of the above coordinates (*x*_1_, **x**_2_) = (*x*_1_, *x*_1_,…, *x_n_*) as

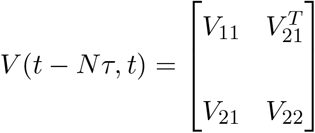

then, as *N* → ∞, while *V*_21_ and *V*_22_ have finite limits, *V*_11_ →+∞. Using the Schur complement, the precision matrix of *P* ^(*N*)^ = *P*_LNA_(ξ(0)*|X*(*t* – *N τ*) = *g*(*t*)) is

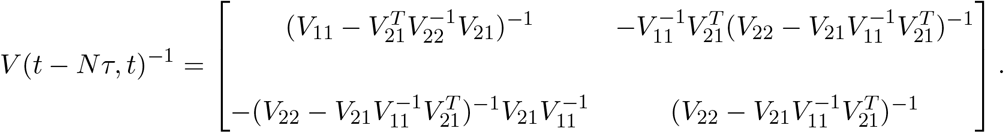

This has a well-defined limit as *N*→ ∞ where, since *V*_11_ → ∞, all entries but (*V*(*t*-*N τ, t*)^−1^)_22_ converge to zero and (*V*(*t* – *N τ, t*)^−1^)_22_ has a finite non-zero limit equal to (lim_*n* → ∞_ *V*_22_)^−1^. Thus, if *P̌*^(*N*)^ is the distribution given by conditioning *P* ^(*N*)^ on *x*_1_ = 0 then the precision matrix of *P̌*^(*N*)^ is 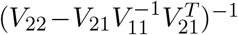. Consequently, the covariance matrix of *P̌*^(*N*)^ is 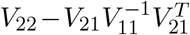 which in the limit *N* → ∞ equals 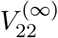 = lim_*n* → ∞_ *V*_22_. We thus deduce that the conditional distribution converges as *n* → ∞ to a MVN distribution *P* with this covariance matrix. Since *P* is derived via the limiting process as *N* → ∞, it is clear that 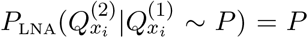 so 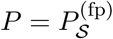

#### 8.2 Extending this result to arbitrary transversals

Any transversal hyperplane *𝒮*is of the form *W𝒮_N_* where *𝒮_N_* is the normal hyperplane and *W* is a *n* × *n* orthogonal matrix with *W*_11_ *>* 0. This latter condition ensures that *𝒮* is transversal to the circle. Thus if **z** = (*z*_1_, **z**_2_) = *W* **y** then in the **z** coordinate system *𝒮* is given by *z*_1_ = 0. In this **z** coordinate system, the precision matrix of 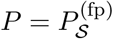 is given by *WV*(*t* – *n τ, t*)^*-*1^*W ^T^*.

However, from (8.1)

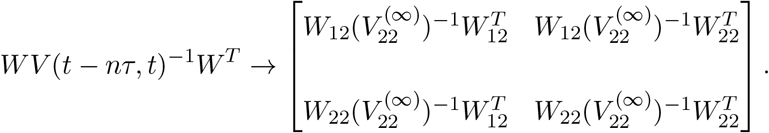

Therefore, in the limit, **z**_2_ |(*z*_1_ = 0) has covariance matrix 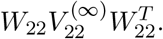.

#### 8.3 Alternative geometrical proof that convergence on *𝒮_N_* implies convergence on other transversals *𝒮*

Let *𝒮_N_* be given by *y*_1_ = 0 in the **y** coordinate system. Consider the ellipsoid *ε* given by 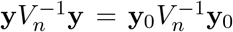 where **y**_0_ ∈ *𝒮_N_*. If *p* ∈ *𝒮_N_* consider *p* + δ*e*_1_ where *δ* ∈ ℝ and *e*_1_ is the vector (1, 0,…, 0) in the **y** system. Clearly there is a unique *δ* such that *p* + *δe*_1_∈ *𝒮*. Denote this point by *ϕ*(*p*). Then *ϕ* is linear. Consider 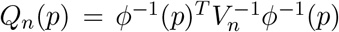. By the above result for *𝒮_N_*, on *𝒮_N_*, *Q_n_* converges to a *Q*_∞,*N*_. In the region |*y*_1_| < *r*, the ellipsoids given by *Q_n_*(*p*) = ε> 0 converge to cylinders of the form {(*y*_1_, **y**_2_): *Q*_∞*N*_ (**y**_2_) = ε}. Therefore, *Q_n_*(*p*) → *Q*_∞_(*p*) where *Q*_∞_(*p*) = *Q*_∞,*N*_ (**y**_2_) if *p* has coordinate (*y*_1_, **y**_2_).

**Extension to** 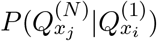 *x_j_* = *g(t_j_)*, *x_i_* = *g(t_i_)*, *t_j_* ≥ *t_i_*. The result is easily derived using that *V*(*t_i_, t_j_* + *N τ*) = *C*(*t_i_, t_j_*)*V*(*t_i_, t_i_* + *N τ*)*C*(*t_i_, t_j_*)^*T*^ + *V*(*t_i_, t_j_*) which implies existence of the limit of *V*^−1^(*t_i_, t_j_* + *N τ*), as *N* → ∞, if lim_*N*→ ∞_ V^−1^(*t_i_, t_i_* + *N τ*) exists.

#### 8.4 Entrained forced oscillators

The above proof of Section S8.1 has to be slightly modified as follows. In this case all eigenvalues of *C*(*s, t*) have | *λ*| *<* 1 and therefore all eigenvalues > of *C^k^* ⊗*C^k^* have | *λ*| < *cv^k^* where 0 < *v* < 1 and *c* are constants independent of *k*. Consequently, using (S8.2), all entries of *V*(*t* – *Nτ, t*) converge to a finite limit as *N* → ∞.

This shows that in contradistinction to free-running oscillators, for entrained forced oscillators, *P*_LNA_(*X*(*Nτ*)|*X*(0)) converges as *N* → ∞. However, it does not mean that this is a good approximation to *P*(*X*(*Nτ*)|*X*(0)) for an exact simulation. As we show in the simulation study displayed in S2 Fig. 9, these two distributions appear to remain relatively close to each other as N increases. In these simulations, *P*(*X*(*Nτ*)|*X*(0)) tends to wrap itself around the limit cycle while *P*_LNA_(*X*(*Nτ*)|*X*(0)) remains a bounded MVN. On the other hand, the conditional distributions 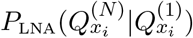do appear to remain accurate as N → ∞.

### 9 Analytic expression for pcLNA transversal distributions

To calculate an analytic expression for

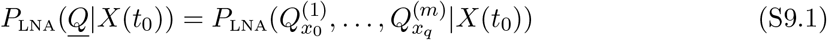

we need to introduce some new matrices. At each of the points *x_k_* we fix a coordinate system *C_x_k__* defined by a set of orthonormal vectors *e*_1_(*x_k_*), …, *e*_n_(*x_k_*) where *e*_1_(*x_k_*) is normal to the hyperplane *𝒮_x_k__*. We write coordinates in *𝒞_x_k__* in the form (*y*_1_, ***y***_2_) where *y*_1_ ∈ ℝ and ***y***_2_ ∈ ℝ^n_1^ and ***y***_2_ are coordinates on *𝒮_x_k__*. The matrices *C(s, t)* and *V (s, t)* in this coordinate systems are

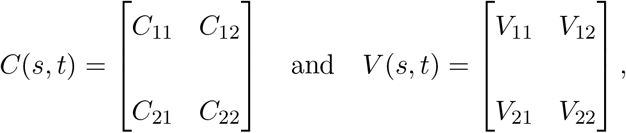

where 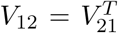. Let 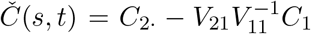 and 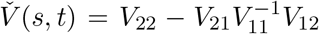. Note that *V̌(s, t)* is the Schur complement of *V_22_* in *V(s, t)* so if (*y_1_, **y**_2_*) is MVN with covariance *V(s, t)* then the distribution of *y_2_* conditional on y1 having a specific value has covariance *V̌(s, t)*.

Firstly, we consider 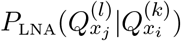 where either *k* < *l* or *k* = *l* and *i* < *j*. In Section S7 we showed that this is very well approximated by a MVN distribution with mean 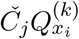 and covariance *V̌_j_* where

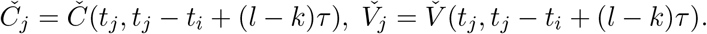

We also proved in Section 8 that as *l*−*k*→ ∞, 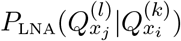 converges to a limit *P_x_j__* which is independent of 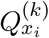 and does this exponentially fast in the sense that the mean and covariance 13 of 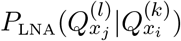 converge exponentially fast to their limiting values.

#### 9.1 Fixed point distribution

The special case where the transitions are returns to the same transversal sections (*i* = *j*) is especially interesting. Clearly, 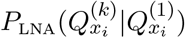 converges to the same limit *P_x_i__* as above and this distribution is a fixed point in the sense that the distribution 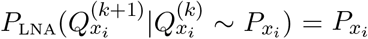. We therefore denote it 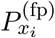. Because of the aforementioned convergence the transversal distribution has the following important property

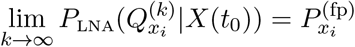

for any initial condition *X*(*t*_0_). That is, the distribution of the system at any transversal section converges to a limiting multivariate normal distribution.

This distribution can be easily calculated numerically. To derive its mean and covariance matrix one solves the equations *m* = *Č*_i_*m* and 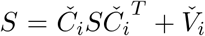, respectively. The latter can be solved by using the fact that 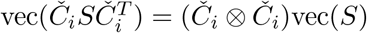 which implies that

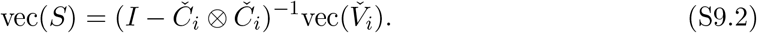

Here vec(*S*) is the vector obtained by stacking all the columns of *S* on top of each other and ⊗ is the Kronecker or tensor product (see e.g. Chapter 4 of [7]).

For the circadian clock the convergence is fast: the *L*^2^-norm of the difference between the covariance matrices of the limiting distribution, 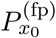 and 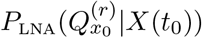 for *r* = 1, 2,…, 5 is respectively (1250, 70, 3.9, 0.2, 0.1) 10^−3^.

#### 9.2 The joint distribution of multiple transversals

We now consider the distribution of the sequence of transitions between transversal sections of different phases *P_LAN_*(*<u>Q</u>*|*X*(*t_0_*)). We relabel 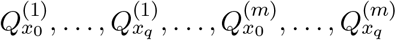 as *Q_1_*,…, *Q_N_* where *N* = *m*(*q* + 1). To each 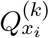 there is a corresponding time *t_i_* + (*k* −1)*τ*. We label these times in increasing order as *T*_1_,…*T_N_* so that *T_n_* corresponds to *Q_n_*. Let *T*_0_ = *t*_0_. With this notation if follows from the above that *P*_LNA_(*Q_i_*_+1_*|Q_i_*) is approximately MVN with mean *Č_i_Q_i_* and covariance *V̌*_*i*_ where *Č*_*i*_ = *Č*(*T_i_, T_i+1_*) and *V̌*_*i*_ = *V̌*(*T_i_, T_i_*_+1_). Consequently, if *Q_i_* has covariance matrix *Š*_*i*_ then *Q_i_*_+1_ has covariance matrix given by 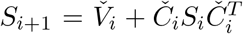. As explained in the next section, it follows that *P*_LNA_(*<u>Q</u>*|*Q*_0_) is MVN with mean

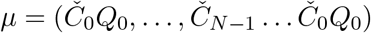

and covariance ∑ where the precision matrix ∑^−1^ is a tridiagonal matrix with only non-zero entries, the main diagonal 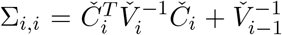 for *i* = 1, 2,…, *N* − 1 and 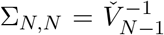, upper diagonal 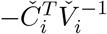 and lower diagonal 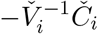.

#### 9.3 Precision matrix for *P*(*<u>Q</u>*|*X*(*t*_0_))

It is clear from the above that the mean of *P*_LNA_(*Q*_*m*+1_|*Q_m_*) is *Č_m_Q_m_* and that *Š*_*i*+1_ = *V̌*_*i*_ + 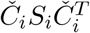 which imply

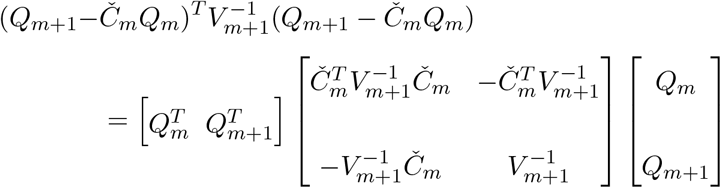

and the formula for the precision matrix for *P*(*<u>Q</u>*|*X*(*t*_0_)) follows from this using *P*_LNA_(*<u>Q</u>*|*X*(*t*_0_)) = 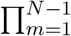 *P*_LNA_(*Q*_*m*+1_|*Q*_*m*_).

### 10 Other transversal sections

For each state variable *x_i_* two interesting transversals to *γ* are given by the submanifolds on which *ẋ_i_*(*t*) = 0, i.e. where the maximum or minimum of *x_i_*(*t*) occurs at time *t*. These are both given by *F_i_*(*x*) = 0 and their tangent spaces consist of those vectors *ν* such that ∂*F_i_/*∂*x*(*g*(*t*))·*ν* = 0. A continuous time trajectory intersects these transversal sections at times when the *i*th variable is at its maxima and minima. Thus *<u>Q</u>* gives the state of the system at their (large volume limit) maxima and minima.

Another very interesting transversal system is that given by the isochrons of *γ*. These are defined as follows. Let Φ(*t, x*) be the flow associated to *F*. For *x* ∈ *γ* the stable manifold *W ^s^*(*x*) of *x* is the set of points *y* such that Φ(*t, y*) – Φ(*t, x*) → 0 as *t* → ∞. If *γ* is an elementary and stable limit cycle as defined in Sect. S1, then, in a neighbourhood of *γ*, these stable manifolds are codimension one submanifolds that fill the neighbourhood. This follows from the Stable Manifold Theorem [6] and is explained in [4]. They are called isochrons because two points in the same isochron asymptotically have the same phase. They are invariant manifolds in the sense that *W^s^*(Φ(*t, x*)) = Φ(*W^s^*(*x*), *t*) i.e. at time *t* the flow caries the stable manifold of *x* exactly onto the stable manifold at Φ(*t, x*). In particular, if τ is the period then Φ(*τ, ⋅*) maps *W ^s^*(*x*) to *W ^s^*(*x*) and therefore, using the notation above, if *x* = *g*(*t*), the linear map given by *C*(*t, t* + *τ*) maps the (*n* − 1)-dimensional tangent space *V_x_* of *W ^s^*(*x*) to itself. By using the fact that *V_x_* is transverse to *γ*, or directly from the Invariant Manifold Theorem, it follows that *V_x_* is spanned by the generalised eigenspace of *C*(*t, t* + *τ*) corresponding to the eigenvalues λ with |λ| < 1. Consequently, a very interesting transversal system is that where *𝒮_g_*_(*t*)_ = *V_g_*_(*t*)_ is the tangent space to the isochron through *g*(*t*). By the invariance we have *C*_12_ = 0 so that *Č* = *C*_22_.

A final interesting transversal system is based on the *V* matrices. For an arbitrary point on the limit cycle *x_t_* = *g*(*t*), the matrix *V* = *V*(*t, t* + *τ*) is symmetric positive definite and therefore its orthonormal eigenvectors *ẽ*_*i*_, *i* = 1,…, *n* span ℝ^*N*^. The eigenvector corresponding to the largest eigenvalue of *V* is approximately equal to the unit tangent. We can define the transversal section as the hyperplane spanned by *ẽ*_2_,…, *ẽ*_*n*_. The advantage of this transversal system is that the largest variability direction is eliminated in the transversal distributions.

### 11 Details behind the sensitivity analysis

If *P* is MVN with mean and covariance *μ* = *μ*(*θ*) and ∑ = ∑(*θ*) then the entries of the FIM are given by

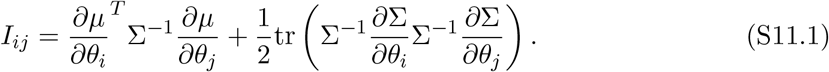

As above we consider a family of probability distributions *P*(*X, θ*) depending on parameters in *θ*. We assume these are MVN with mean *μ*(*θ*) and covariance matrix ∑(*θ*). In calculating the FIM we have to determine the partial derivatives ∂*μ/*∂*θ_i_* and ∂∑/∂*θ_i_*. The derivative *M* of the mapping *θ* → (*μ*(*θ*), ∑(*θ*)) at a parameter value *θ*_0_ is given by

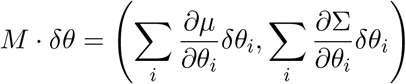

where the derivatives are calculated at *θ*_0_.

As is well-known in Information Geometry the set of multivariate normal distributions MVN^*n*^ on ℝ^*n*^ can be given the structure of a Riemannian manifold in which the Riemannian metric is given the line element *ds*^2^ = *dμ^T^* ∑^−1^ *dμ* + (1/2)tr{(∑^−1^*d*∑)^2^}. Points in this are denoted by Θ = (*μ*, ∑) where *μ* is the mean and ∑ the covariance matrix. The inner product in the tangent space at Θ_0_ = (*μ*(*θ*_0_), ∑(*θ*_0_)) is given by

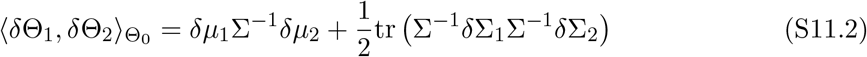

where Θ_0_ = (*μ*, ∑) and bΘ_*j*_ = (*δμ_j_, δ*∑_*j*_), *j* = 1, 2.

The derivative *M* above at *θ*_0_ is a linear map of the parameter space into MVN^*n*^ and therefore (taking the standard inner product 〈*δθ*_1_,*δθ*_1_〉 = 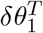·*δθ*_2_ on the parameter space) has an adjoint *M ^*^* such that 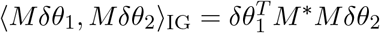. In fact, the adjoint operator *M ^*^* to *M* is given by

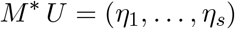

where

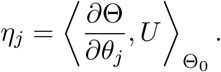

By (S11.1) and (S11.2), 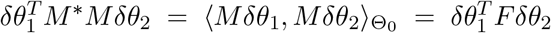 and therefore it follows that

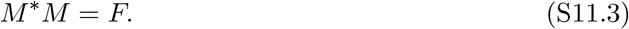

By applying Singular Value Decomposition appropriately (see below) we can find orthonormal vectors *V_i_* spanning the parameter space ℝ^*s*^, *s* orthonormal vectors *U_i_* in the space MVN^*n*^ and numbers σ_1_ ≥…≥σ_*s*_ such that

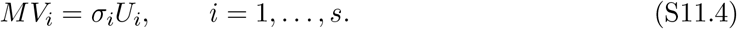

Note that the orthonormality of the *U_i_* is with respect to the inner product h 〈*·,·〉*Θ_0_ It follows from (S11.4) that *M ^*^MV_i_ ·V_j_* = σ_*i*_σ*_j_δ_ij_* and therefore that 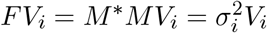

We regard a *s* × *s* matrix *S′_ij_* as a sensitivity matrix if there is a corresponding basis for 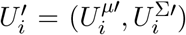 such that for any change of parameters *δθ*, the change in *μ* and ∑ is given by

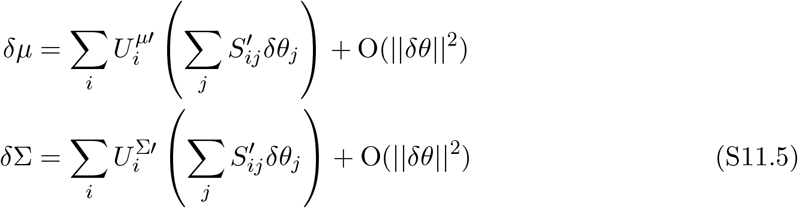

Note that the role of the *S′_ij_* as sensitivities is seen from the following relation which follows from (S11.5),

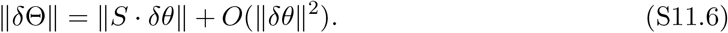

With this definition *S_ij_* = *σ_i_V_ji_* is a sensitivity matrix (with 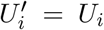 which has the following optimality property: for all *k* = 1,…, *s*

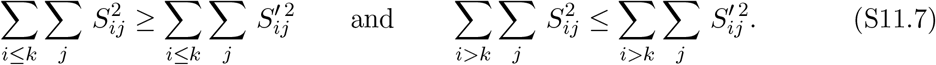

This tells us that the basis *U_i_* and the corresponding sensitivities *S_ij_* are optimal for capturing as much sensitivity as possible in the low order principal components. Note also that (i) the rows 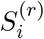 of *S_ij_* are orthogonal, and (ii) 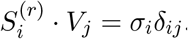

#### 11.1 Deducing equation (S11.4) from SVD

We can write the points of MVN^*n*^ in Cartesian coordinates as (*μ*, vec(∑)) and a straightforward calculation shows that in this representation the inner product 〈(*μ*_1_, ∑_1_), (*μ*_2_, ∑_2_)〉_(*μ*,∑)_ is given by

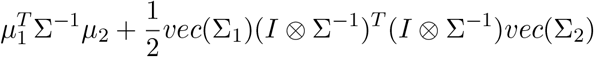

which can be written as (*μ*_1_, *vec*(∑_1_))*𝓕*(*μ*_1_, *vec*(∑_2_)) where *𝓕* the (*n* + *n*^2^) × (*n* + *n*^2^) square matrix

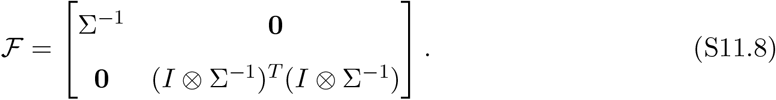

Let *R^T^ R* = *𝓕* be the Cholesky decomposition of *𝓕*. Then, if *y_i_* = *R ·*(*μ_i_*, vec(∑_*i*_))^*T*^, *i* = 1, 2, the above inner product equals 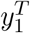· *y*_2_. Thus, we let *WDV^T^* be the classical SVD of *RM* where *D* is a diagonal matrix with entries σ_1_ ≥… ≥σ_*s*_. Then the columns *V_i_* of *V* and the vectors *U_i_* = *R^−^*^1^*W_i_*(*W_i_* the *i*th column of *W*) satisfy (S11.4). This also gives a practical algorithm to calculate the *V_i_, U_i_* and σ_*i*_.

### 12 Multiple crossings in each rotation

The exact stochastic trajectories considered in this article are generated by the SSA that simulates from the Markov jump process with rates dependent on the state of the system as described in **I**. In this Markov jump process, at almost every state of the system, *X*(*t*), with phase *G*(*X*(*t*)) = *g*(*s*), *s* > 0, there is a possibility for the next point, *X*(*t* +*δ*), where *δ* the time of the next jump, to have phase *G*(*X*(*t* + *δ*)) = *g*(*s′*) with *s′* < *s*. That is, the probability of backward (in terms of phase) moves is positive in almost every state, with only few exceptions (e.g. absorbing states).

If we fix a phase and consider the crossings of the exact stochastic trajectory to the corresponding transversal section, the probability of more than one crossings in each rotation is again almost always positive. There are a few related measures that are worth considering here. First, the number of crossings in each rotation that we know it is finite as the Markov jump process can only perform a finite number of jumps in a finite time interval. An interesting question is how fast the probability of the number of crossings declines with increasing number of crossings. Secondly, how much time does the stochastic trajectory spend in this forward and backward moves and how does this compare with the period of the limit cycle. Thirdly, whether the distribution of the first crossings differ from the distribution of the second crossings and henceforth.

To address these questions we performed a simulation study using the *Drosophila* circadian clock system for system size Ω = 300. We computed 1000 simulated trajectories by letting the SSA run for a time interval of 4.5 periods of the limit cycle solution. Note that this simulation study requires all jumps of the SSA to be recorded and therefore it requires a substantially greater computer memory space than the other simulation studies where each trajectory is thinned.

As we can see in Figure A, the probability of the number of crossings declines exponentially fast, with only ≈ 20% of the trajectories giving more than 5 crossings. Furthermore, as we can see in Figure B, the distribution of the time between the first and the last crossing is also declining exponentially fast with only ≈ 5% time intervals being larger than 0.014 which is *O*(10^−4^) compared to the period of the limit cycle. Finally, as we can see in Figure C, the distribution of the points at the different crossings of the same transversal section does not differ substantially in any rotation or transversal coordinate. The deviations are expected to decline with smaller variability levels (larger Ω) as well as the number of crossings and the time spend at each phase.

**Figure A:**
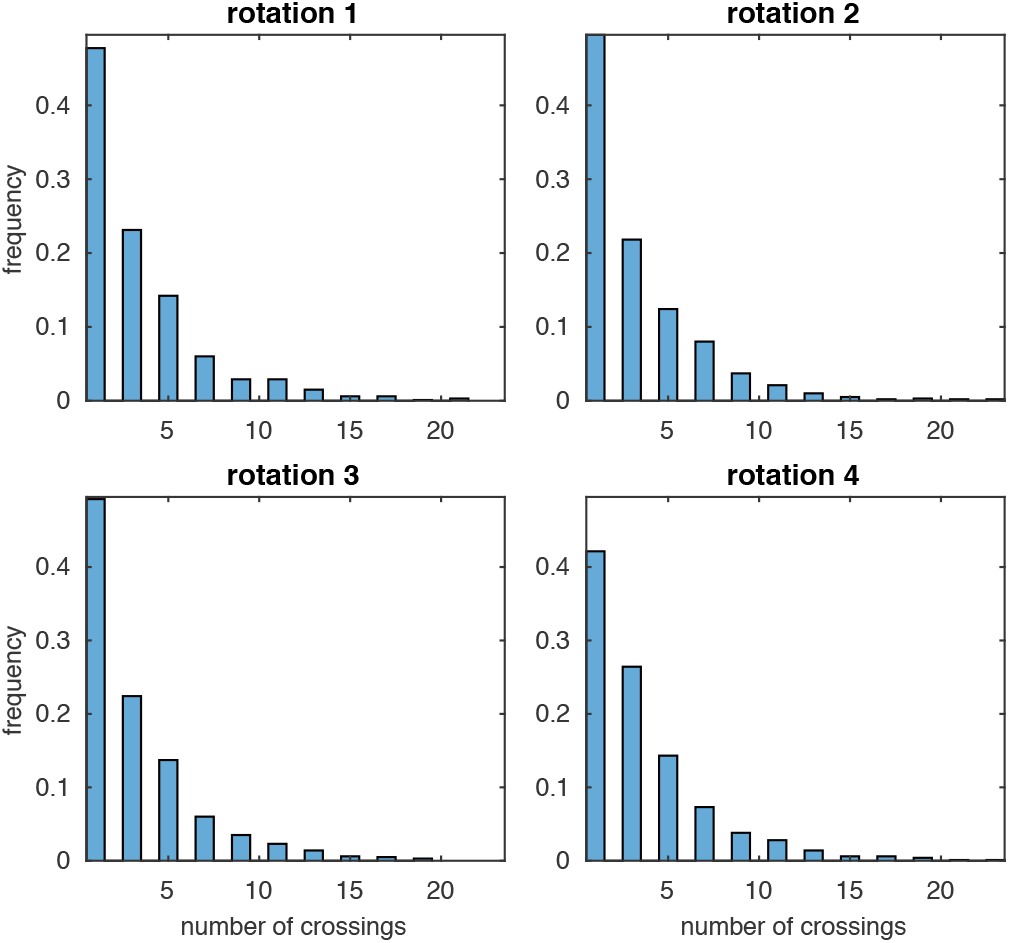
Histograms of the empirical distribution of the number of crossings from a transversal section in each of the four rotations of the stochastic trajectories of the *Drosophila* circadian clock simulated using the SSA at system size Ω = 300.

**Figure B:**
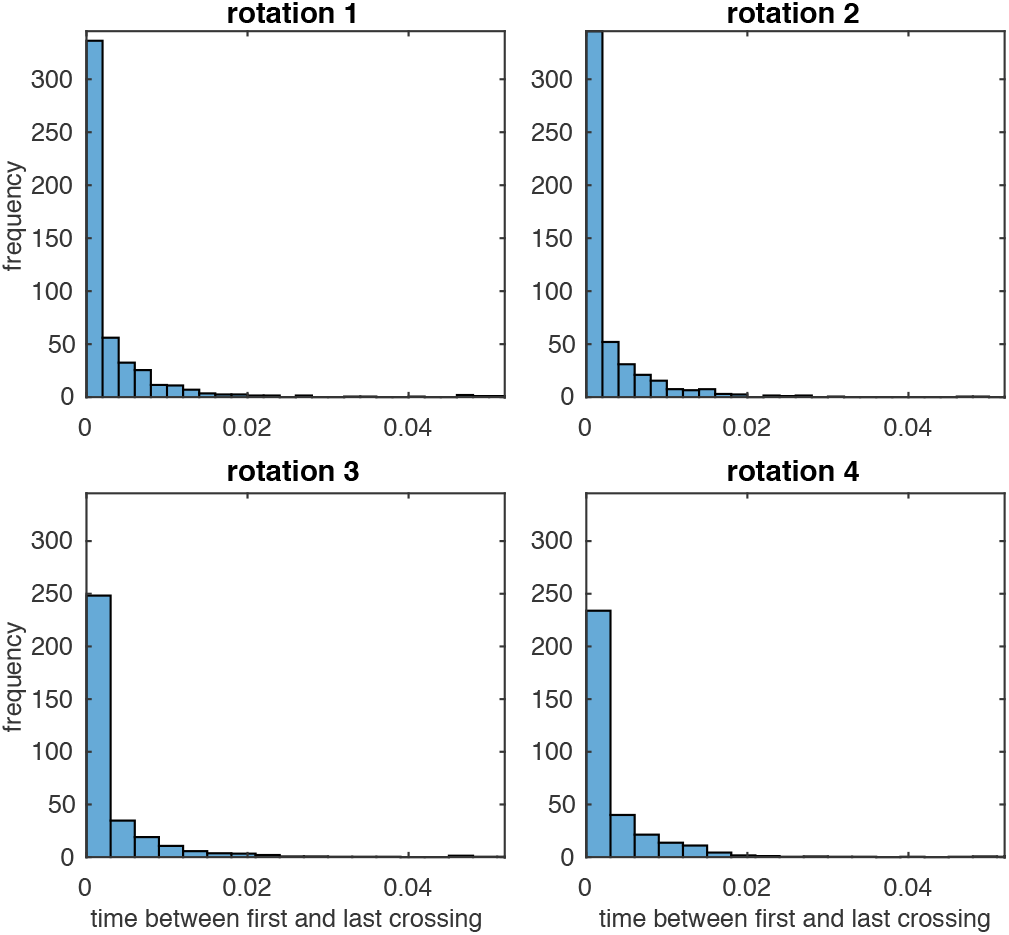
Histograms of the empirical distribution of the time between the first and the last crossing from a transversal section in each of the four rotations of the stochastic trajectories of the *Drosophila* circadian clock simulated using the SSA at system size Ω = 300.

**Figure C:**
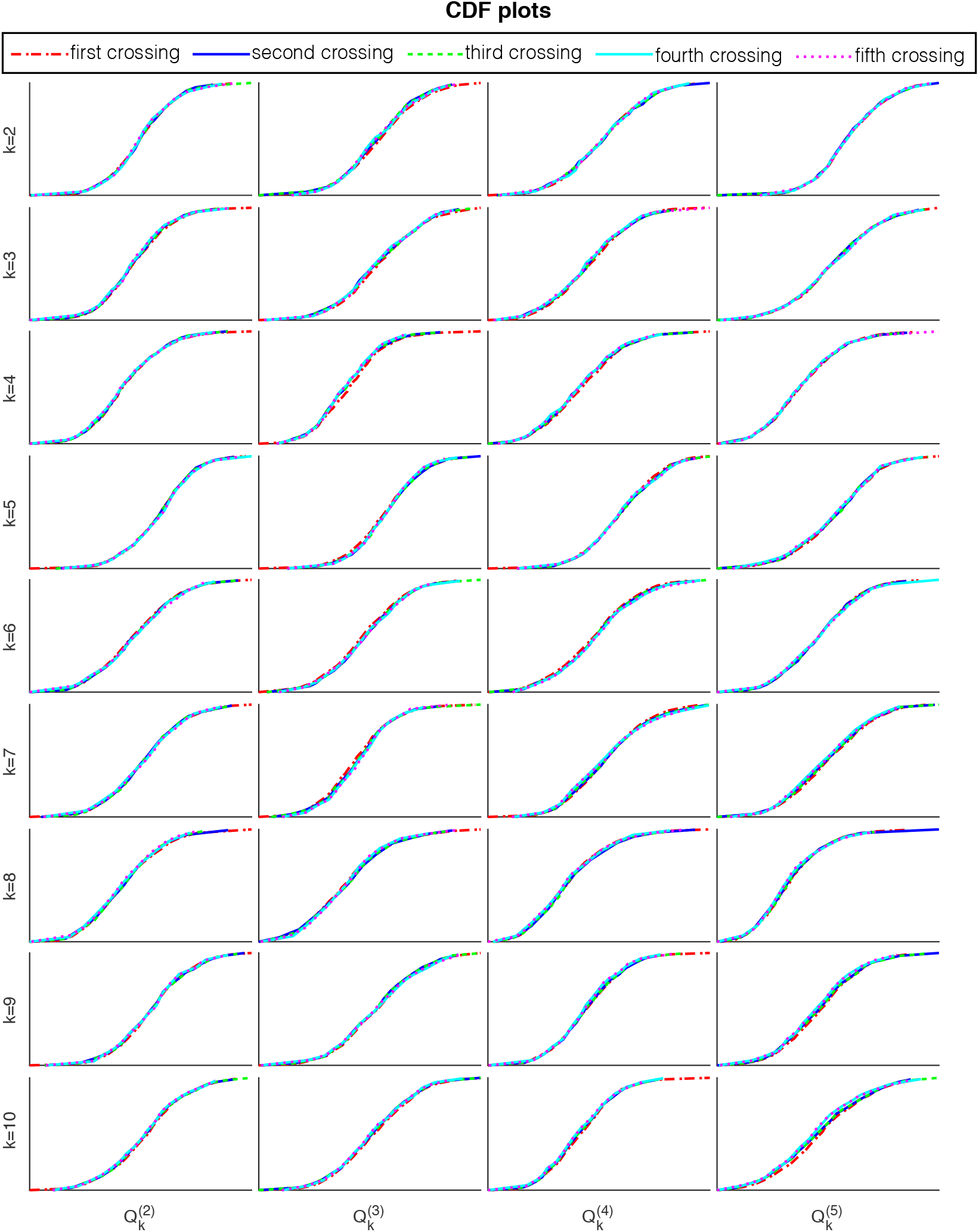
Empirical CDF plots of the distribution of the points, 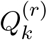, on the transversal section at the first, second, …, fifth crossings (see legend) in each of the four rotations, *r* = 1, 2, 3, 4 and for transversal coordinates *k* = 2, 3, …, 10 of the stochastic trajectories of the *Drosophila* circadian clock simulated using the SSA at system size Ω = 300.

This suggests that the transversal points derived using the thinned stochastic trajectories of the SSA are good representation of all crossings, at least for the variability levels considered in this article.

It is also worth clarifying here that for the pcLNA transversal distributions, as we discuss in Section S6 the distributions of the points in all crossings is considered and then approximated by the distribution of the crossing point at a specific time.

### 13 Negative populations

For the tau-leap, integration of CLE and pcLNA approximation algorithms considered here, the probability of generating negative populations is non-zero. To make the comparison fair to the SSA exact algorithm, we slightly modify these approximation algorithms to ensure that no negative species populations are derived. We use the same simple approach for all approximation algorithms to ensure the comparison between the approximations is fair. This is simply to replace any steps that result in negative populations by an SSA simulation run for the same time interval and then continue with the approximation algorithm. This increases CPU times as SSA is slower than the simulation algorithms, but also improves precision by introducing some bias towards SSA results. However, from our experience, the effect of this change in our comparisons is minimal in terms of precision and substantially affects CPU times only in rare cases. In particular, for large Ω ( = 1000, 3000) the SSA simulation is very rarely employed. For pcLNA, we also take advantage of the property, which also applies to standard LNA, of being able to analytically compute the distribution at the next time-point which is MVN and, before using the SSA step to replace negative populations, we sample from a truncated MVN distribution. For the latter we use a Matlab program^1^ that is much faster than SSA simulation but in the challenging circumstances of the illustrations used in this article, it often fails to generate samples. In that case, an SSA simulation is used as in the other algorithms.

### 14 PeTTSy

PeTTSy is freely available at http://www2.warwick.ac.uk/fac/sci/systemsbiology/research/software/.

In PeTTSy the drift and diffusion matrices, *C*(*t*) and *V*(*t*), respectively, are computed for a large set of time-points. We next provide the description of the computation method for the drift matrix *C*(*s, t*).

The deterministic solution is divided into a number of blocks, *tb_i_, tb_i_*_+1_ *…, tb_N_*. By default, *N* = 70 blocks. The model Jacobian *J*(*t*) is integrated over each of these time blocks in both directions to give forward and reverse solutions for each time block. Forward solution, *C*(*s, t*) where *s* is *tb_i_*, *t* runs from *tb_i_* to *tb_i_*_+1_ is a solution of

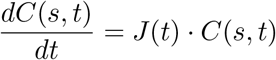

with initial condition *C*(*s, s*) = *I* and *I* the identity matrix. Reverse solution, *C*(*s, t*), where *t* is *tb_i_*, *s* runs from *tb_i_* to *tb_i−_*_1_ is a solution of the adjoint equation

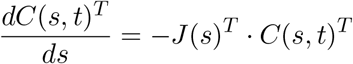

with initial condition *C*(*t, t*) = *I*. This is equivalent to solving,

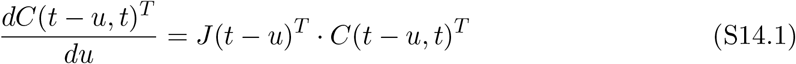

for *t* = *tb_i_*, *i* = 1, 2,…, *N*, and where *u* runs from 0 to *tb_i_− tb_i−_*_1_. The results of this is that the forward and backward solutions allow us to construct any solution, *C*(*a, b*), where if *a< tb_i_* and *b> tbi*, we can write the following

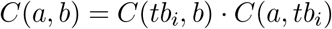

where *C*(*a, tbi*) comes from the reverse solutions and *C*(*tb_i_, b*) comes from the forward solution. However, when *a* and *b* are inside the same interval, [*tb_i_, tb*_*i*+1_] then

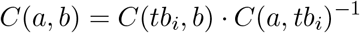

or,

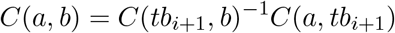

namely we have to invert either the reverse solution or the forward solution in order to calculate *C*(*a, b*). PeTTSy provides a condition number that is equal to the minimum of one over the eigenvalue of the forward and backward solutions and if it is too large the matrix may be close to singular and then the inverse inaccurate. This means that forward and reverse solutions were not calculated finely enough, i.e., there were too few time blocks, and this will affect the accuracy of the results. One can check the accuracy of the *C*(*s, t*) matrices, after forward and reverse solutions have been calculated, by a plot of the condition numbers. As a guide, if condition numbers are above 1*/eps*, where eps is the floating accuracy of Matlab, the inverse is not accurate. The user is then advised to increase the value of *N*.

In a similar fashion, forward and reverse solutions are computed for the diffusion matrix *V*, using the equation (S4.3). Alternatively, one may solve the ODE

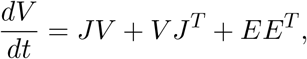

where *E* = *E*(*s*) the matrix described in equation (S4.3), with initial condition *V*(*t*_0_, *t*_0_) = 0. The reverse solutions are derived in a similar fashion to above by using the ODE

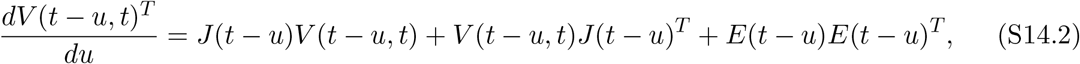

with initial condition *V*(*t*_0_, *t*_0_) = 0, for *t* = *tb_i_*, *i* = 1, 2,…, *N*, and where *u* runs from 0 to *tb_i_* − *tb*_*i*−1_.

### 15 The pcLNA Kalman Filter

The pcLNA Kalman Filter algorithm uses the following recursive algorithm for computing the terms in *L (θ*;*<u>X̂</u>*). Note that (*μ^*^*, ∑^*\**^) denote posterior estimates of (*μ*, ∑) conditional on the observed measurement at the current time.

1. Input (*X̂*(*t_0_*), *X̂*(*t*_1_), …, *X̂*(*t_N_*)), *μ*(*t*_0_), ∑(*t*_0_), *B* and ∑_∈_.
2. Compute *P*(*X̂*(*t*_0_); *θ*) from *MVN*(*μ̂*(*t*_0_), ∑̂ (*t*_0_)).
3. **For iteration** *i* = 1, 2,…

a. Derive (*X*(*t*_*i*-1_) | *X̂*(*t*_*i*-1_)) ~ *MVN*(*μ\**(*t_i-1_*), ∑*(*t*_*i*-1_));
b. Compute *g(s*_*i*-1_) = *G(μ*^*^(*t*_*i*-1_)) and set (*κ(s*_*i*-1_)|*X̂(t*_*i*-1_)) ~ *MVN*(*m*^*^(*s*_*i*-1_), *S^*^(s*_*i*-1_));
c. Derive (*X*(*t_i_*)|*X̂*(*t*_*i*-1_)) ~ *MV N*(*μ*(*t_i_*), ∑(*t_i_*));
d. Compute *P*(*X̂*(*t_i_*)|*X̂*(*t*_*i*-1_); *θ)* from *MV N*(μ ̂ (*t_i_*), ∑̂(*t_i_*)).

In the for loop *μ^*^(t*_i−1_) = *μ(t*_*i*−1_) + ∑(*t*_*i*−1_)*B^T^* ∑̂(*t*_i−1_)^−1^∈ ̂(*t*_i−1_), ∑^*^(*t*_*i*−1_) = ∑(*t*_*i*−1_) − ∑(*t*_*i*−1_)*B^T^* ∑̂(*t*_*i*−1_)^−1^*B*∑(*t*_*i*−1_), *m^*^(s*_*i*−1_) = Ω^1/2^(*μ^*^(t*_i−1_) − *g(s*_i−1_)), with *∈ ̂(t_i_)* = *X̂(t_i_)* − *μ̂(t_i_)* and

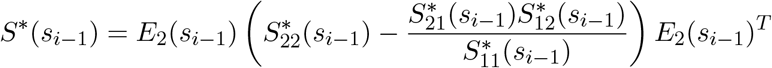

where the matrix *E*_2_(*t*) = [*e*_2_(*t*)…*e_n_*(*t*)] has columns orthonormal vectors which are also orthogonal to the unit tangent vector *e*_1_(*t*) = *F*(*x*(*t*))/‖*F*(*x*(*t*))‖ and

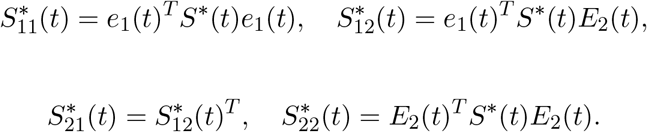

The measurement equation (Eq. 12 in **I**) is used to derive the predictive probabilities *P*(*X̂*(*t*_0_); *θ*) and *P*(*X̂*(*t_i_*)*|X̂*(*t_i−_*_1_); *θ*) in steps 2 and 3(d), respectively. The posterior distribution (*X*(*t*)|*X̂*(*t*)) in step 3(a) is derived by Bayes rule, while the posterior distribution of the corrected noise process (*κ*(*s*)|*X̂*(*t*)) in step 3(b) is derived by restricting the posterior distribution of the noise process on the transversal section *𝒮_g_*_(*s*)_ normal to *e*_1_(*s*) using the Schur complement as described earlier. The cartesian coordinate system is used here. If this correction is omitted, step 3(b) of the above algorithm is replaced by the standard LNA step, (ξ(*t*_*i*−1_)|*X̂*(*t_i−_*_1_)) ~ *MV N*(*m̂*(*t_i−_*_1_), *Ŝ*(*t_i−_*_1_)) where *m̂*(*t_i−_*_1_) = Ω^1/2^*μ̂*(*t_i−_*_1_) − *g*(*t_i−_*_1_), *Ŝ*(*t_i−_*_1_) = Ω∑^ˆ^(*t_i−_*_1_). The LNA Ansantz (Eq. S4.1) and transition (Eq. S4.2) are used to derive the prior distribution (*X*(*t_i_*)*|X̂*(*t_i−_*_1_)) in step 3(c).

### 16 Details related to various computations

All computations for the illustrations in **I** and SI including:

1. SSA, tau-leap, integration of diffusion and pcLNA simulation algorithms described in Sections “pcLNA Simulation algorithm” and “Comparisons to other simulation algorithms” (see also the note in S1 Sect. 13),
2. the computation of Fisher Information and principal control coefficients for the sensitivity discussed in the Section “Sensitivity analysis for stochastic systems” and S1 Sect. 11

are done using the PeTTSy software which is discussed in S1 Sect. 14 and it is freely available at http://www2.warwick.ac.uk/fac/sci/systemsbiology/research/software/.

The phase correction that involves identifying the normal mapping *G*(*X*) = *x* ∈ *γ* of a given state *X* = *X*(*t*) of the trajectory is performed by Newton’s optimisation minimising the square of the function

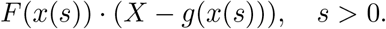

Note that the minimum is achieved when *s >* 0 is such that *X* − *g*(*x*(*s*)) is normal to the tangent vector *F*(*x*(*s*)). Multiple repetitions of the above optimisation are performed, using different initial conditions *s >* 0, to ensure that all local minima, *s′* = *s*_1_, *s*_2_,…, are derived and then, as explained in S1 Sect. 1, *G*(*X*(*t*)) = *x*(*s*) with *s* = min_*i*_ |*s_i_* − *t*| the closest time to *t*. Note that other methods of optimisation could be equally applicable.

**FIM computation** We use the results in [5] to derive the necessary derivatives for the computation of the FIM. For the readers’ convenience some of the derivations in chapter V in [5] are reproduced here. Let

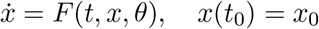

be the initial value problem with parameter vector *θ* which has solution *x* = *g*(*t, t*_0_, *x*_0_, *θ*). The Jacobian matrix *J*(*t*) = *J*(*t, x, θ*) at *x* = *g*(*t, t*_0_, *x*_0_, *θ*) has entries

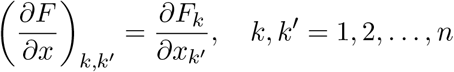

and the Hessian matrix *H*(*t*) = *H*(*t, x, θ*) at *x* = *g*(*t, t*_0_, *x*_0_, *0θ*) has entries

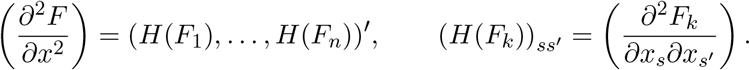

Define

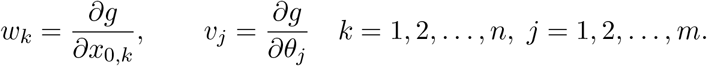

Then, *w_k_* is the solution of

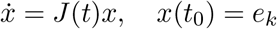

and *e_k_* a vector with all zero entries except the *k*th entry which is equal to 1.

Note that the principal fundamental matrix *C* is the solution of the latter initial value problem. The matrix *C*(*t*_0_, *t*) has columns *w_k_*(*t*), *k* = 1, 2,…, *n*. Furthermore, *v_j_* is the solution of

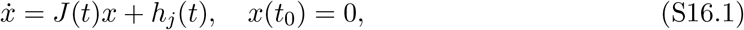

where *h_j_*(*t*) = ∂*F/*∂*θ_j_*.

The *mn* × *n* matrix (*dC/dθ*_1_ *… dC/dθ_m_*)^*T*^ is the solution of

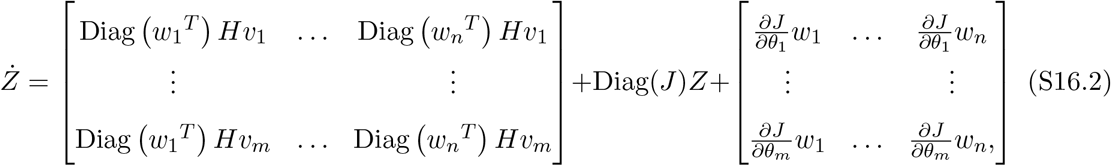

where Diag(*J*) is the *mn* ×; *mn* block matrix with main diagonal block entries the *n* × *n* matrix *J*. The *mn* × *n* matrix (*dV/dθ*_1_,…, *dV/dθ_m_*)^*T*^ has blocks *dV/dθ_j_*, *j* = 1, 2,…, *m* which are the solutions of

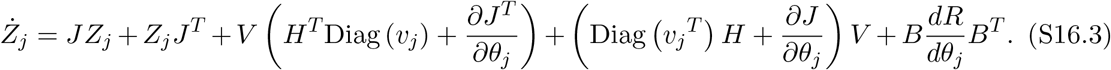

Here *B* is the stoichiometry matrix of the system and *dR/dθ_j_* the diagonal matrix with main diagonal entries the derivatives of the reaction rates

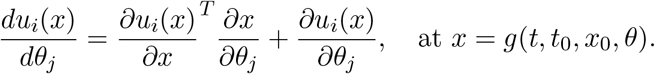

In Section “Fisher Information” of **I** we discuss properties of the FIM of the distribution

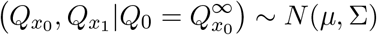

where 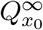 the fixed point pcLNA distribution at *x*_0_ = *g*(*t*_0_). The parameters of this distribution are

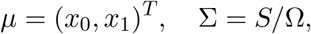

and *S* = *A*^−1^ with

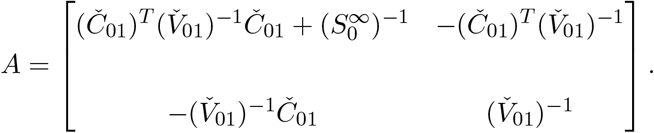

Here

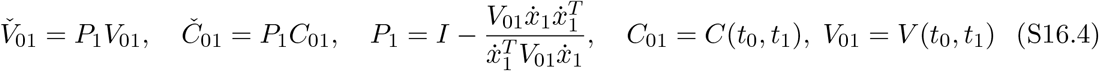

and 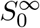 the solution of the contraction equation

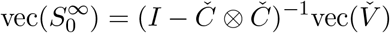

where

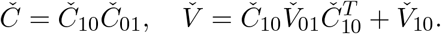

The matrices *V̌*_01_ and *Č*_01_ are derived in (S16.4) and

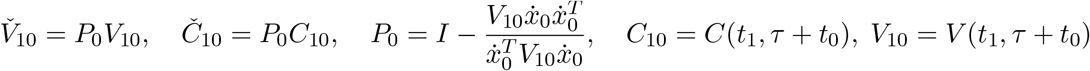

and τ the period of the system. Thus,

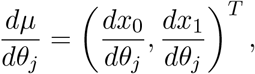

and

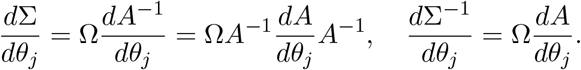

The derivative *dA/dθ_j_* is a block matrix with blocks

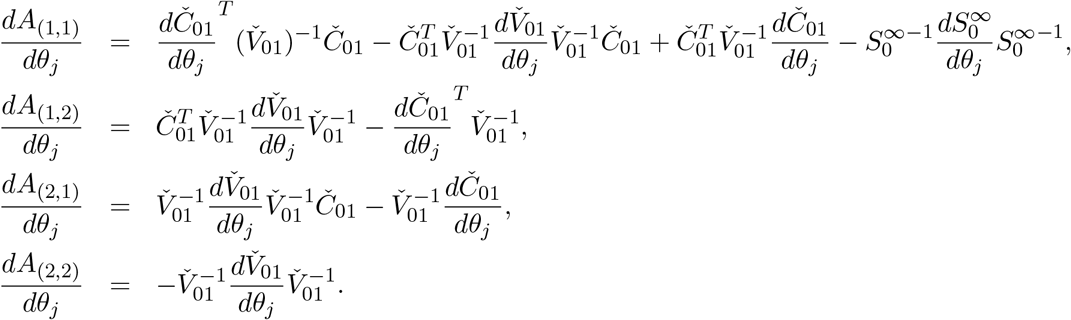

The necessary derivatives for computing the entries

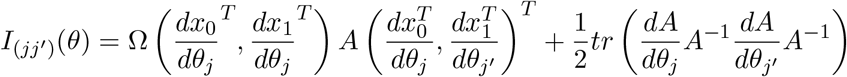

of the FIM are derived as described above. In particular, *v_j_* = *dx_i_/dθ_j_* is the solution of (S16.1) and the derivative *dẋ*_*i*_/*dθ*_*j*_ used to derive *dP_i_/dθ_j_* is equal to *J*(*t_i_*)*v_j_* + *g_j_*(*t_i_*). Furthermore, the derivatives *dC/dθ_j_* and *dV/dθ_j_* are derived as solutions of the initial value problems in (S16.2) and (S16.3), respectively.

1 http://uk.mathworks.com/matlabcentral/fileexchange/34402-truncated-multivariate-normal

## Supplementary information S2 for the paper Long-time analytic approximation of large stochastic oscillators: simulation, analysis and inference

> We refer to the paper “Long-time analytic approximation of large stochastic oscillators: simulation, analysis and inference” by **I.** In this note we give details about the *Drosophila* circadian clock system and use it to illustrate further the accuracy of distributions and simulations discussed in **I**.

### 1 *D*rosophila circadian clock system

A schematic representation of the *Drosophila* circadian clock system, as provided in [1], is displayed in Figure A. The oscillations are driven by the negative feedback exerted on the *per* and *tim* genes by the complex formed from PER and TIM proteins following phosphorylation. The *per* and *tim* mRNA, *M_P_* and *M_T_*, respectively, are transported into the cytosol where they are degraded and translated into protein (*P*_0_ and *T*_0_). These proteins are multiply phosphorylated (PER: *P*_0_ →*P*_1_ → *P*_2_; TIM: *T*_0_ →*T*_1_ →*T*_2_) and these modifications can be reversed by a phosphatase. The fully phosphorylated form of the proteins is targeted for degradation and forms a complex, *C*, which is transported into the nucleus in a reversible manner where the nuclear form of the PER–TIM complex, *C_N_*, represses the transcription of *per* and *tim* genes.

**Figure A:**
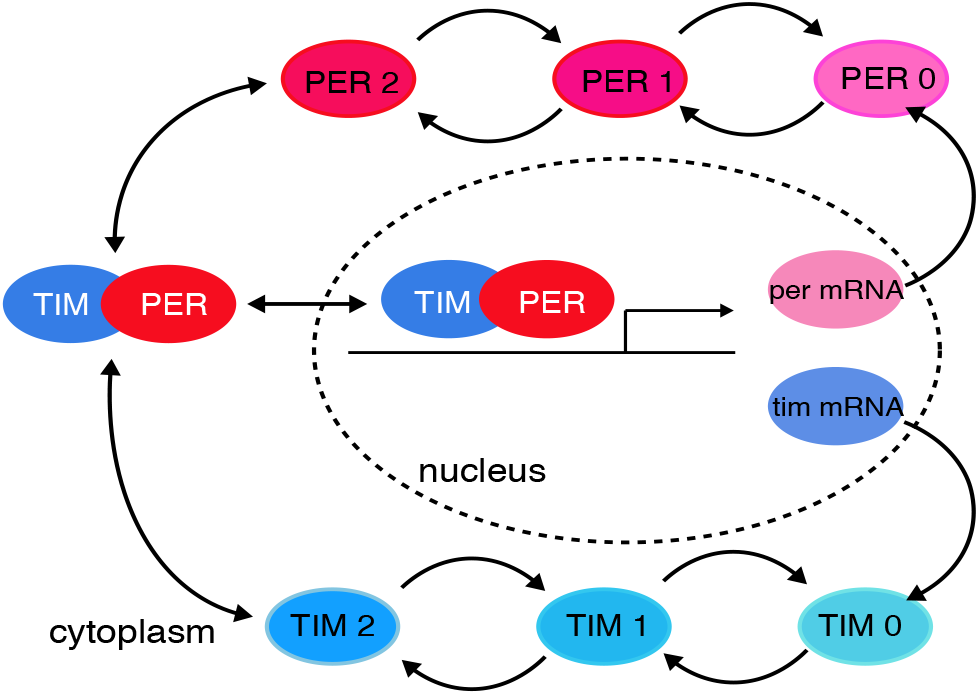
Schematic representation of *Drosophila* circadian clock system [1]

The variables of the model along with the initial conditions (in nanomolar concentrations) used in our implementation are provided in Table A.

**Table A:**
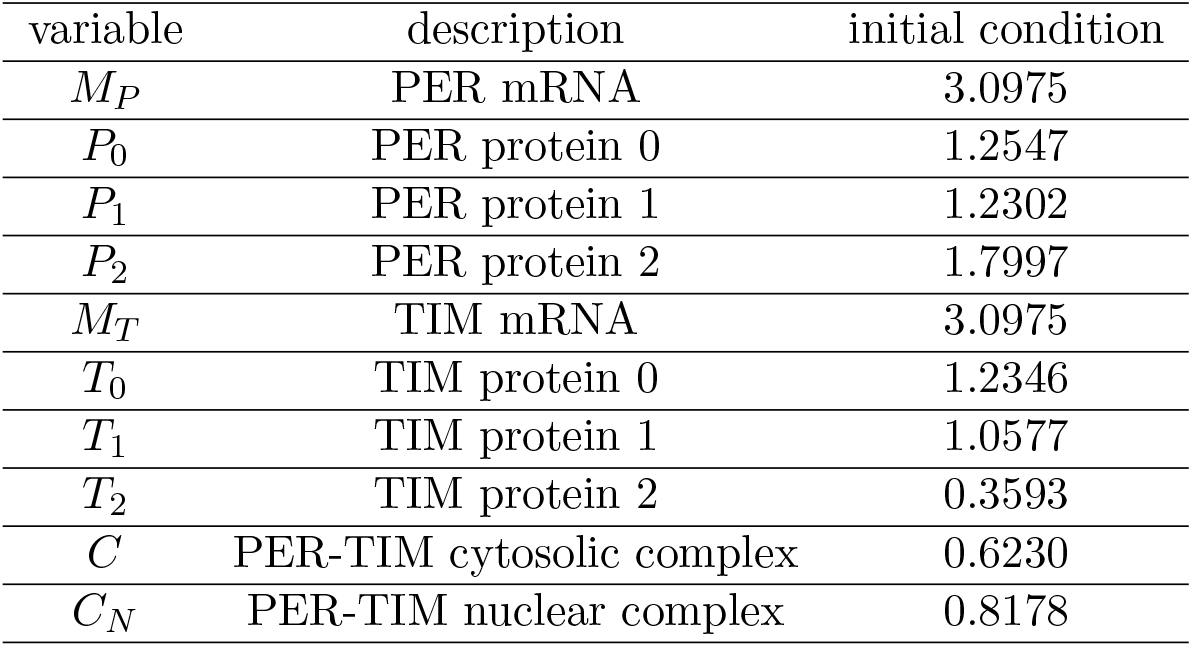
The variables of *Drosophila* circadian clock system and the initial conditions (in nanomolar concentrations) used to derive their ODE solution.

| variable | description | initial condition |
| --- | --- | --- |
| $M_P$ | PER mRNA | 3.0975 |
| $P_0$ | PER protein 0 | 1.2547 |
| $P_1$ | PER protein 1 | 1.2302 |
| $P_2$ | PER protein 2 | 1.7997 |
| $M_T$ | TIM mRNA | 3.0975 |
| $T_0$ | TIM protein 0 | 1.2346 |
| $T_1$ | TIM protein 1 | 1.0577 |
| $T_2$ | TIM protein 2 | 0.3593 |
| $C$ | PER-TIM cytosolic complex | 0.6230 |
| $C_N$ | PER-TIM nuclear complex | 0.8178 |

The parameter values used to derive the ODE solution of the system are provided in Table B.

**Table B:** The parameters of *Drosophila* circadian clock system and the values used to derive their ODE solution.

| parameter | description | value | measurement unit |
| --- | --- | --- | --- |
| $v_{sP}$ | $M_P$ transcription | 1.10 | $nMh^{-1}$ |
| $v_{sT}$ | $M_T$ transcription | 1.00 | $nMh^{-1}$ |
| $v_{mP}$ | $M_P$ degradation | 1.00 | $nMh^{-1}$ |
| $v_{mT}$ | $M_T$ degradation | 0.70 | $nMh^{-1}$ |
| $v_{dP}$ | $P_2$ degradation | 2.20 | $nMh^{-1}$ |
| $k_{sP}$ | $M_P$ translation | 0.90 | $h^{-1}$ |
| $k_{sT}$ | $M_T$ translation | 0.90 | $h^{-1}$ |
| $k_1$ | $C \rightarrow C_N$ | 0.80 | $h^{-1}$ |
| $k_2$ | $C_N \rightarrow C$ | 0.20 | $h^{-1}$ |
| $k_3$ | $P_2 + T_2 \rightarrow C$ | 1.20 | $h^{-1}$ |
| $k_4$ | $C \rightarrow P_2 + T_2$ | 0.60 | $h^{-1}$ |
| $K_{mP}$ | $M_P$ enzymatic degradation | 0.20 | $h^{-1}$ |
| $K_{mT}$ | $M_T$ enzymatic degradation | 0.20 | $h^{-1}$ |
| $K_{IP}$ | $M_P$ Hill coefficient | 1.00 | $h^{-1}$ |
| $K_{IT}$ | $M_T$ Hill coefficient | 1.00 | $h^{-1}$ |
| $K_{dP}$ | $P_2$ enzymatic degradation | 0.20 | $h^{-1}$ |
| $K_{dT}$ | $T_2$ enzymatic degradation | 0.20 | $h^{-1}$ |
| $k_d$ | linear degradation | 0.01 | $h^{-1}$ |
| $k_{dC}$ | $C$ degradation | 0.01 | $h^{-1}$ |
| $k_{dN}$ | $C_N$ degradation | 0.01 | $h^{-1}$ |
| $v_{dT}$ | $T_2$ degradation | 3.00 | $nMh^{-1}$ |
| $K_{1P}$ | $P_0 \rightarrow P_1$ enzymatic | 2.00 | $h^{-1}$ |
| $K_{1T}$ | $T_0 \rightarrow T_1$ enzymatic | 2.00 | $h^{-1}$ |
| $K_{2P}$ | $P_1 \rightarrow P_0$ enzymatic | 2.00 | $h^{-1}$ |
| $K_{2T}$ | $T_1 \rightarrow T_0$ enzymatic | 2.00 | $h^{-1}$ |
| $K_{3P}$ | $P_1 \rightarrow P_2$ enzymatic | 2.00 | $h^{-1}$ |
| $K_{3T}$ | $T_1 \rightarrow T_2$ enzymatic | 2.00 | $h^{-1}$ |
| $K_{4P}$ | $P_2 \rightarrow P_1$ enzymatic | 2.00 | $h^{-1}$ |
| $K_{4T}$ | $T_2 \rightarrow T_1$ enzymatic | 2.00 | $h^{-1}$ |
| $V_{1P}$ | $P_0 \rightarrow P_1$ | 8.00 | $nMh^{-1}$ |
| $V_{1T}$ | $T_0 \rightarrow T_1$ | 8.00 | $nMh^{-1}$ |
| $V_{2P}$ | $P_1 \rightarrow P_0$ | 1.00 | $nMh^{-1}$ |
| $V_{2T}$ | $T_1 \rightarrow T_0$ | 1.00 | $nMh^{-1}$ |
| $V_{3P}$ | $P_1 \rightarrow P_2$ | 8.00 | $nMh^{-1}$ |
| $V_{3T}$ | $T_1 \rightarrow T_2$ | 8.00 | $nMh^{-1}$ |
| $V_{4P}$ | $P_2 \rightarrow P_1$ | 1.00 | $nMh^{-1}$ |
| $V_{4T}$ | $T_2 \rightarrow T_1$ | 1.00 | $nMh^{-1}$ |
| $n$ | Hill power | 4.00 | NA |

The ODE system for the *Drosophila* circadian clock is

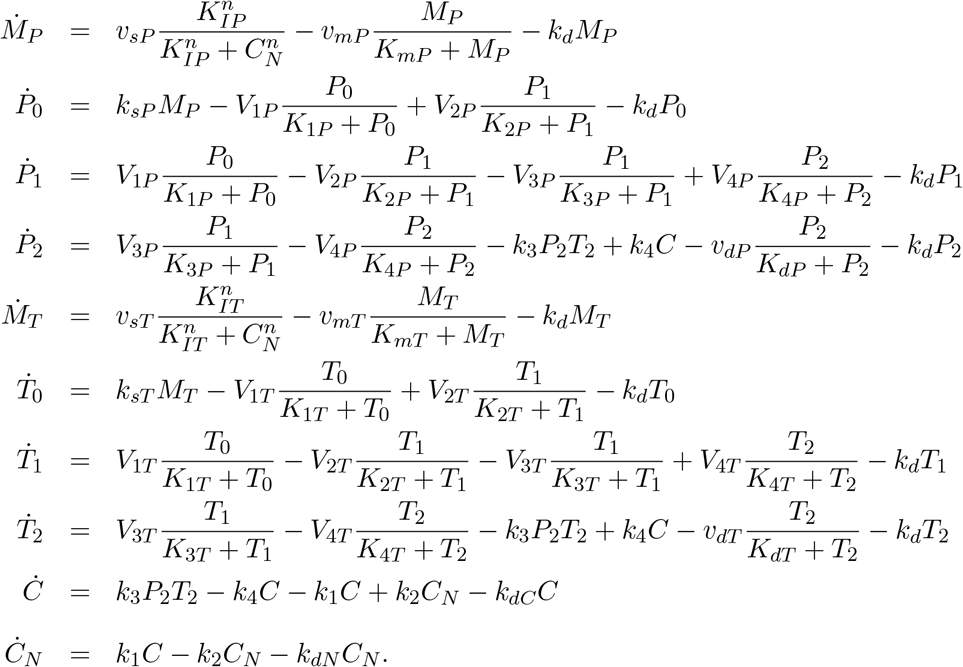

The SSA trajectories are derived using the rates provided in Table 1 of [1].

In the simulations of the *Drosophila* circadian clock provided in the next section and in **I**, we consider five values of the system size Ω = 200, 300, 500, 1000 and 3000. In Table C, we provide the approximate ranges of the molecular numbers for each species observed in our simulations. Notice the increase in the maximum number of molecules with increasing Ω, but also the decrease in the length of the range in terms of concentrations, *X*(*t*) = *Y*(*t*)/Ω, that reflects the lower levels of stochasticity for increasing Ω.

**Table C:** The approximate ranges of the molecular numbers, *Y*(*t*), of each of the species of *Drosophila* circadian clock system in simulations derived using SSA for various system sizes Ω

| variable | $\Omega = 200$ | $\Omega = 300$ | $\Omega = 500$ | $\Omega = 1000$ | $\Omega = 3000$ |
| --- | --- | --- | --- | --- | --- |
| $M_P$ | (0, 800) | (0, 1200) | (0, 1900) | (0, 3500) | (40, 10000) |
| $P_0$ | (0, 500) | (0, 600) | (0, 900) | (0, 1600) | (10, 4500) |
| $P_1$ | (0, 500) | (0, 700) | (0, 1000) | (0, 1700) | (10, 4700) |
| $P_2$ | (0, 1400) | (0, 1700) | (0, 2500) | (0, 4300) | (30, 10600) |
| $M_T$ | (0, 800) | (0, 1200) | (0, 1900) | (0, 3500) | (40, 10000) |
| $T_0$ | (0, 400) | (0, 600) | (0, 900) | (0, 1600) | (10, 4000) |
| $T_1$ | (0, 400) | (0, 500) | (0, 800) | (0, 1400) | (10, 3900) |
| $T_2$ | (0, 200) | (0, 300) | (0, 400) | (0, 600) | (10, 1400) |
| $C$ | (6, 400) | (10, 500) | (30, 800) | (75, 1400) | (280, 3700) |
| $C_N$ | (60, 700) | (100, 900) | (200, 1400) | (400, 2600) | (1300, 7000) |

For more details considering the *Drosophila* circadian clock system see [1].

### 2 Exact simulations

**Figure B:**
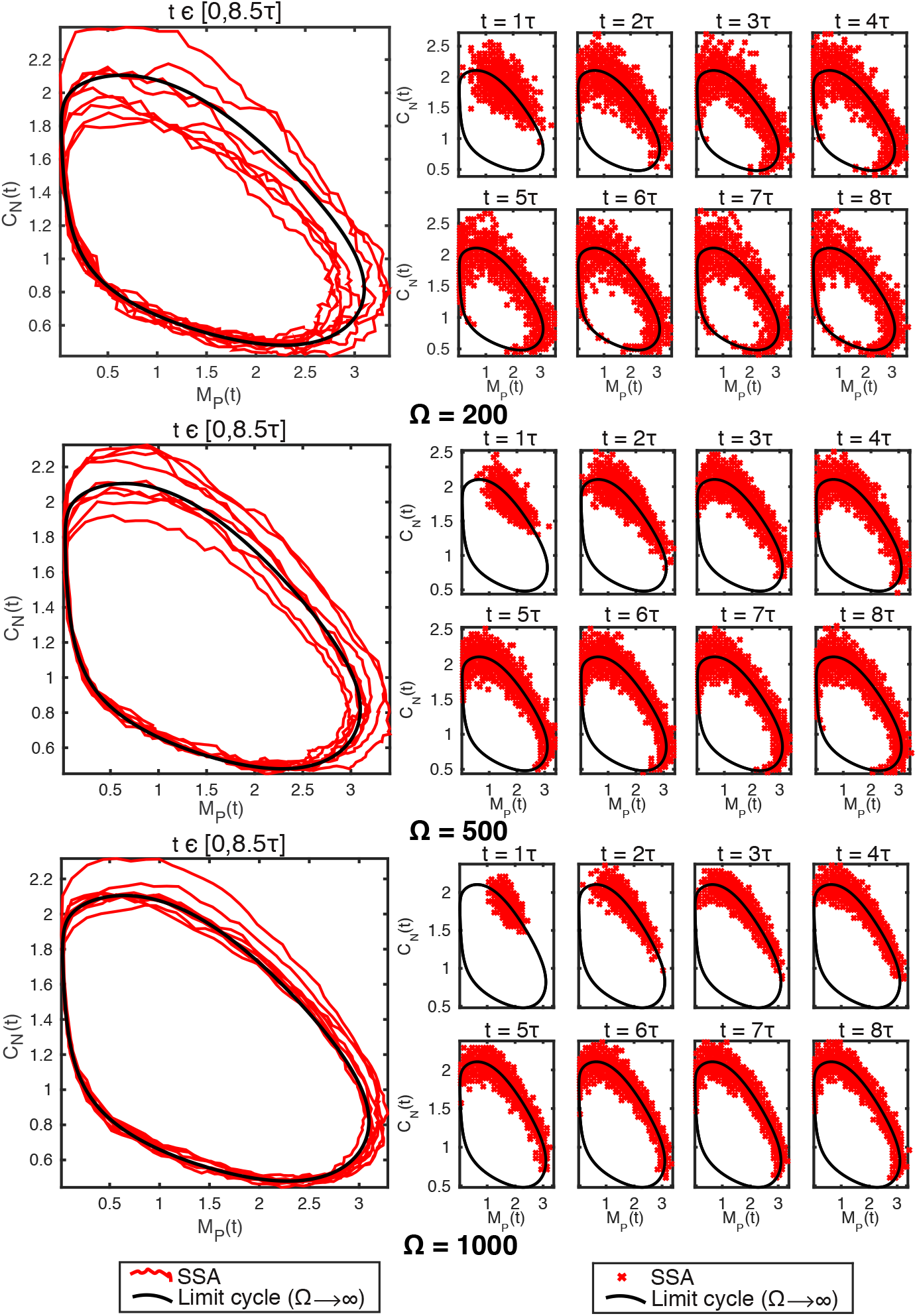
Exact stochastic simulation of the *Drosophila* circadian clock system in three different system sizes, Ω = 200 (top panel), Ω = 500 (middle panel), Ω = 1000 (bottom panel). A stochastic trajectory obtained by running the SSA over the time-interval *t* ∈[0, 8.5_*τ*_] is displayed on the left panels and SSA samples (*R* = 3000) at times *t* = *τ*, 2*τ, …*, 8*τ* are displayed on the right panels. Two (out of 10) of the species are displayed (*per* mRNA *M_P_* (x-axis) and nuclear PER-TIM complex *C_N_* (y-axis)). The black solid curve is the large volume, Ω → ∞, limit cycle solution.

### 3 Exact transversal distributions: normality

**Figure C:**
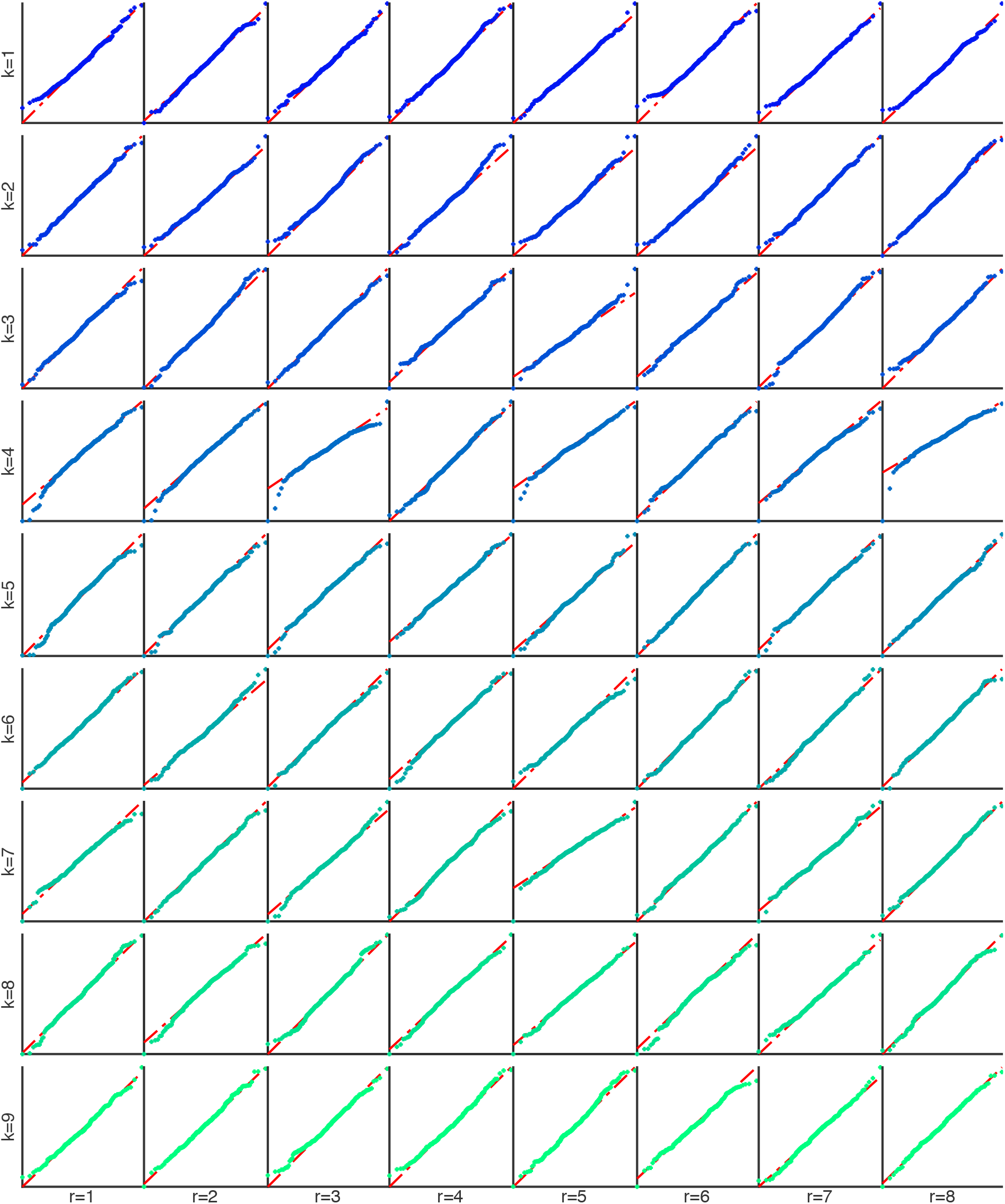
Quantile-Quantile (Q-Q) plots of the distribution of the first intersection in the *r*-th pass, 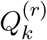, in transversal coordinates *k* = 2, 3,…,10 for *r* = 1, 2,…,8.

### 4 pcLNA and exact transversal distributions

**Figure D:**
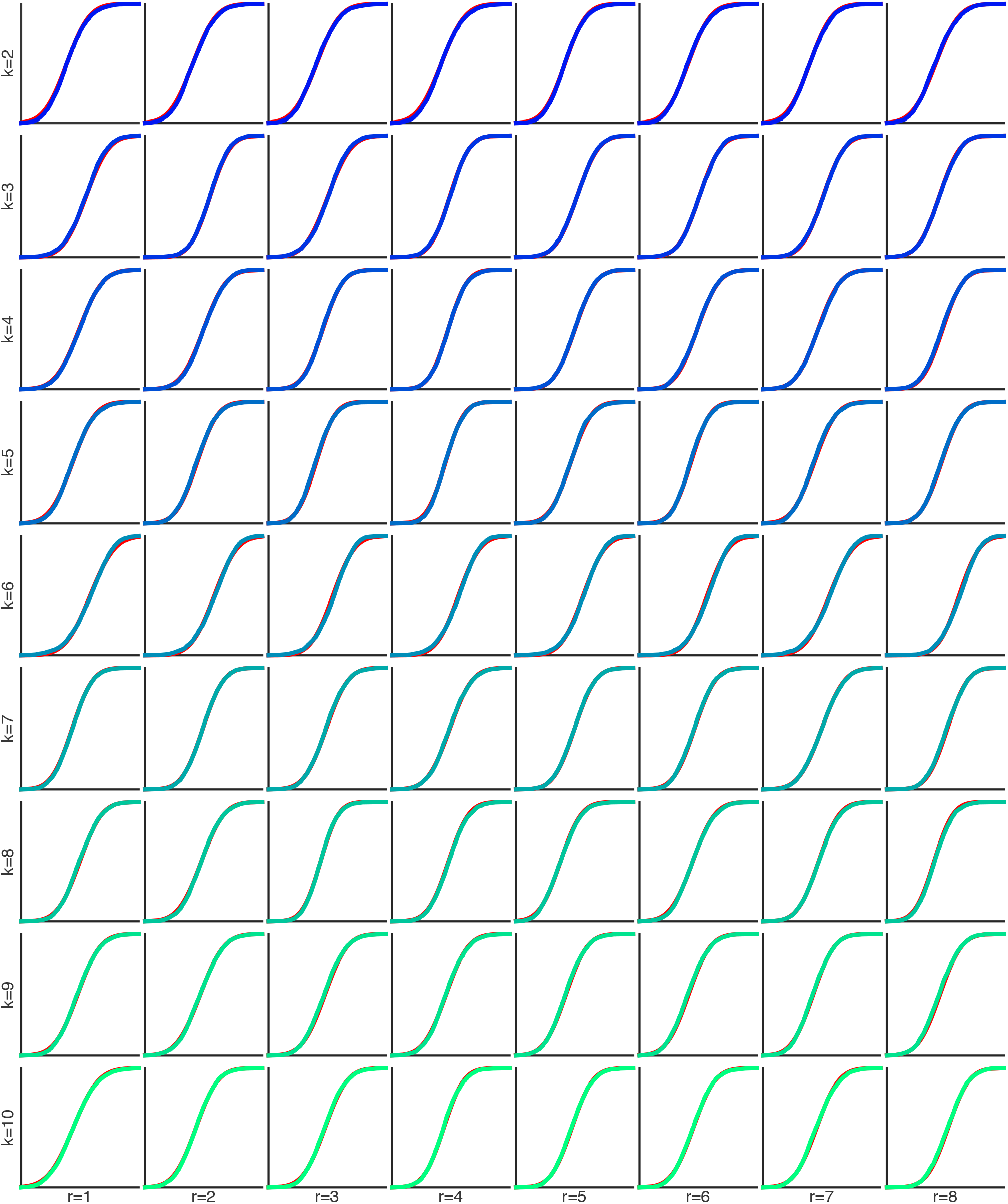
CDF plots of the transversal distributions 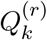under the pcLNA (red line) and the SSA (empirical CDF, crosses) for transversal coordinates *k* = 2, 3,…, 10 and round *r* = 1, 2,…, 8.

**Figure E:**
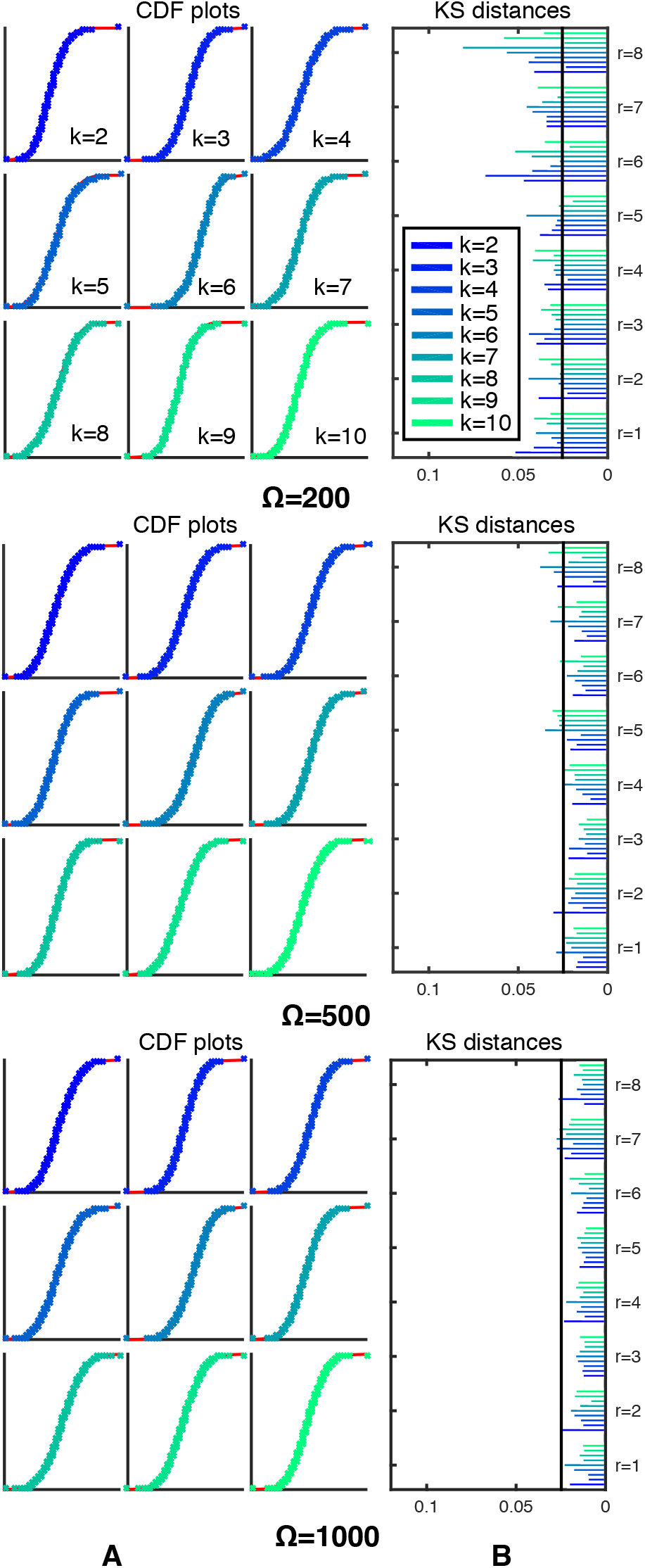
Comparison of pcLNA and exact transversal distributions in systems sizes Ω = 200 (top panel), Ω = 500 (middle panel), Ω = 1000 (bottom panel).(A) CDF plots under the pcLNA (red line) and the SSA (empirical CDF, crosses) of the first intersection in the *r*-th pass, 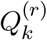, in transversal coordinates *k* = 2, 3,…, 10 for *r* = 4. (B) KS distances between the 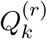distributions under pcLNA and SSA, *r* = 1, 2,…, 8, *k* = 2, 3,…, 10.

### 5 Less frequent correction

**Figure F:**
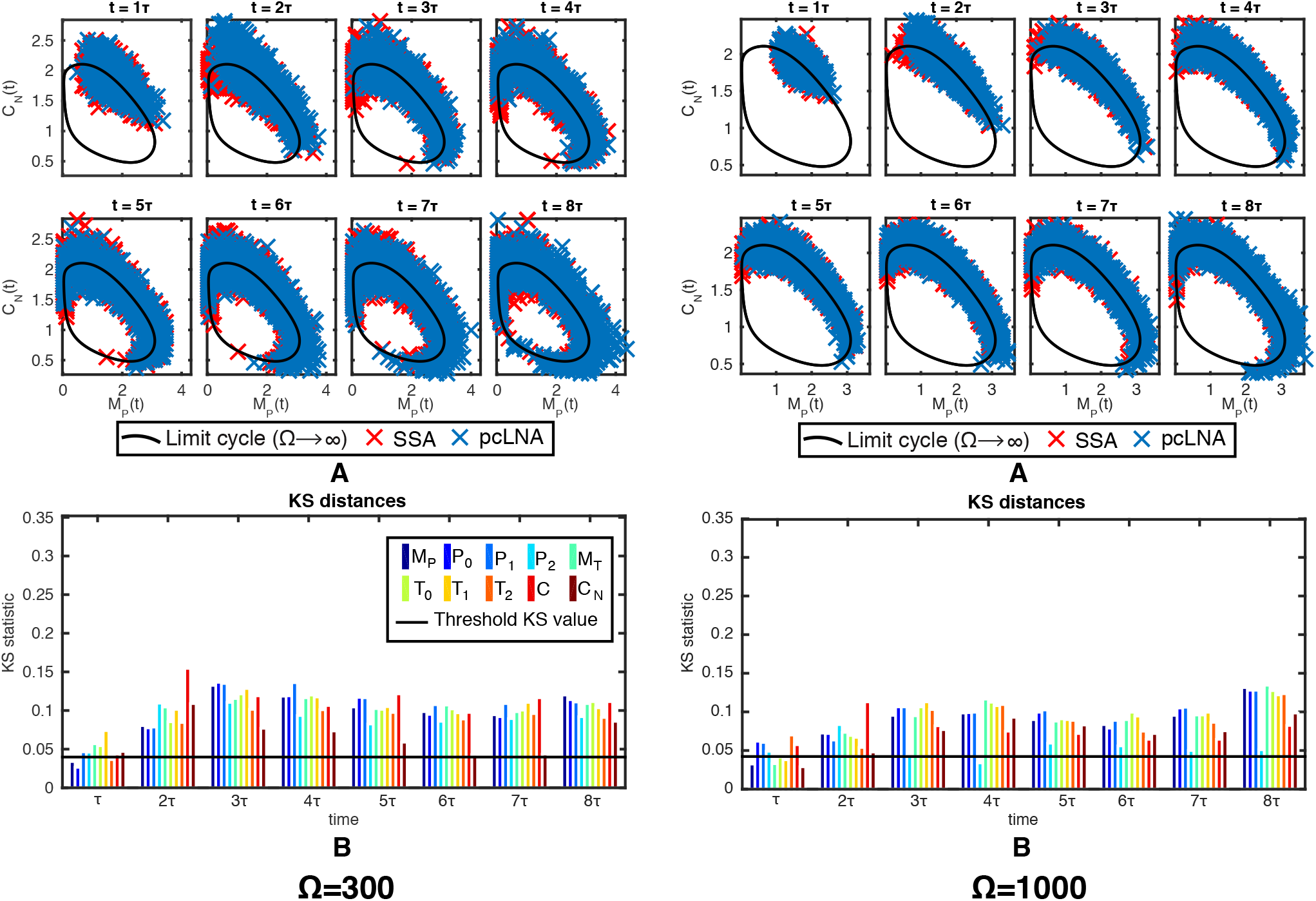
Comparison between pcLNA and SSA simulations for correction frequency 24*h*. This is 3 times less frequent than the correction used in the simulations presented in Fig. 9 of **I** and the figures in the next section. Panels (A) contain the samples produced by the pcLNA and the exact simulation (SSA) at time-points, *t* = 1*τ*, 2*τ, …*, 8*τ*, Ω for = 300 (left panel) and Ω = 1000 (right panel), and panels (B) the KS distances between the empirical distributions of pcLNA and the SSA for each system variable (colored bars) for two system sizes Ω = 300 (left panel) and Ω = 1000 (right panel).

Less frequent phase correction in pcLNA algorithm has substantial impact on pcLNA simulations. However, the precision of the approximation seems to stabilise after the third round with KS distances much smaller compared to standard LNA. The median CPU times under this correction frequency are 0.45*secs* for Ω = 300 and 0.22*secs* for Ω = 1000, compared to 0.45*secs* and 0.42*secs*, respectively, for correction frequency 6*h*. The reason for the absence of speed improvement for Ω = 300 is because at this system size and with less corrections negative populations appear more frequently and therefore SSA simulation are used more frequently, as explained in Sec. 13 in **S**1, and therefore any improvements due to applying correction less frequently are counterbalanced. For Ω = 1000, where negative populations are much less likely the speed improvement, is almost twofold.

### 6 Comparison between different simulation algorithms

**Figure G:**
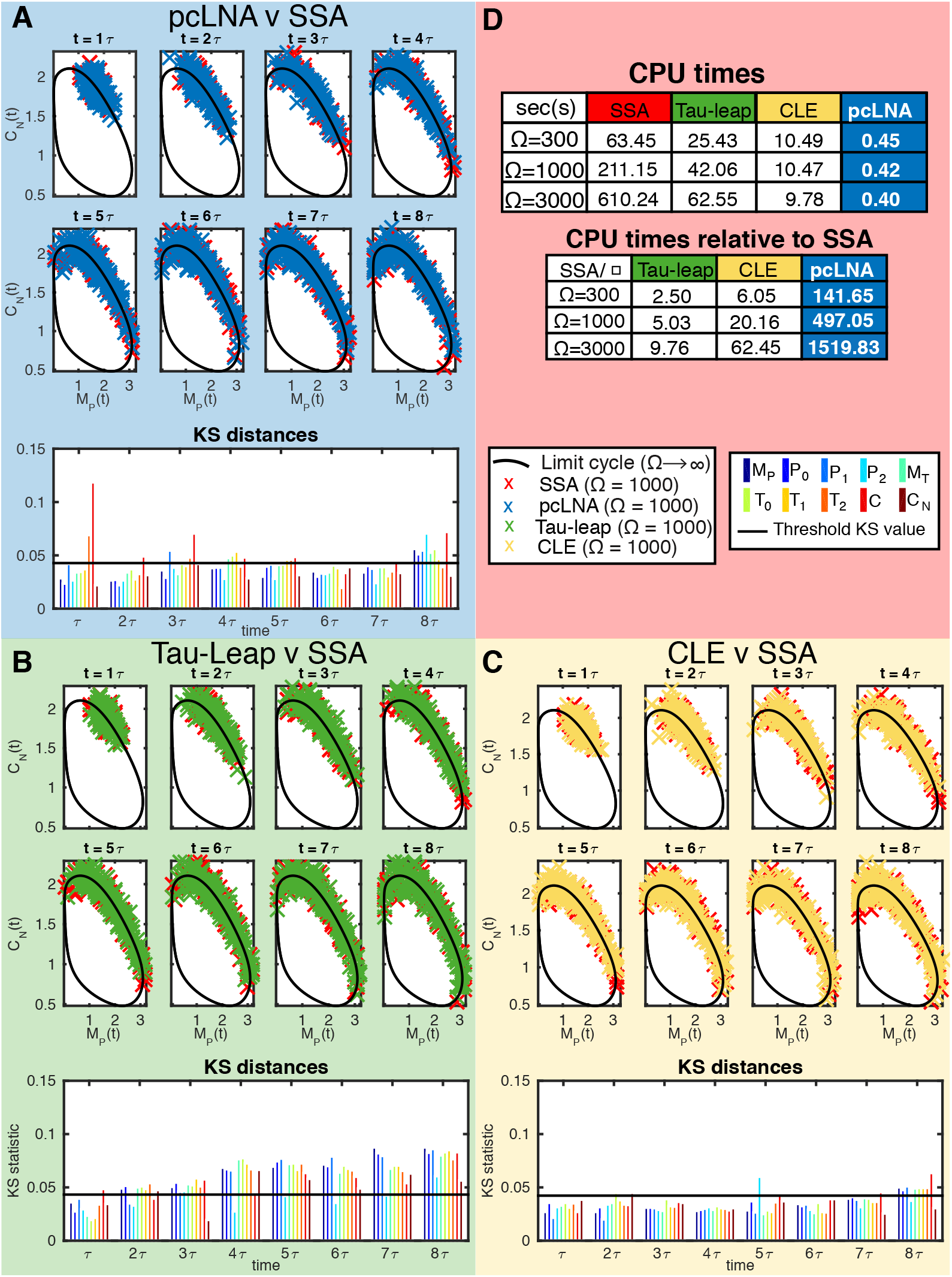
Comparison between pcLNA, tau-leap and CLE simulation algorithms for the *Drosophila* circadian clock. This figure has the same form as Fig. 9 of **I**, but (A), (B) and (C) panels contain results for Ω = 1000. As in Fig. 9 of **I**, the parameter values ∈ = 0.002 and Δ*t* = 0.002 for the tau-leap and the CLE approximation are the largest values to achieve precision similar to the pcLNA simulation, and the results displayed here are simulated using these values.

**Figure H:**
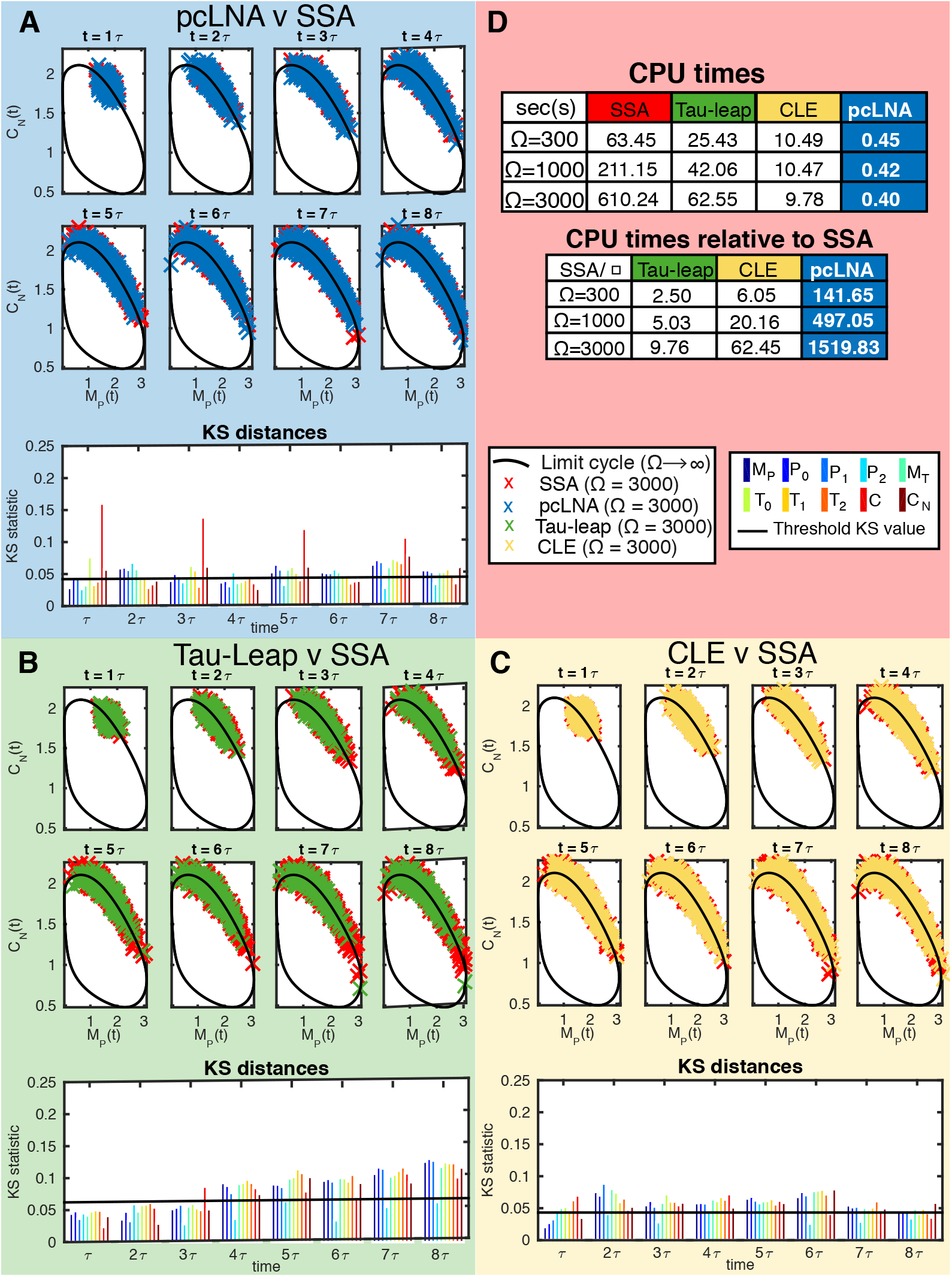
Comparison between pcLNA, tau-leap and CLE simulation algorithms for the *Drosophila* circadian clock. This figure has the same form as Fig. 9 of **I**, but (A), (B) and (C) panels contain results for Ω = 3000. As in Fig. 9 of **I**, the parameter values ∈ = 0.002 and. Δ*t* = 0.002 for the tau-leap and the CLE approximation are the largest values to achieve precision similar to the pcLNA simulation, and the results displayed here are simulated using these values.

### 7 Light entrained *D*rosophila Circadian Clock

The Light entrained *Drosophila* circadian clock system first proposed in [2] is exactly the same with the *Drosophila* circadian clock without entrainment, apart from the degradation rate, *v_dT_*, of the phosphorylated TIM protein, *T*_2_, which instead of being equal to 3.4, it is equal to 4 during the day-time (6-18h) and equal to 2 during night-time (18h-6h).

We first compare the LNA distributions at fixed times with the empirical distributions derived using SSA simulations. As we can see in Fig. I, despite that the empirical distributions of the exact stochastic model do not spread along the curved limit cycle as much as the corresponding distributions for the system without entrainment (see Fig. 3 of **I**) and that the spread of the distributions in the light entrained system appears to stabilise even after the second cycle, the standard LNA distributions still do not approximate well these distributions.

**Figure I:**
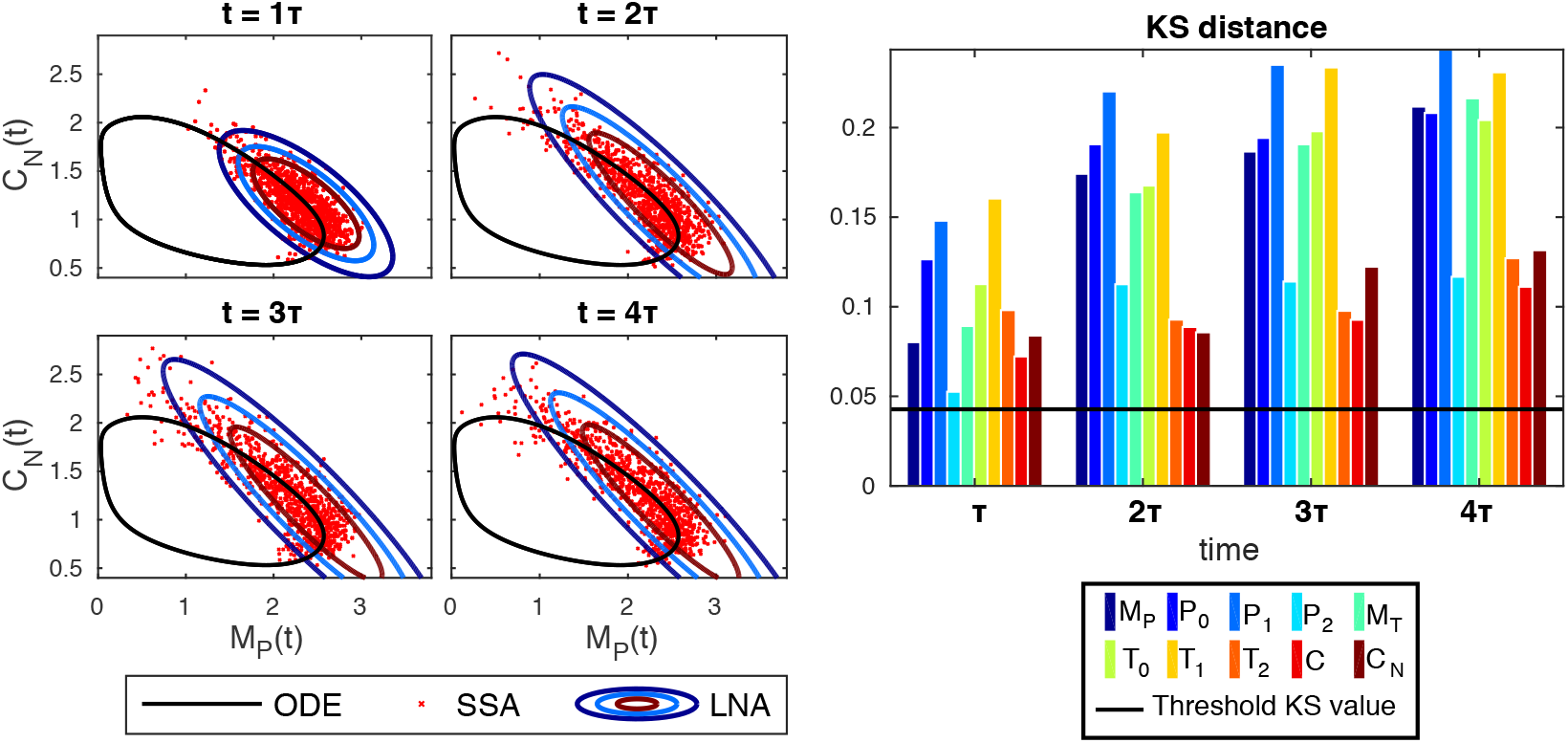
Comparison between LNA and exact simulations in the light entrained *Drosophila* circadian clock. (a) Samples (in nanomolar concentrations) obtained from the SSA simulation algorithm (red crosses) and 0.01, 0.05, 0.40 contours of the LNA probability density (black ellipsoids) at fixed times, *t* = *τ*, 2*τ*, 3*τ*, 4*τ*(*τ*: minimal period). The limit cycle ODE solution is also displayed (black solid line). (b) KS distance between the empirical distribution of SSA samples and the LNA distribution of each species (different colors, see legend) at the fixed times. The threshold level is also displayed (black solid line). The system size is Ω = 300.

On contrary, as we can see in Fig. J, the pcLNA distributions appear to approximate well the empirical transversal distributions derived under the SSA.

**Figure J:**
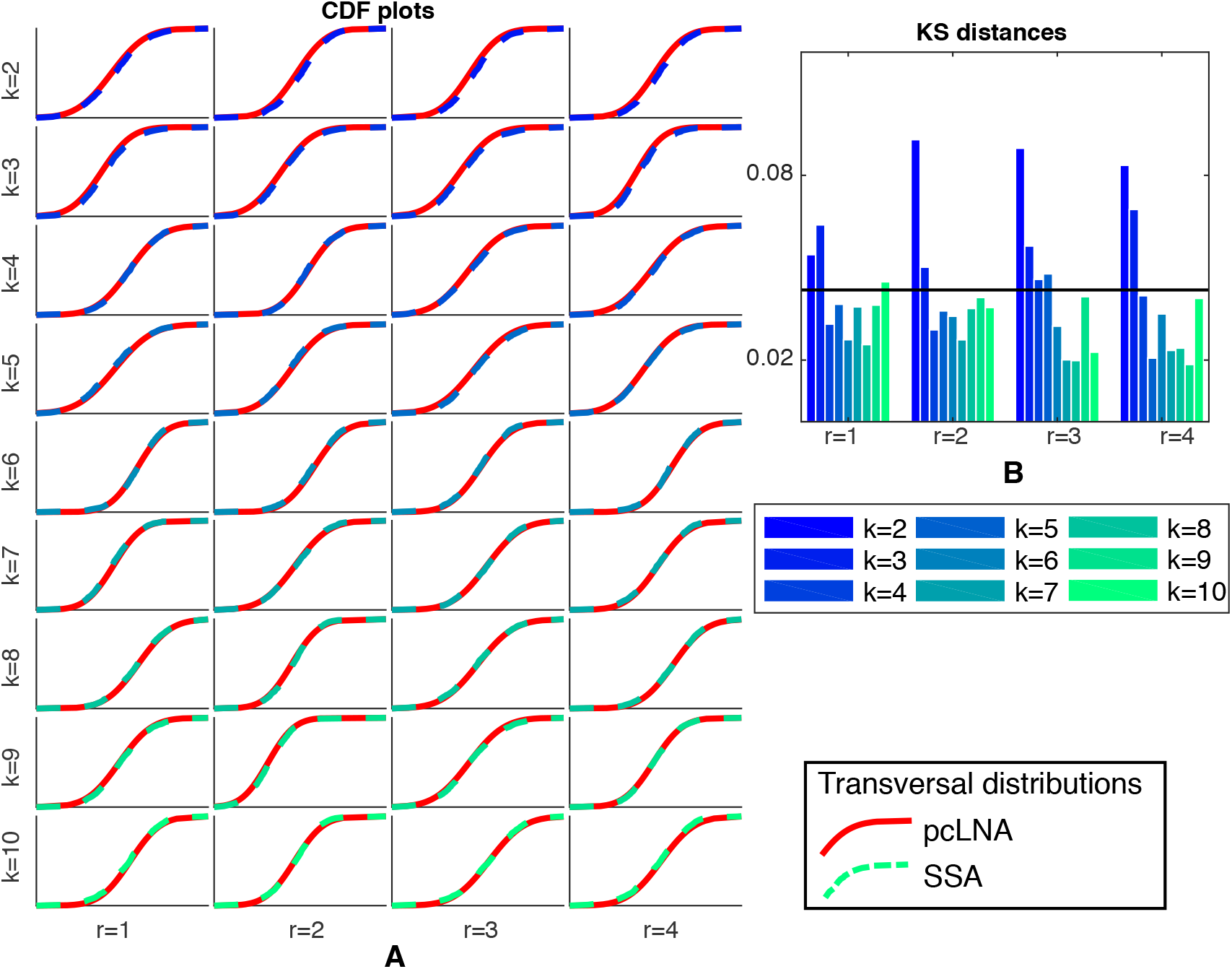
Comparison of pcLNA and exact empirical transversal distributions of the light entrained *Drosophila* circadian clock. (A) CDF plots of the transversal distributions 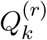 under the pcLNA (red line) and the SSA (empirical CDF, colored dashed line, see legend) in transversal coordinates *k* = 2, 3,…, 10 and round *r* = 1, 2, 3, 4. (B) KS distances between the 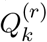distributions under the pcLNA and the SSA, *k* = 2, 3,…, 10, *r* = 1, 2, 3, 4. The system size is Ω = 300.

## Supplementary information S3 for the paper Long-time analytic approximation of large stochastic oscillators: simulation, analysis and inference

> We refer to the paper “Long-time analytic approximation of large stochastic oscillators: simulation, analysis and inference” by **I**. In this note we give further illustrations for the Brusselator and the NF-*κ*B system.

### 1 Brusselator system

The ODE system of the Brusselator is

> *Ȧ* = 1 – *A*(1 + *b – cAB*),
>
> *Ḃ* = *A*(*b – cAB*).

For the limit cycle ODE solution, we use the initial conditions *A*(*t*_0_) = 0.75, *B*(*t*_0_) = 2.00, and the parameter values *b* = 2.20, *c* = 1.00. We use the SSA to exactly simulate the system and produce *R* = 3000 samples of stochastic trajectories (see Figure A) for a time-length of 8.5 × *τ* where *τ* ≈ 6.37 is the period of the periodic solution of the ODE. The rates of the reactions used for the SSA of the Brusselator are provided in Table A.

**Table A:** Reactions of the Brusselator system and their rates.

| reaction | rate |
| --- | --- |
| $\emptyset \xrightarrow{1} A$ | $\Omega$ |
| $A \xrightarrow{1} \emptyset$ | $A$ |
| $A \xrightarrow{b} B$ | $bA$ |
| $B \xrightarrow{c} A$ | $cA^2B/\Omega^2$ |

**Figure A:**
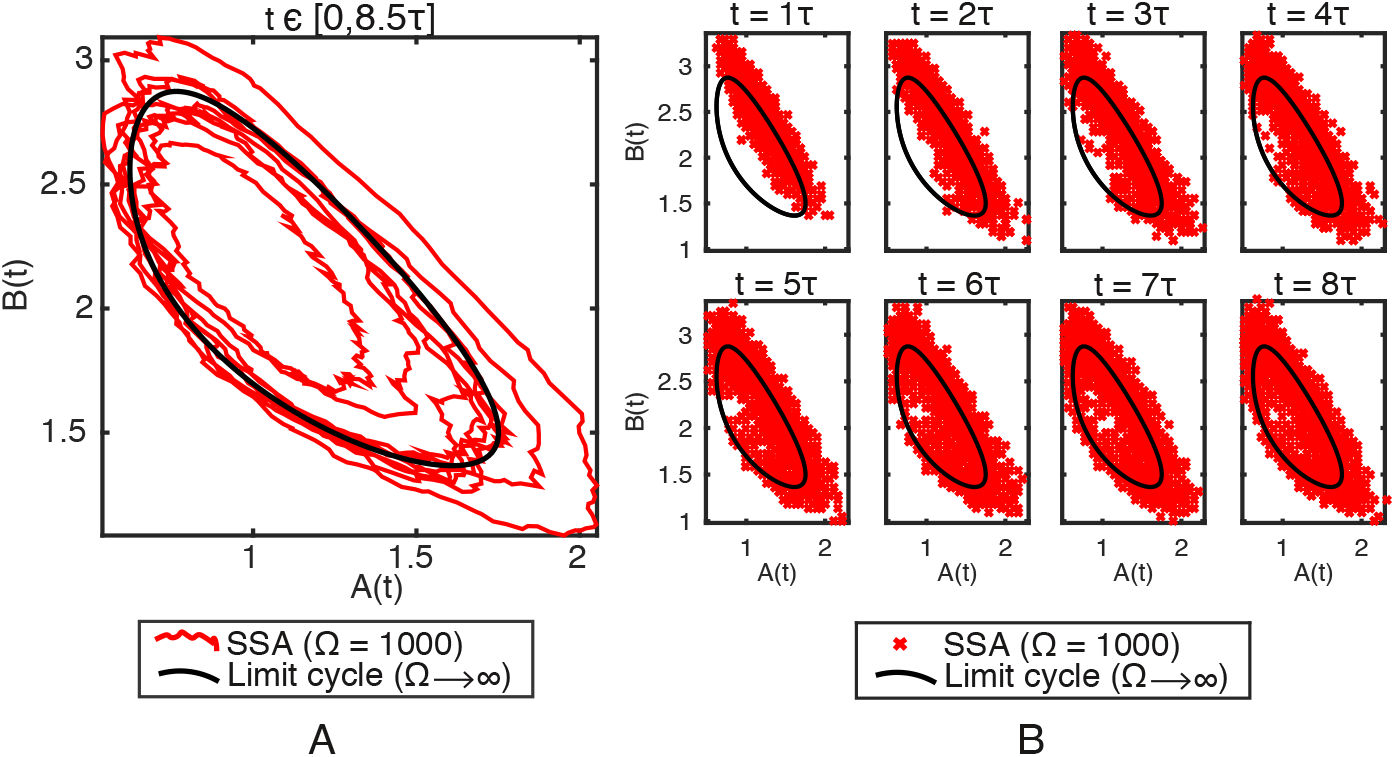
Exact stochastic simulation of the Brusselator system. (A) A stochastic trajectory of *X*(*t*) = *Y*(*t*)/Ω obtained by running the SSA over the time-interval *t* ∈ [0, 8.5*τ*] and (B) SSA samples (*R* = 3000) at times *t* = *τ*, 2*τ, …*, 8*τ*. The volume size is Ω = 1000. The black solid curve is the large volume, Ω → ∞, limit cycle solution.

**Figure B:**
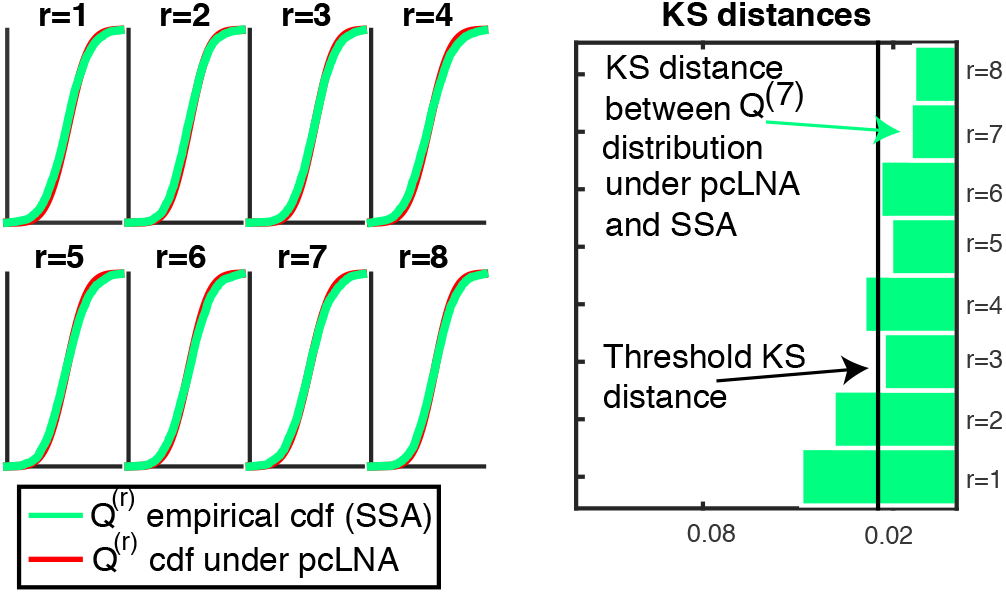
Comparison of pcLNA and exact empirical transversal distributions. (A) CDF plots of the (one-dim) transversal distributions *Q*^(r)^ under the pcLNA (red line) and the SSA (empirical CDF, crosses) in round *r* = 1, 2,…, 8. (B) KS distances between the 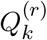distributions under pcLNA and SSA, *r* = 1, 2,…, 8.

#### 1.1 Fixed point system

For comparisons, we also consider a set of parameter values that give a limiting system (Ω → ∞) with an equilibrium fixed point instead of a limit cycle. Here, the initial conditions are *A*(*t*_0_) = 1.156, *B*(*t*_0_) = 1.461, and the parameter values *b* = 0.25, *c* = 0.25, which give an equilibrium point *A_eq_* = *B_eq_* = 1 with Jacobian matrix of the system (referred as linearisation matrix in **I**) with both eigenvalues equal to –0.5.

We use the SSA to exactly simulate this system and produce *R* = 2000 samples of stochastic trajectories (see Fig. CA) for a time-length of 700, which is somewhat larger than 100*τ*, where *τ* the period of the limit cycle ODE solution considered above. In Fig. CB we provide the empirical CDF plots of the SSA at time 637 ≈100*τ* and compare them with the LNA distributions at the same time.

**Figure C:**
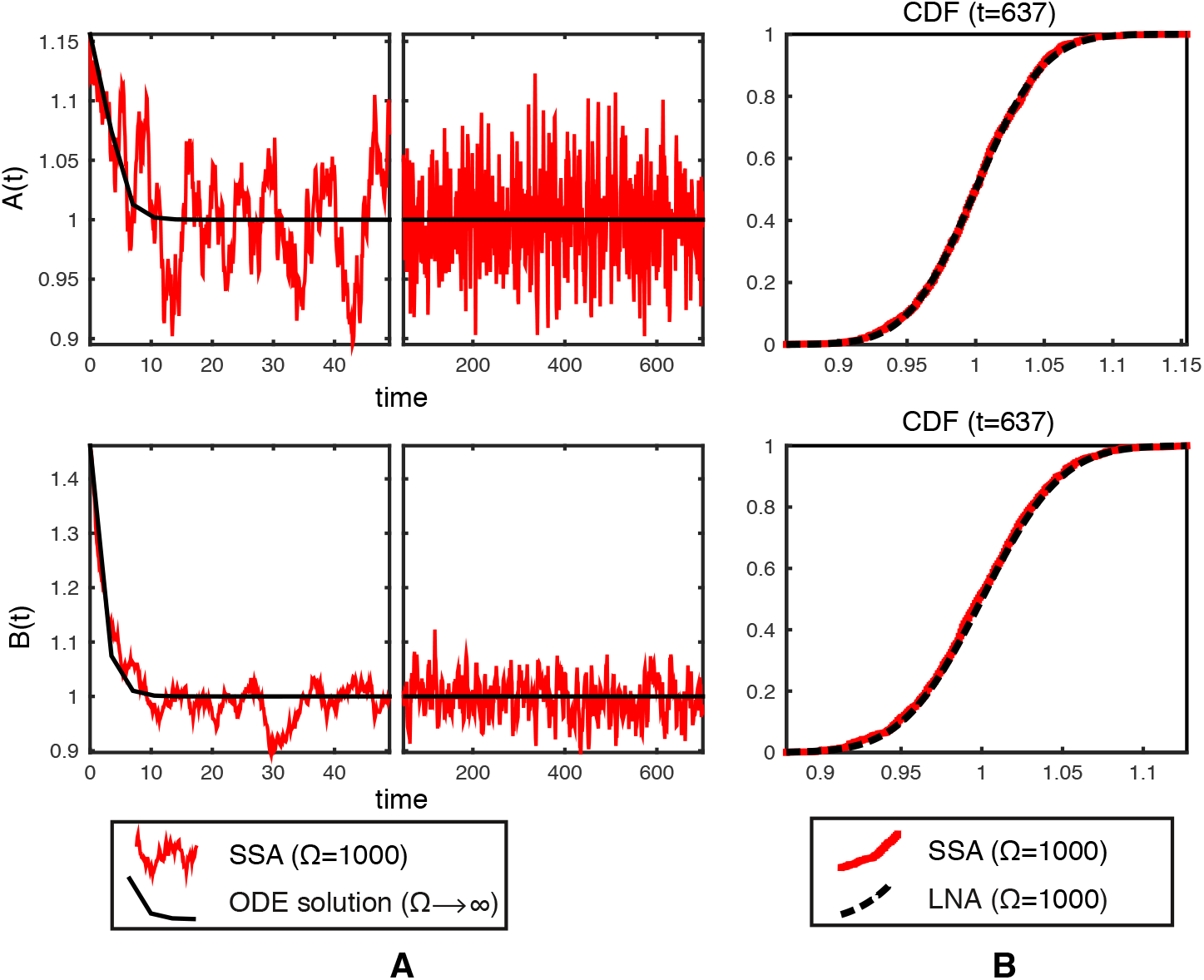
Exact stochastic simulation and comparison to the LNA for the fixed point Brusselator system. The parameter values are *b* = *c* = 0.25 and the system size Ω = 1000. (A) A stochastic trajectory of *X*(*t*) = *Y*(*t*)/Ω for the two variables *A* (top panel) and *B* (bottom panel) of the Brusselator system obtained by running the SSA over the time-interval *t* ∈ [0, 700] and its large volume, Ω → ∞, solution. (B) Empirical CDF plots of the SSA samples (*R* = 2000) and CDF plots of the LNA at time *t* = 637 for variables *A* (top panel) and *B* (bottom panel).

### 2 NF-*κ*B system

The NF-*κ*B system model used in this SI describes the oscillatory response of the system following stimulation by tumor necrosis factor alpha (TNF*α*). With no stimulation the system remains in a stable equilbrium state. Following TNF*α* stimulation that is constant in time the system responds by a transient pulse followed by relaxation to a stable limit cycle.

We first consider the case where the system starts in the stable limit cycle solution, and in the next section consider the biologically interesting situation where the signal is received while the system is inactive and the initial oscillation is transitory. The species of the system and the initial conditions (in micromolar concentrations) used to derive the limit cycle deterministic solution are provided in Table B.

**Table B:** The variables of NF-*κ* B system and the initial conditions (in concentrations) used to derive the limit cycle ODE solution.

|  | name | description | initial value |
| --- | --- | --- | --- |
| 1 | $N_c$ | Free Cytoplasmic NFkB | 0.0056 |
| 2 | $I_c$ | Free Cytoplasmic IkBa | 0.0298 |
| 3 | $NI_c$ | Cytoplasmic NFkB-IkBa | 0.0723 |
| 4 | $N_n$ | Free nuclear NFkB | 0.0011 |
| 5 | $I_n$ | Free nuclear IkBa | 0.0075 |
| 6 | $NI_n$ | Nuclear NFkB-IkBa | 0.0013 |
| 7 | $I_m$ | IkBa transcription | 0.0004 |
| 8 | $K_n$ | Kinase IKKn | 0.0020 |
| 9 | $K_a$ | Kinase IKKa | 0.0024 |
| 10 | $A_m$ | A20 transcription | 0.0001 |
| 11 | $A$ | A20 | 0.0076 |

The system model considered here is a slight modification of the system model in [1]. In particular, some variables are omitted because the sum of the concentrations of all forms of NF-*κ*B and IKK are both fixed and the level of phosphorylated IkBa is an end state that plays no further role in the model. Therefore, the redundant species IKKi, phosphorylated IkB*α* and phosphorylated NF*κ* B-I*κ*B*α* are removed to avoid rank-deficiencies in the covariance matrices of the LNA.

We also write the system in a form where concentrations are all written in terms of the same volume i.e. the total cell volume. The original system in [1] is written in cytoplasmic concentrations for all species except, *N_n_*, *I_n_* and *NI_n_* that are written in nuclear concentrations and are set to be 3.3 larger than cytoplasmic concentrations. The nuclear and cyctoplasmic concentrations can easily be recovered from our concentration if one knows the ratio of nuclear to cytoplasmic volume.

We used the same parameter values as in [1] to derive the ODE solution. These are provided in Table C.

**Table C:**
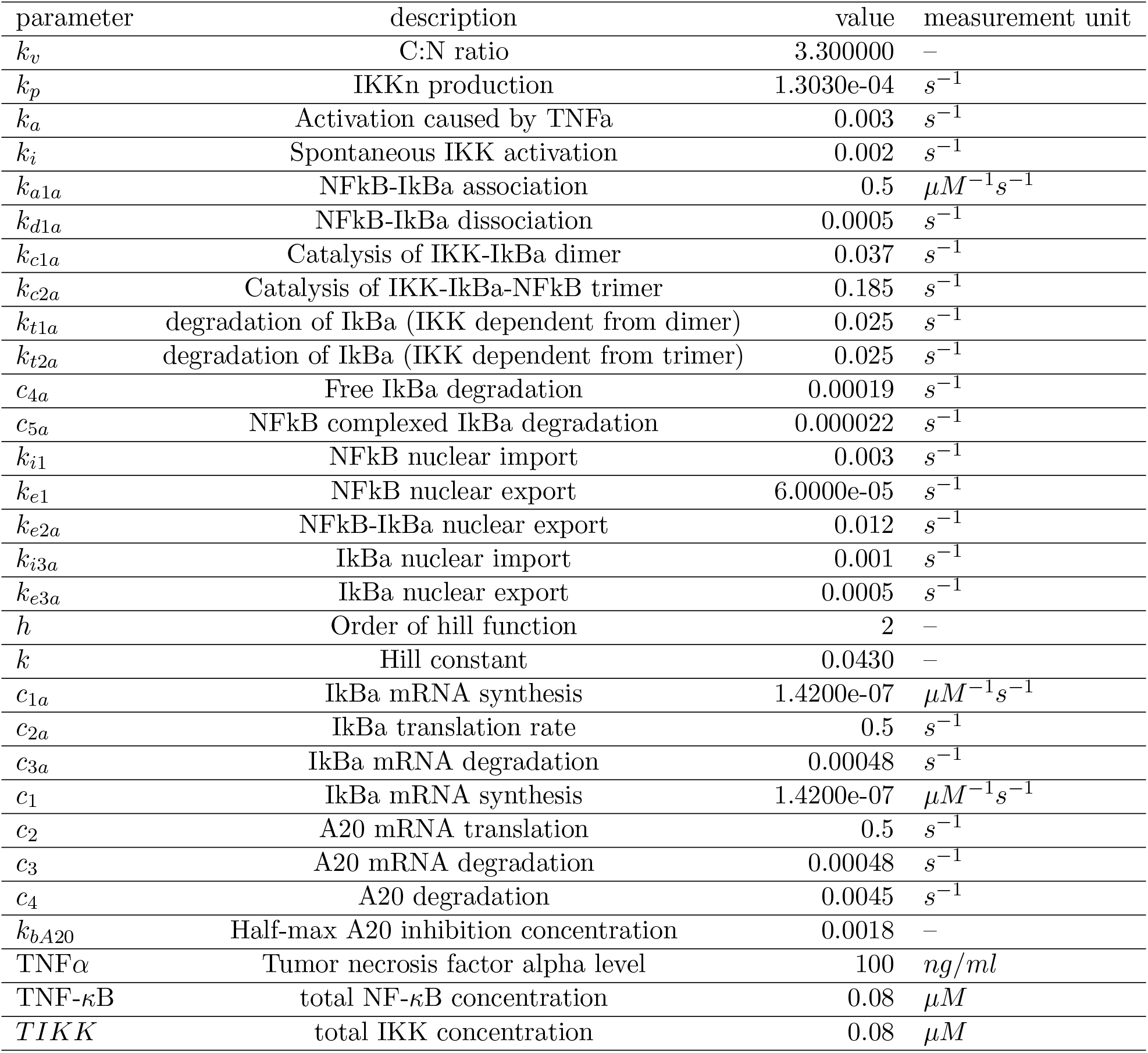
The parameters of NF-*κ*B system and the values used to derive their ODE solution.

The ODE system for the NF-B system considered in this SI is:

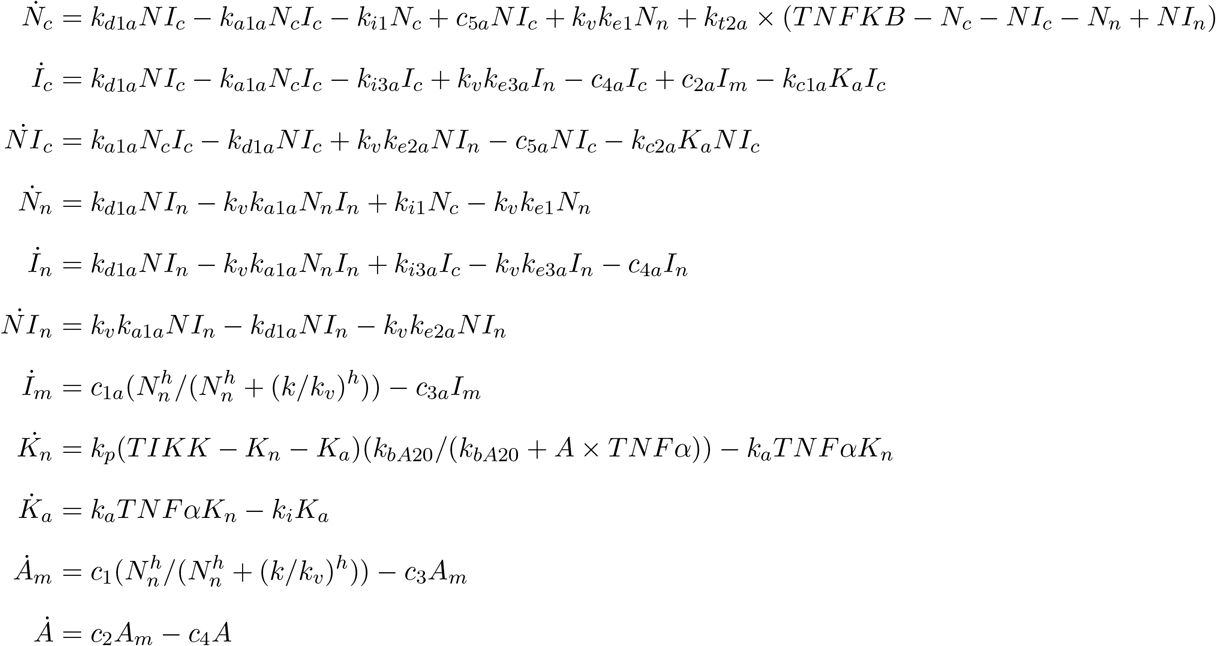

The reaction rates used for the SSA are provided in Table D. The values of the parameters are the same as in Table C.

**Table D:** Reactions of the NF-*κ*B system and their rates

| reaction | rate |
| --- | --- |
| $N_c + I_c \xrightarrow{k_{a1a}} NI_c$ | $k_{a1a} \times I_c \times N_c / \Omega$ |
| $NI_c \xrightarrow{k_{d1a}} N_c + I_c$ | $k_{d1a} \times NI_c$ |
| $N_n + I_n \xrightarrow{k_{a1a}} NI_n$ | $k_v k_{a1a} \times I_n \times N_n / \Omega$ |
| $NI_n \xrightarrow{k_{d1a}} N_n + I_n$ | $k_{d1an} \times NI_n$ |
| $K_a + I_c \xrightarrow{k_{c1a}} K_a + I_p$ | $k_{c1a} \times K_a \times I_c / \Omega$ |
| $K_a + NI_c \xrightarrow{k_{c2a}} K_a + NI_p$ | $k_{c2a} \times K_a \times NI_c / \Omega$ |
| $NI_p \xrightarrow{k_{t2a}} N_c$ | $k_{t2a} \times NI_p$ |
| $N_c \xrightarrow{k_{i1}} N_n$ | $k_{i1} \times N_c$ |
| $N_n \xrightarrow{k_{e1}} N_c$ | $k_{e1} \times k_v \times N_n$ |
| $NI_n \xrightarrow{k_{e2a}} NI_c$ | $k_{e2a} \times k_v \times NI_n$ |
| $I_c \xrightarrow{k_{i3a}} I_n$ | $k_{i3a} \times I_c$ |
| $I_n \xrightarrow{k_{e3a}} I_c$ | $k_{e3a} \times k_v \times I_c$ |
| $\emptyset \xrightarrow{H_I} I_m$ | $(c_{1a}\Omega) \frac{N_n^h}{N_n^h + (k\Omega/k_v)^h}$ |
| $I_m \xrightarrow{c_{2a}} I_m + I_c$ | $c_{2a} \times I_m$ |
| $I_m \xrightarrow{c_{3a}} \emptyset$ | $c_{3a} \times I_m$ |
| $I_c \xrightarrow{c_{4a}} \emptyset$ | $c_{4a} \times I_c$ |
| $I_n \xrightarrow{c_{4a}} \emptyset$ | $c_{4a} \times I_n$ |
| $NI_c \xrightarrow{c_{5a}} N_c$ | $c_{5a} \times NI_c$ |
| $\emptyset \xrightarrow{H_A} A_m$ | $(c_1\Omega) \frac{N_n^h}{N_n^h + (k\Omega/k_v)^h}$ |
| $A_m \xrightarrow{c_2} A_m + A$ | $c_2 \times A_m$ |
| $v A_m \xrightarrow{c_3} \emptyset$ | $c_3 \times A_m$ |
| $A \xrightarrow{c_4} \emptyset$ | $c_4 \times A$ |
| $K_i \xrightarrow{M_A} K_n$ | $k_p(TIKK - K_n - K_a) \frac{k_{bA20} \times \Omega}{(k_{bA20}\Omega) + A \times TNF\alpha}$ |
| $K_n \xrightarrow{TNF\alpha \times k_a} K_a$ | $TNF\alpha \times k_a \times K_n$ |
| $K_a \xrightarrow{k_i} K_i$ | $k_i \times K_a$ |

**Figure D:**
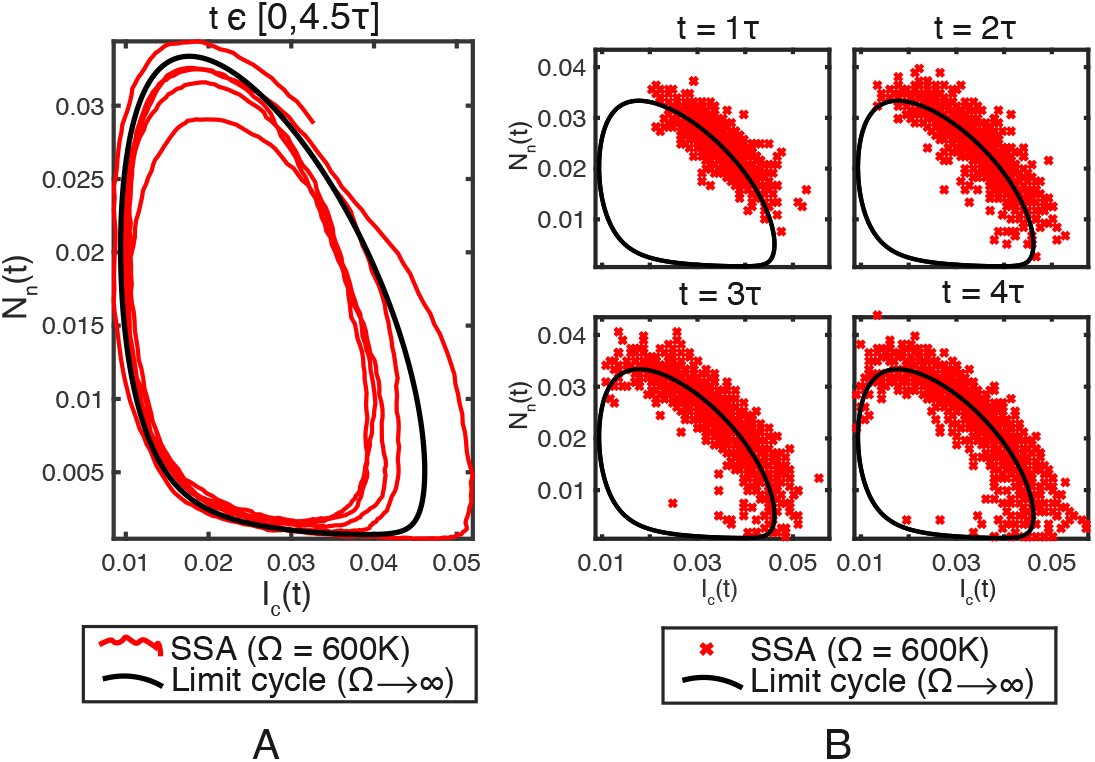
Exact stochastic simulation of the NF-*κ*B system. (A) A stochastic trajectory (in concentrations) obtained by running the SSA over the time-interval *t* ∈ [0, 4.5*τ*] and (B) SSA samples (*R* = 3000) at times *t* = *τ*, 2*τ*, 3*τ*, 4*τ*. Two (out of 11) of the species are displayed (I*κ* B*α I_C_* (x-axis) and nuclear NF-*κ*B *N_n_* (y-axis)). The volume size is Ω = 0.6*M*. This is substantially smaller than the system size used in [1] (Ω = 1.25*M*) to provide higher noise levels. The black solid curve is the large volume, Ω → ∞, limit cycle solution.

**Figure E:**
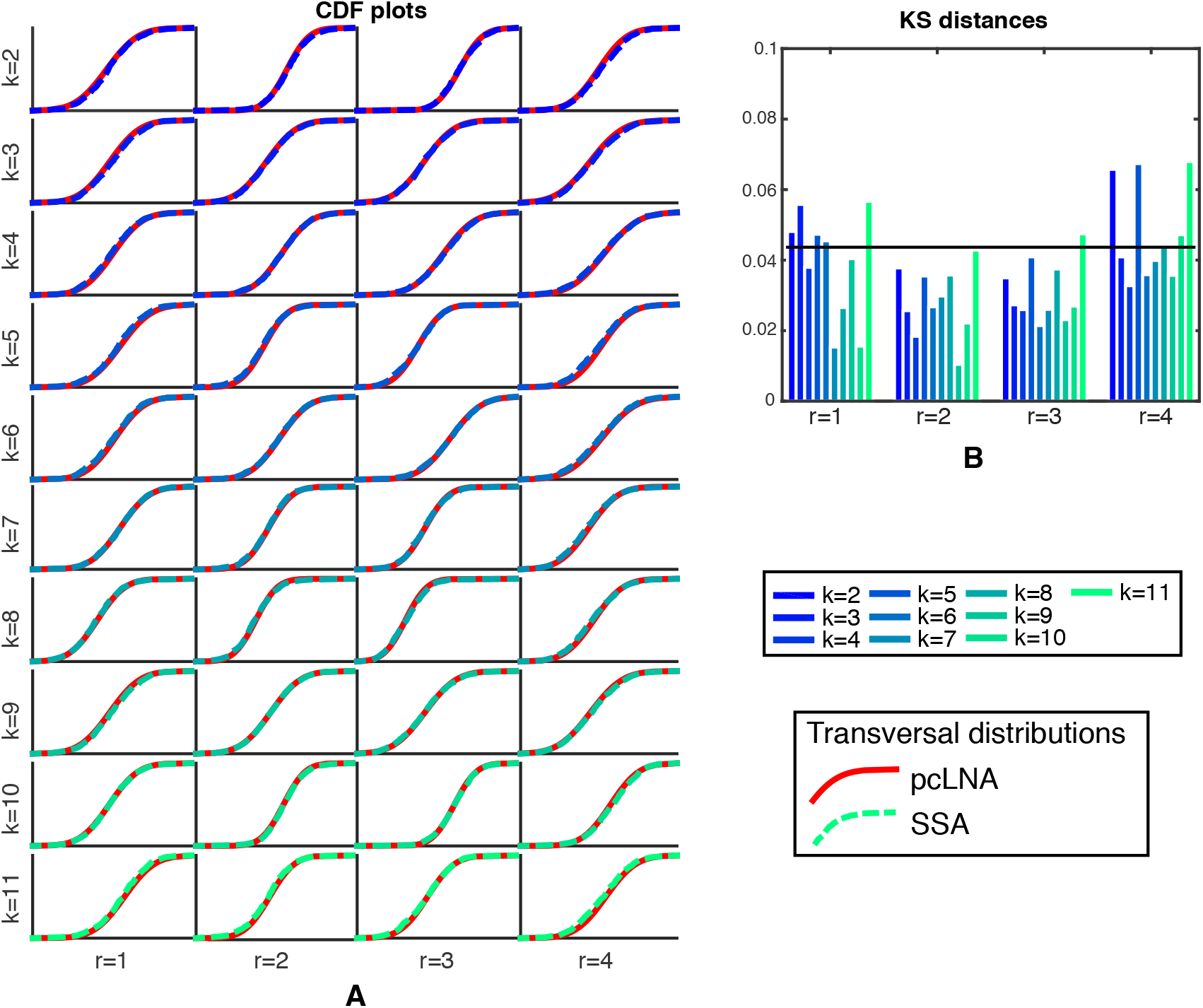
Comparison of pcLNA and exact transversal distributions. (A) CDF plots of the transversal distributions 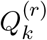 under the pcLNA (red line) and the SSA (empirical CDF, colored dashed line, see legend) in transversal coordinates *k* = 2, 3,…, 11 and round *r* = 1, 2, 3, 4. (B) KS distances between the 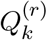distributions under pcLNA and SSA, *k* = 2, 3,…, 11, *r* = 1, 2, 3, 4. The system size is Ω = 0.6*M*.

#### 2.1 The NF-*κ*B response to TNF*α* activation

Here we study the response of the *NF-κB* system to a continuous TNF*α* signal received while the system being in the equilibrium fixed point state. This equilibrium state which is given in Table E is therefore the initial conditions used to derive the deterministic solution provided in Fig. F.

**Table E:** The initial conditions (in concentrations) used to derive the ODE solution in Fig 6.

|  | name | initial value |
| --- | --- | --- |
| 1 | $N_c$ | 0.0036 |
| 2 | $I_c$ | 0.0160 |
| 3 | $NI_c$ | 0.0719 |
| 4 | $N_n$ | 0.0042 |
| 5 | $I_n$ | 0.0013 |
| 6 | $NI_n$ | 0.0003 |
| 7 | $I_m$ | 0.0000 |
| 8 | $K_n$ | 0.0800 |
| 9 | $K_a$ | 0 |
| 10 | $A_m$ | 0.0000 |
| 11 | $A$ | 0.0014 |

**Figure F:**
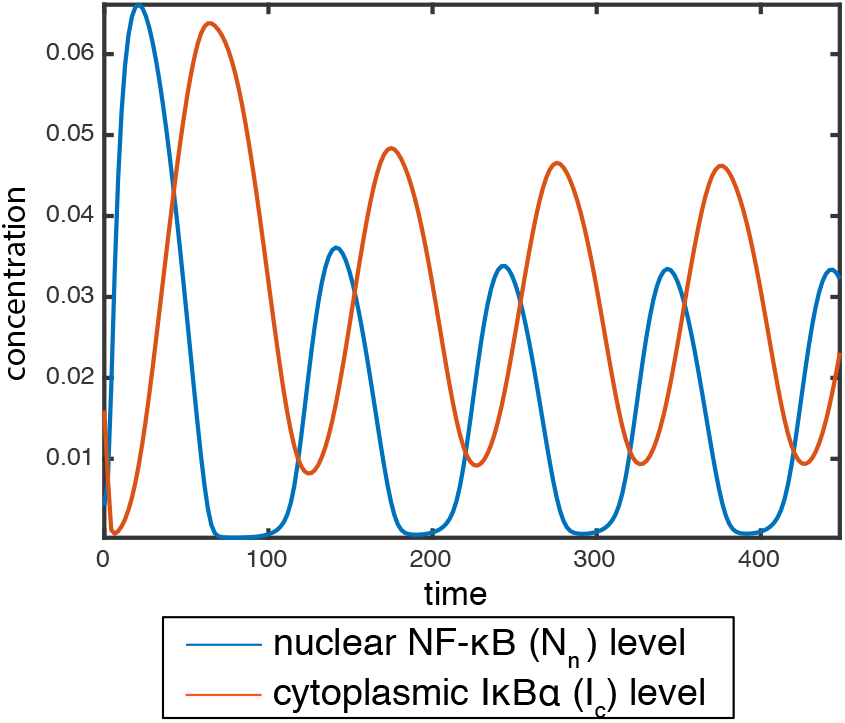
The limiting (Ω → ∞) deterministic solution of the NF-*κ* B system. The initial conditions are provided in Table E. Two species of the system are displayed.

**Figure G:**
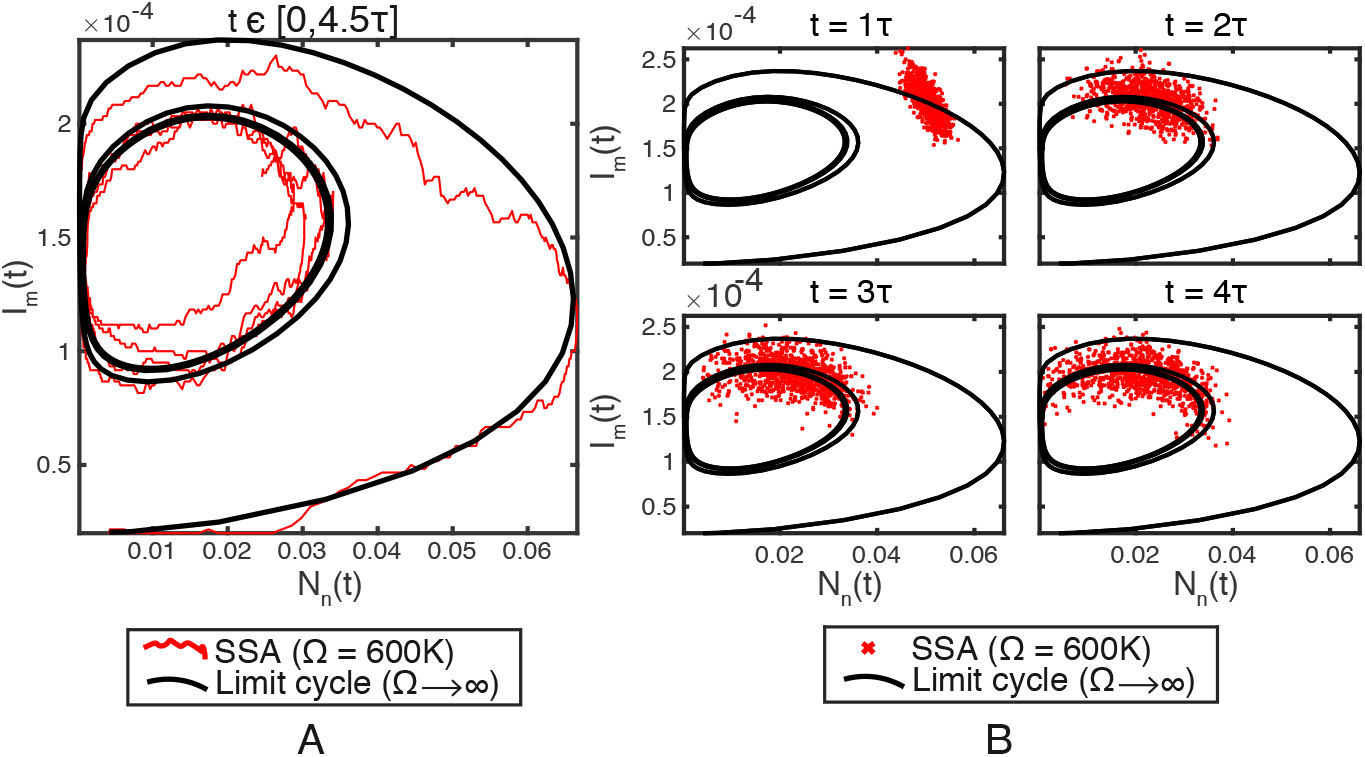
Exact stochastic simulation of the NF-*κ*B system. (A) A stochastic trajectory (in concentrations) obtained by running the SSA over the time-interval *t* ∈ [0, 4.5*τ*] and (B) SSA samples (*R* = 3000) at times *t* = *τ*, 2*τ*, 3*τ*, 4*τ*. Two (out of 11) of the species are displayed (nuclear NF-*κ*B (x-axis) and I*κ*B*α* mRNA (y-axis)). The volume size is Ω = 0.6*M*. The black solid curve is the large volume, Ω → ∞, limit cycle solution.

**Figure H:**
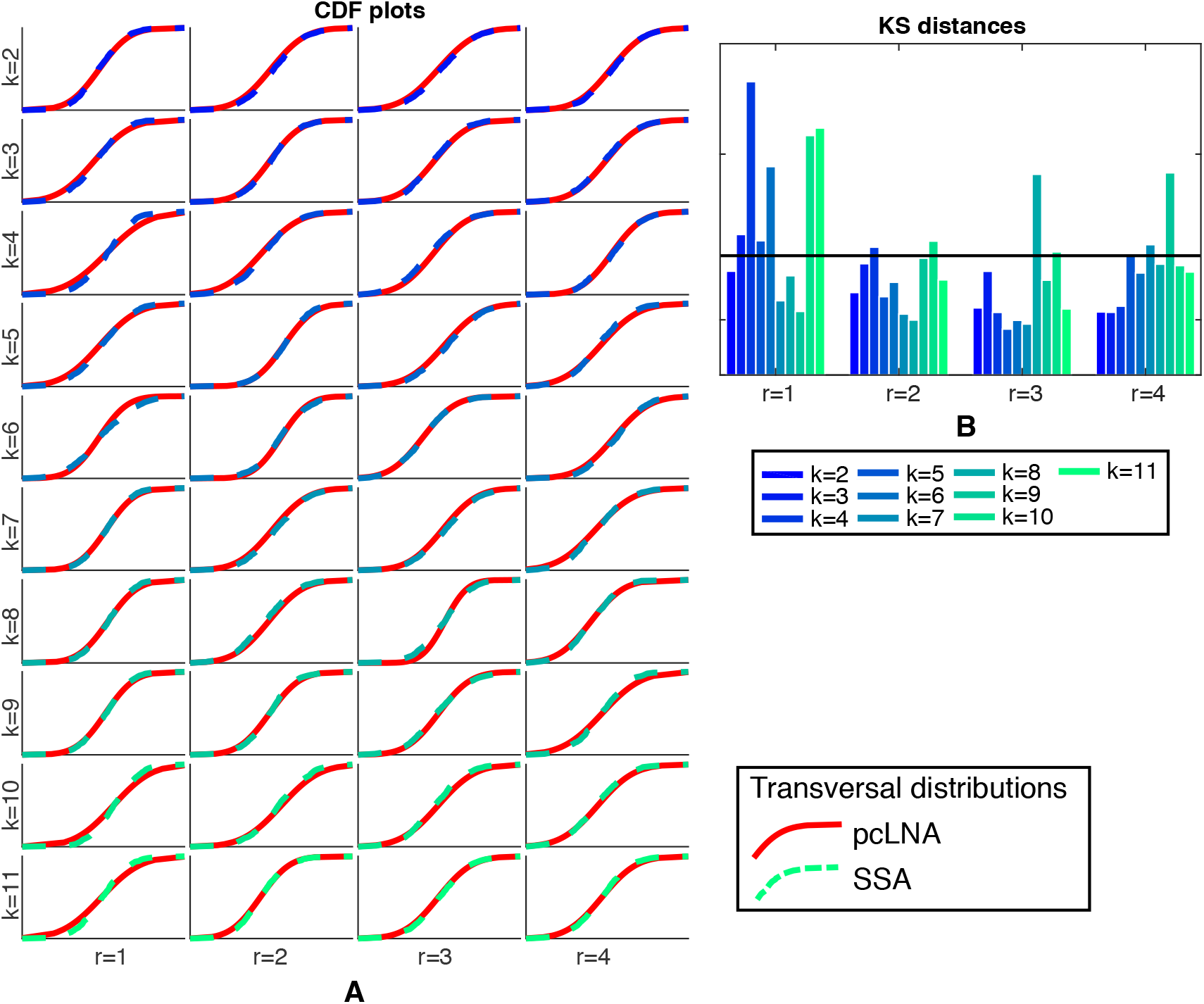
Comparison of pcLNA and exact transversal distributions. (A) CDF plots of the transversal distributions 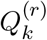under the pcLNA (red line) and the SSA (empirical CDF, colored dashed line, see legend) in transversal coordinates *k* = 2, 3,…, 11 and round *r* = 1, 2, 3, 4. (B) KS distances between the 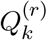distributions under pcLNA and SSA, *k* = 2, 3,…, 11, *r* = 1, 2, 3, 4. The system size is Ω = 0.6*M*.

